# Resolving malignant cell heterogeneity from bulk tumor RNA-seq data with CDState

**DOI:** 10.1101/2025.03.01.641017

**Authors:** Agnieszka Kraft, Josephine Yates, Florian Barkmann, Valentina Boeva

**Affiliations:** Institute for Machine Learning, Department of Computer Science, ETH Zurich, Switzerland; Department of Thoracic Surgery, University Hospital Zurich, Switzerland; Swiss Institute for Bioinformatics, Switzerland; ETH AI Center, ETH Zurich, Switzerland

## Abstract

Intratumor transcriptional heterogeneity (ITTH), defined by the coexistence of diverse cell states within one tumor, complicates cancer treatment by contributing to variable therapeutic responses. Although single-cell RNA sequencing can resolve this complexity, its cost and technical demands limit its large-scale use. Bulk RNA-seq data provide a scalable alternative, but most deconvolution methods depend on predefined references, restricting their ability to detect novel malignant states. Unsupervised approaches avoid these constraints but are not tailored to capture heterogeneity within the malignant compartment. To address these limitations, we introduce CDState, an unsupervised method for inferring malignant cell subpopulations from bulk RNA-seq data. CDState utilizes non-negative matrix factorization improved with sum-to-one constraint and a cosine similarity-based optimization to deconvolve bulk gene expression into distinct cell state profiles. We demonstrate robustness of CDState on bulkified single-cell RNA-seq datasets from five cancer types, showing that it outperforms existing unsupervised deconvolution methods in the estimation of both cell state proportions and gene expression profiles. Applied to 33 cancer types from The Cancer Genome Atlas, CDState reveals recurrent gene programs, including epithelial-mesenchymal transition, MYC targets, and oxidative phosphorylation, as major contributors to malignant cell ITTH. We further link malignant states to patient clinical features, identifying states associated with poor prognosis. We propose an intratumor heterogeneity index and show its association with patient survival, clinical characteristics, and therapeutic response. Finally, we identify mutations and copy number alterations in genes such as *TP53*, *KRAS*, *PIK3CA, SOX2,* and *SATB1* as potential genetic drivers of malignant cell ITTH across cancer types.

## Introduction

Single-cell sequencing technology has significantly advanced our understanding of intratumor transcriptional heterogeneity (ITTH), revealing the diverse gene expression programs within tumor-resident cells. This heterogeneity has been shown to be implicated in cancer progression, metastasis, and therapy resistance [1–3]. While pioneering studies focused specifically on the tumor microenvironment (TME) composition, recently more emphasis has been put on the heterogeneity within the malignant compartment [4–9]. For instance, single-cell transcriptomic analyses of glioblastoma, melanoma, head and neck squamous cell carcinoma, esophageal adenocarcinoma, and colorectal cancer have identified distinct malignant cell states, characterized by differences in differentiation status, stress response, and plasticity [4,5,7,10,11]. These states are closely intertwined with TME composition, influencing processes such as angiogenesis [12] and immune evasion [7,13,14]. Moreover, pan-cancer studies have suggested that recurrent malignant cell states are shared across multiple tumor types, implying that common oncogenic programs may drive ITTH across cancers [15,16]. These findings raise the possibility that targeting specific malignant states could offer a broadly applicable therapeutic strategy.

Although highly informative, single-cell sequencing remains costly and technically demanding, making it impractical for comprehensive screening of large-scale tumor datasets and for application in clinical settings [17]. Conversely, bulk RNA sequencing (RNA-seq) data from multiple databases, spanning diverse samples and cancer types, provide a promising alternative for robust characterization of tumor composition. Over the past decade, numerous computational deconvolution methods have been developed to infer cell type proportions from bulk RNA-seq data [18–22]. However, most of these methods rely on reference transcriptomic data for cell types present within the bulk mixture, requiring a priori knowledge of the sample composition and often assuming a stable distribution of cell types across samples [23]. More critically, these reference-based methods are unable to discover malignant cell states absent in the reference data, limiting their ability to accurately resolve tumor-specific transcriptional heterogeneity. Therefore, in the absence of appropriate reference data, unsupervised methods that simultaneously recover cell-type and cell-state expression profiles and relative abundances from bulk samples offer a more effective strategy. Yet most unsupervised methods focus on general cell type identification rather than resolving malignant cell states.

To address these limitations, we introduce CDState, an unsupervised deconvolution framework based on non-negative matrix factorization (NMF). Unlike conventional deconvolution approaches, CDState is designed to identify and characterize malignant cell states in bulk RNA-seq data without requiring predefined marker genes or reference datasets. To ensure biologically meaningful deconvolution, CDState integrates sum-to-one constraints on the estimated proportions and a cosine similarity-based optimization term, which encourages the disentanglement of cell states with similar gene expression profiles.

We validate CDState using bulkified single-cell RNA-seq (scRNA-seq) datasets from glioblastoma [5], lung adenocarcinoma [24], breast cancer [25], ovarian high grade serous carcinoma [26], and squamous cell carcinoma [27], showing that it outperforms existing unsupervised deconvolution methods [28–33] in both cell state proportions and gene expression estimates across diverse tumor contexts. Further, we apply CDState to 33 cancer types from The Cancer Genome Atlas (TCGA) to provide a comprehensive characterization of malignant ITTH across cancers. We identify recurrent gene programs contributing to ITTH and show the association between ITTH and clinical outcomes and treatment response across cancer types. Finally, we identify potential genetic drivers of transcriptional heterogeneity, which could help better understand mechanisms shaping intratumor heterogeneity. Together, our results highlight the utility of CDState for robust reference-free characterization of malignant cell states from bulk RNA-seq data, offering a framework to investigate mechanisms and consequences of ITTH.

## Results

### Overview of CDState

CDState is a deconvolution framework based on the NMF algorithm augmented with additional constraints to enhance both accuracy and interpretability (Figure 1A–B). The method takes as input normalized bulk RNA-seq data along with tumor purity estimates, and outputs both gene expression profiles of underlying sources (cell types or cell states) and their relative proportions across samples.

**Figure 1.**
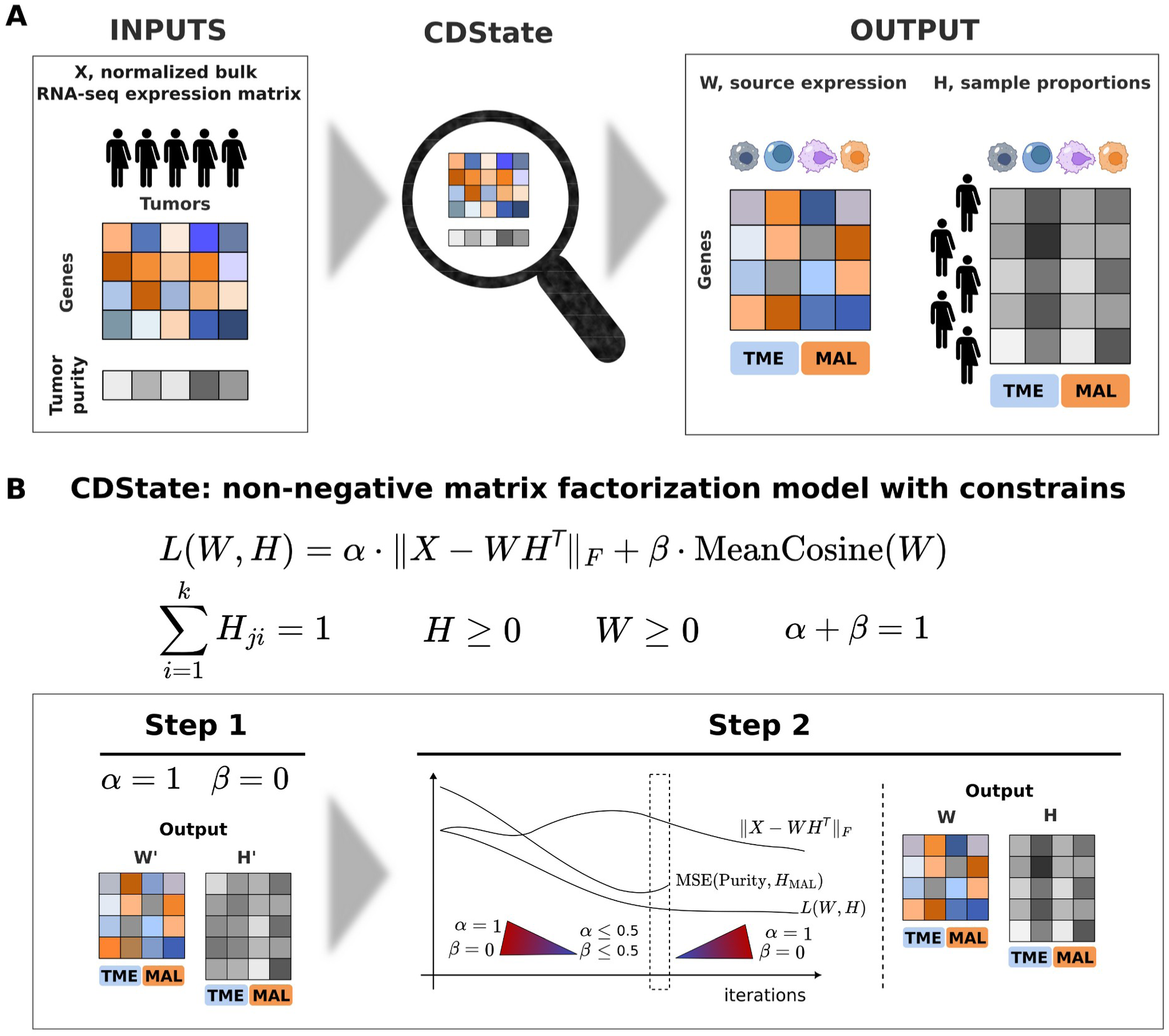
CDState overview. **A.** CDState takes as input a normalized bulk RNA-seq expression matrix and tumor purity estimates. It outputs matrices of gene expression of sources (W) and source proportions in samples (H), with source identities assigned to tumor microenvironment (TME) or malignant (MAL) states. **B.** CDState uses a constrained non-negative matrix factorization (NMF) model that balances reconstruction accuracy and source separation. The algorithm proceeds in two steps: initialization with α = 1, β = 0, followed by an annealing optimization to minimize reconstruction error and improve proportions fit to purity estimates under the constraint α + β = 1.

Compared to other NMF-based solutions, which rescale weights after deconvolution, CDState imposes sum-to-one constraints on estimated weights into the optimization process, making them more biologically interpretable, and mitigating biases in gene expression estimates arising from rescaling the matrices (Figure 1B, Methods). In addition, the objective function includes a term that penalizes high cosine similarity between deconvolved source gene expression profiles. This encourages greater distinction between transcriptionally similar states, such as fibroblasts and malignant cells with mesenchymal properties.

CDState consists of two steps guided by parameters α and β summing to 1, which control the balance between reconstruction error and source transcriptional profile separation during the optimization (Figure 1B, Methods). In the first step, CDState minimizes reconstruction error (β = 0). In the second step, β is gradually increased, giving more importance to minimizing the cosine similarity of source gene expression profiles, while temporarily allowing for higher reconstruction error. This procedure has been inspired by simulated annealing, a technique commonly used in algorithm optimization to expand the search space and avoid convergence to local minima [34]. The selection of values for α and β is guided by monitoring the mean squared error (MSE) between the estimated and true purity of deconvolved samples (Methods). Once the MSE stops decreasing, β is gradually reduced back to zero for a final optimization step that minimizes reconstruction error (Figure 1B, Suppl. Fig. S1 A-D, Methods).

### CDState outperforms existing methods in cell-type deconvolution

We first validated CDState on bulkified scRNA-seq data from lung adenocarcinoma (*Kim et al.* [24]), by sampling cells to simulate scenarios that reflect common challenges in bulk deconvolution, such as the absence of pure samples and imbalanced cell-type frequencies. Specifically, we created four synthetic bulk RNA-seq datasets (mixtures), each composed of different proportions of three cell types: malignant cells, B cells, and fibroblasts. For each mixture, tumor samples were generated by aggregating scRNA-seq read counts, with cell-type proportions sampled from Dirichlet distributions with varying concentration parameters (Figure 2, Methods). This setup allowed us to evaluate CDState’s performance under different conditions, including the presence of pure samples for some cell types (MixA and MixC), the absence of pure samples (MixB and MixD), and the inclusion of rare cell types (MixD).

**Figure 2.**
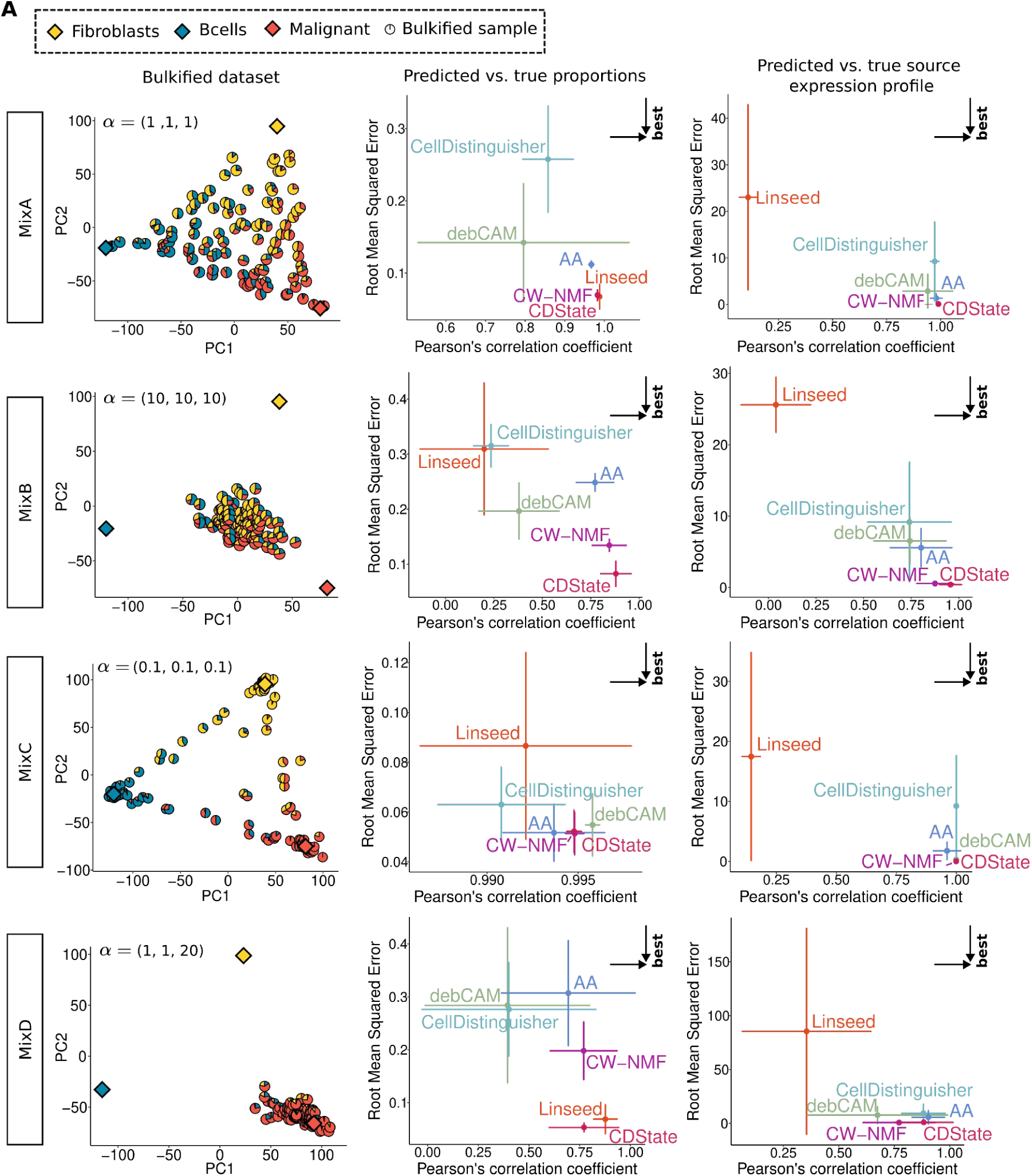
Validation of CDState for cell type deconvolution. Performance on four synthetic mixtures (in rows), comparing CDState with four external methods and with the first step of CDState (CW-NMF). Mixtures were generated by sampling cell type proportions from four Dirichlet distributions with varying alpha parameters. These proportions were used to aggregate read counts from malignant cells, B cells, and fibroblasts from the *Kim et al.* dataset [24] (Methods). Bulkified samples were visualized in the PCA space projected on the ground truth source gene expression. Pie chart coloring corresponds to the proportion of the respective cell type in each bulkified sample (left column). Deconvolution performance was compared using Root Mean Squared Error and Pearson’s correlation coefficient on both inferred proportions (middle column) and source gene expression profiles (right column).

We compared CDState against four state-of-the-art unsupervised deconvolution methods: Archetype Analysis (AA) [32,33], debCAM [30,31], Linseed [29], and CellDistinguisher [28]. These methods were selected based on a recent benchmark highlighting them as top-performing deconvolution tools [35], while AA was included as it was previously used to study molecular phenotypes in cancer [36]. To account for variability in the inferred solutions, all methods (except for the deterministic CellDistinguisher) were run 20 times with different random seeds. To ensure a fair comparison, we set all methods to deconvolve the data into three components, matching the known number of cell types in the mixtures. Additionally, we included results from CDState’s first optimization step (constrained-weight-NMF, CW-NMF) to verify the impact of the cosine similarity term.

CDState consistently achieved the highest correlation and lowest root mean squared error (RMSE) with both ground truth source profiles and proportions (Figure 2, Suppl. Figure S2). Notably, it showed the smallest variability in performance metrics across multiple runs, indicating robust and stable results. Furthermore, the cosine similarity-based optimization improved CDState performance by reducing RMSE and improving correlation with true source profiles compared to the CW-NMF results (Figure 2, Suppl. Figure S2). This improvement was especially evident in more complex settings, such as MixB and MixD, where no pure samples were available. As expected, we also observed a decrease in cosine similarity between the inferred and ground truth sources, further validating the role of the cosine similarity term (Suppl. Figure S3).

Next, we investigated the importance of the annealing procedure, which balances reconstruction accuracy against source separation by gradually updating their relative weights, α and β, during optimization. Specifically, we evaluated CDState performance under four scenarios (Suppl. Fig. S4): (1) the first-step solution (β=0, ‘Initial’), (2) optimization with fixed positive β>0 (‘Fixed beta’), (3) the second-step solution without the final cooldown (β gradually increased from β=0, ‘No cooldown’), and (4) the final solution (β gradually increased from 0 and then decreased to β=0, ‘Final’). We observed consistently accurate results for MixA and MixC in modes 1, 3, and 4, with worse relative RMSE (rRMSE) for both proportions and sources in mode 2, *i.e*., in solutions with fixed beta (Suppl. Figure S4). In the case of more complex mixtures, MixB and MixD, modes 3 and 4, “No cooldown” and “Final”, showed the best performance. Of note, particularly in MixB and MixD, we observed improved performance compared to the “Initial” solution, which further supports the use of our cosine-based optimization. While the “Fixed beta” approach already offers improvement compared to the “Initial” solution, it requires an appropriate selection of β by the user. Meanwhile, our proposed annealing procedure avoids this issue by selecting the optimal parameters during the optimization and yields the best results in the most challenging bulk datasets.

Given that CDState requires ground-truth purity information for optimization, we evaluated its performance in the case of noisy purity estimates. For the four synthetic mixtures, we perturbed the true purity values by adding Gaussian noise with varying standard deviations (Methods). CDState maintained high accuracy at σ=0.1 and remained nearly as accurate at σ=0.25, indicating robustness to moderate noise in purity estimates (Suppl. Fig. S5).

Finally, to better understand how CDState compares with supervised deconvolution methods, we evaluated its performance in proportion estimation with results from InstaPrism [37], a fast implementation of the state-of-the-art BayesPrism method [22][38]. CDState achieved lower rRMSE in 3/4 mixtures and higher correlation in 2/4 mixtures (Suppl. Figure S6), demonstrating performance comparable to supervised deconvolution models that leverage matched single-cell references (Methods).

### CDState outperforms existing methods in malignant cell-state deconvolution

To evaluate CDState’s performance in inferring malignant cell states and corresponding cell proportions from bulk RNA-seq data, we applied our deconvolution method along with the four existing approaches to five bulkified scRNA-seq datasets, including malignant and non-malignant TME cells, from breast cancer [25], ovarian cancer [26], glioblastoma [5], lung cancer [24], and squamous cell carcinoma [27] (Methods). In this analysis, we selected the number of components for each method so that the number of components assigned to the malignant compartment matches the expected number of malignant cell states reported in the original studies, without constraining the number of TME components (Methods, Suppl. Table S1-S2).

We first assessed the ability of each method to separate the signal originating from malignant cells from that of non-malignant components of the TME, using tumor purity estimated from each method as a readout. CDState ranked second overall, closely matching the top-performing method Linseed, with high Pearson’s correlation to ground-truth purity and low RMSE across all datasets, and outperforming all four other methods in terms of Pearson’s correlation in breast and ovarian cancers (Figure 3A).

**Figure 3.**
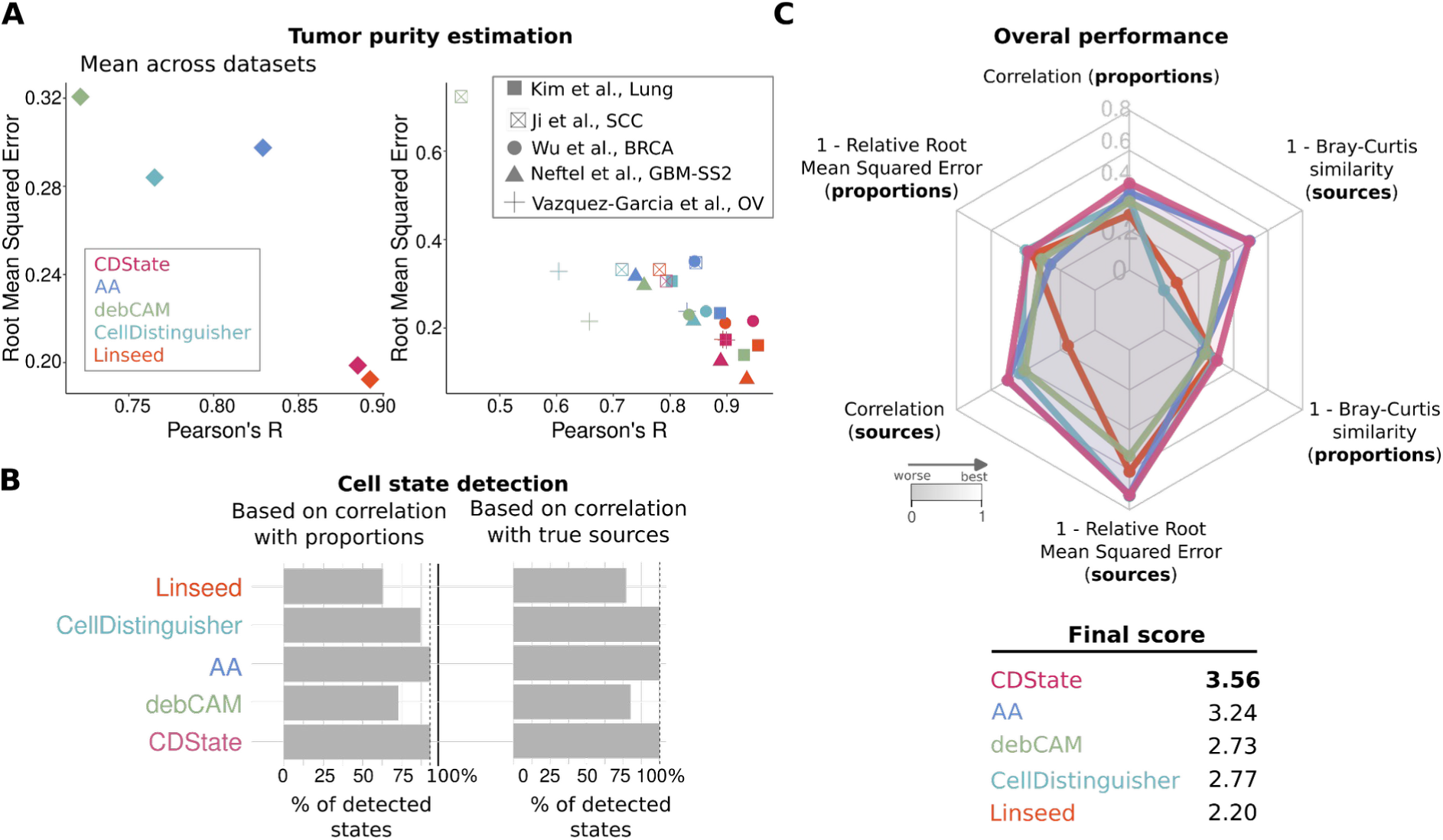
Benchmarking CDState in malignant cell state identification. **A**. Comparison of deconvolved and ground-truth tumor purity across five bulkified cancer datasets: mean performance (left) and performance per dataset (right). Point shapes correspond to the dataset, while colors correspond to deconvolution methods. **B.** The percentage of identified malignant cell states per method. **C.** Performance summary across the five bulkified cancer datasets. Each metric ranges from 0 to 1, with values close to 1 being the best. The final score was calculated as the sum of individual metrics.

Next, we assessed CDState’s performance in identifying malignant cell states and quantifying proportions of cells in corresponding transcriptional states in simulated tumor samples. For each malignant state, we computed Pearson’s correlation coefficient, relative RMSE (rRMSE), and Bray-Curtis index (BC) between the ground-truth and inferred source gene expression profiles and proportions (Figure 3C, Suppl. Figure S7, Suppl. Tables S3-S5, Methods). Finally, the overall performance of each method was calculated as the sum of respective metrics across the datasets.

CDState showed overall the best performance compared to existing deconvolution methods, with a total score of 3.56 (Figure 3C, Suppl. Figure S7, Suppl. Table S5). It achieved the highest overall correlation and BC index with true proportions (R = 0.54, BC = 0.4) and the highest correlation with source expression (R=0.63). For all other metrics, CDState was the second best, consistently showing high performance (Suppl. Table S5).

Additionally, CDState was among the best-performing methods for the number of correctly identified malignant states. Specifically, it correctly inferred 28 out of 30 malignant states based on the correlation between true and inferred proportions, and all of the ground truth states based on the correlation with source gene expression (Figure 3B).

To further validate the robustness of CDState in malignant cell state identification, we repeated the analysis with Gaussian noise added to ground-truth purity values (σ=0.25, Suppl. Tables S6-S7, Methods). Even under very noisy purity estimates, CDState outperformed all other methods in terms of the total score (Suppl. Table S8).

### CDState accurately infers malignant transcriptional programs and proportions without prior knowledge of cell state number

Next, we assessed CDState’s performance in a realistic setting in which the true number of states is unknown. We applied our method to the bulkified glioblastoma (GBM) dataset, where the ground-truth malignant cell states were extensively characterized [5]. In brief, glioblastoma cells have been shown to exist on a continuum of cellular states ranging from those resembling developmental progenitors, *i.e*. oligodendrocyte-progenitor-like (OPC-like) and neural-progenitor-like (NPC-like), to astrocyte-(AC-like) and mesenchymal-like (MES-like) states [5]. Further, the mesenchymal-like cells can be characterized by expression of MES-like-1 (hypoxia-independent) and MES-like-2 (hypoxia-dependent) signatures. Similarly, the NPC-like cells can express OPC-related or neuronal lineage genes, representing NPC-like-1 and NPC-like-2 sub-states, respectively.

We determined the optimal number of states by selecting the number of sources (*k*) that maximized the mean correlation with tumor purity across 20 runs (Methods). This approach led to selecting a seven-state solution (Suppl. Figure S8). After identifying malignant and TME components, we evaluated their correlation with the ground truth proportions of corresponding cell types (Figure 4A). CDState-inferred proportions were highly correlated with ground-truth cell type proportions, correctly separating malignant and non-malignant compartments. Among malignant states, NPC-like-1 and MES-like-1 cells were clearly resolved (de novo states S4 and S6, respectively), whereas MES-like-2 and AC-like states mapped to the same inferred state (S1), and OPC-like, NPC-like-2, and cycling cells all clustered within state S5 (Figure 4A). The difficulty in resolving these states can be explained by the strong correlation of their true proportions in bulk mixtures, particularly evident for OPC-like, NPC-like-2, and cycling cells (Suppl. Figure S9). While the selected seven-state solution merged some closely related malignant states, analyses with a higher number of states revealed a finer separation of these populations (Suppl. Figure S10-S11). However, the number of components must be chosen carefully, as overly large values can lead to sample-specific sources (Suppl. Figure S12). Consistent with the GBM results, CDState also accurately resolved known sources in SCC, BRCA, LUAD, and OV datasets (Suppl. Figures S13-S16).

**Figure 4.**
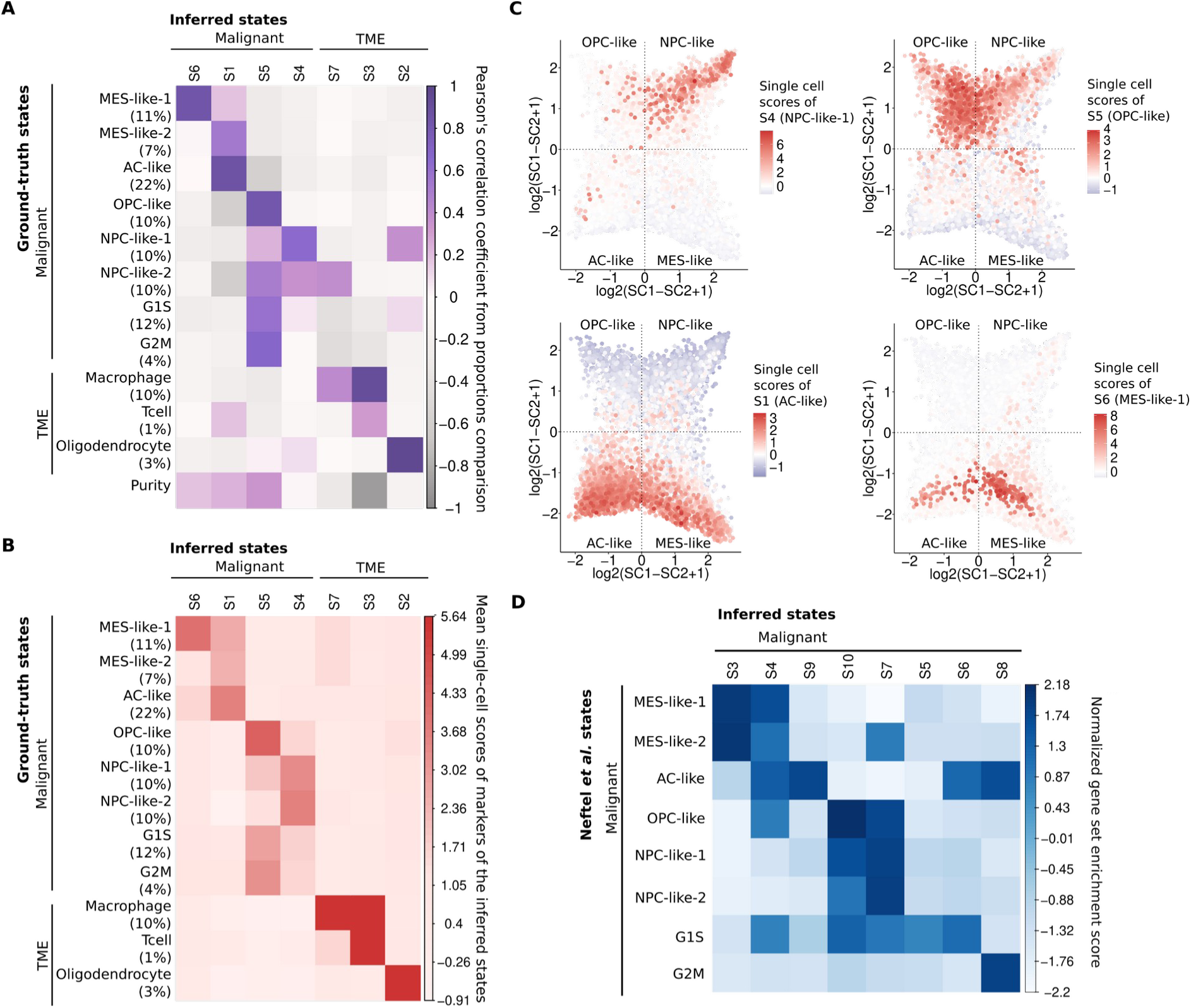
CDState deconvolution of cell states in glioblastoma data. **A.** Heatmap with Pearson’s correlation coefficients between the ground-truth and CDState-estimated proportions in bulkified glioblastoma data [5]. Percentages represent the proportion of respective cell populations in the bulkified dataset. **B.** Mean single cell scores of CDState-deconvolved states across ground-truth-labelled single cell glioblastoma data. Scores were calculated using the ANS method [43], and normalized with the z-score transformation. Percentages represent the proportion of respective cell populations in the bulkified dataset. **C.** Two-dimensional representation of glioblastoma malignant cell states from Neftel *et al.* [5]. Each quadrant corresponds to one of the main four states (OPC-like, NPC-like, AC-like, MES-like). Position of the malignant cells was determined by the relative scores of the cell states reported by Neftel *et al*., and colors reflect the scores of CDState-identified malignant states: S4 (NPC-like-1), S5 (OPC-like), S1 (AC-like), and S6 (MES-like-1). **D.** Normalized enrichment scores of the glioblastoma cell state-marker genes from Neftel *et al.* [5] scored across CDState-identified malignant states in experimental bulk glioblastoma data from Hu *et al*. [39].

To further validate observed correlations between true and inferred proportions, we selected marker genes of CDState-identified states in glioblastoma and scored single cells composing the bulkified dataset. By comparing the distribution of scores across ground-truth labelled single cells, we verified whether CDState accurately inferred true biological signals that distinguish underlying cell populations. The distribution of scores was consistent with the observed correlations between ground-truth and CDState-inferred cell proportions, further confirming CDState performance (Figure 4A, B). Additionally, we used the cell-state plot proposed by Neftel *et al.* [5], presenting the distribution of glioblastoma cells across the four main states, to visualize the concordance between ground-truth cell states and the single-cell scores of CDState-identified malignant states. CDState successfully identified the main axes of variation, with two OPC-NPC gradients (S5 stronger for OPC-like cells, S4 for NPC-like-1 cells) and two AC-MES gradients (S6 stronger for AC-like cells, and S1 for MES-like-1 cells) (Figure 4C).

Next, we assessed the reproducibility of CDState-identified malignant sources in GBM from Neftel *et al.* [5] and an independent cohort of true bulk IDH wild-type GBM samples from Hu *et al*. [39]. We found strong enrichment of the malignant-state signatures reported by Neftel *et al.* in CDState-identified malignant sources from Hu *et al*., with an accurate separation of mesenchymal and OPC-NPC axes (Figure 4D). Reproducibility of the identified malignant states was further confirmed in true bulk data from ovarian carcinoma [40], breast cancer [41], and lung adenocarcinoma [42] (Suppl. Figure S17)[42].

### Pan-cancer inference of malignant cell states using TCGA bulk RNA-seq data

The primary goal of CDState is its application to bulk RNA-seq data from large patient cohorts to identify robust states that drive malignant cell heterogeneity. To achieve this goal, we applied CDState to 33 datasets combining 9,669 patient samples from TCGA, and identified 193 malignant cell states in total (Figure 5A, Suppl. Table S9).

**Figure 5.**
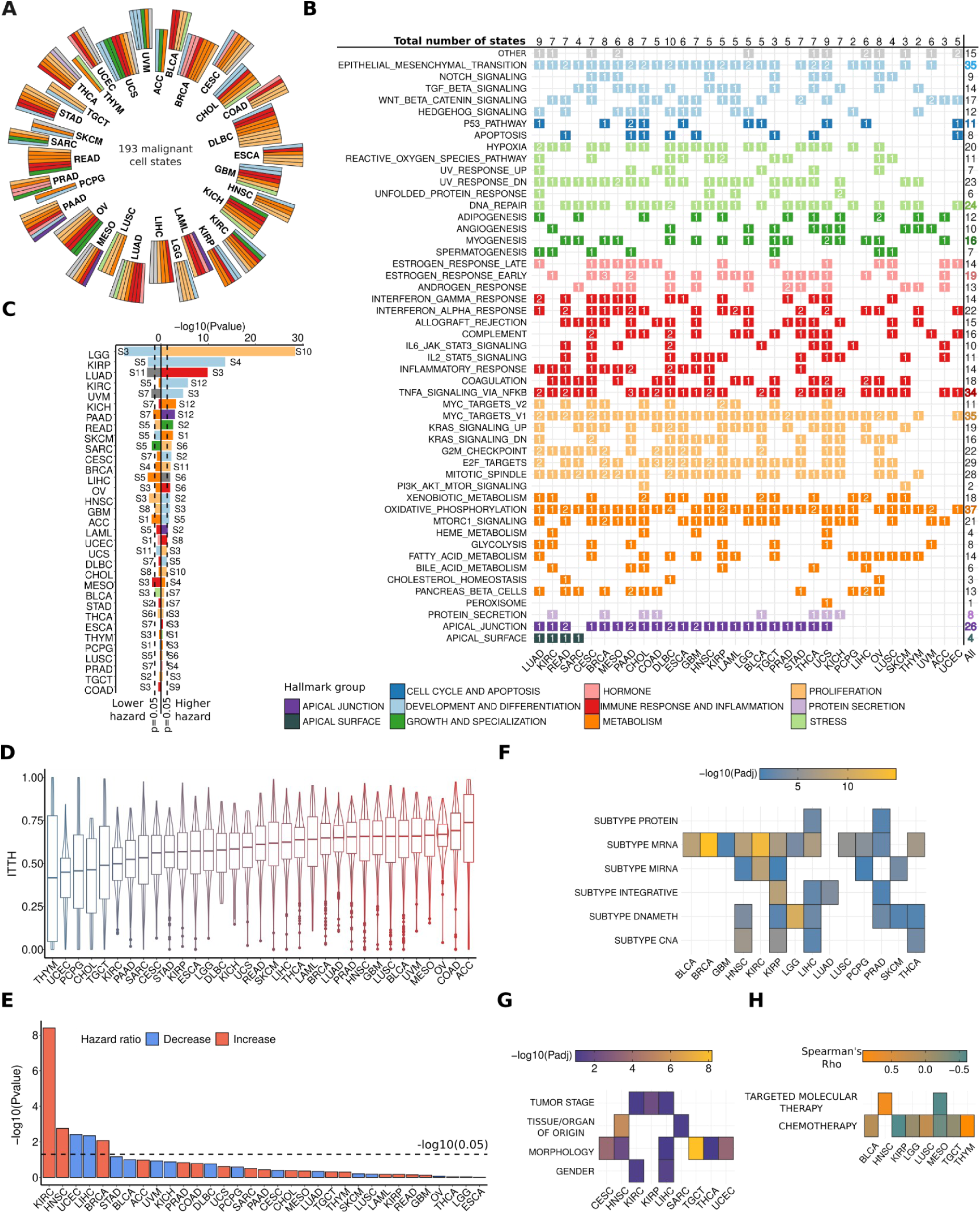
Pancancer characterization of malignant cell heterogeneity using CDState. **A.** Malignant cell states identified by CDState across 33 TCGA cancer types. States are colored based on the hallmark group of the hallmark set with the lowest adjusted p-value. **B.** Hallmarks of cancer enriched in the deconvolved malignant cell states across TCGA datasets. Numbers correspond to the number of states per cancer type in which the hallmark was significantly enriched (p-value < 0.05). Colors correspond to the hallmark grouping based on the shared molecular process. The enrichment was calculated using a hypergeometric test. **C.** States with a significant association with patient survival in TCGA. Colors correspond to the group of the most enriched hallmark in the given state.

CDState achieved good separation of malignant and non-malignant compartments, with 27 out of 33 TCGA cancer datasets showing a correlation of at least 0.5 between estimated and true tumor purity (Suppl. Figure S18). Additionally, in 31 of 33 cancer types, deconvolved malignant gene expression vectors showed stronger associations with copy number alterations compared to original bulk RNA-seq data. This observation indicates that the deconvolved expression more accurately reflects the true malignant compartment, whose expression is typically linked to the sample copy number profiles [4,44] (Suppl. Figure S18).

To characterize the molecular gene programs driving specific malignant states, we tested the enrichment of hallmarks of cancer gene sets deposited in MSigDB in gene expression vectors of identified malignant cell sources (Methods). Overall, we observed recurrent enrichment of molecular programs within and across cancer types, with epithelial-mesenchymal transition (35/192 states in 27/33 tumor types), MYC targets (35/192 states in 30/33 tumor types), and oxidative phosphorylation (37/192 states in 30/33 tumor types), as most frequently enriched programs (Figure 5B, Suppl. Table S10-S12). Among other most recurrent programs, we found pathways linked to immune response, stress response, apical junction, and hormone regulation (Figure 5B, Suppl. Table S10-S12). In addition, clustering of expressed gene programs across states revealed frequent co-expression of pathways such as MYC targets, DNA repair, oxidative phosphorylation, TNFα signaling via NF-κB, apical junction, and epithelial-to-mesenchymal transition (Suppl. Fig. S19). These findings suggest that while each cancer type expresses distinct malignant states, certain core transcriptional programs are conserved across malignancies.

### Proportions of malignant cell states are associated with patient clinical features

To assess the clinical significance of the inferred malignant cell states, we verified the association between their proportions and patient survival. In more than 50% of cancer types (17/33), at least one of the inferred malignant states was significantly associated with worse prognosis (Figure 5C, Suppl. Table S13, p-value<0.05). Additionally, in 10 cancer types, CDState identified cell states significantly linked to better clinical outcomes (Figure 5C, Suppl. Table S13, p-value <0.05). Functionally, states associated with poor prognosis were predominantly enriched in pathways related to development and differentiation, as well as immune response and metabolism (Suppl. Table S10-S11, S13).

Next, we used the inferred proportions of identified malignant states to assess sample-specific transcriptional heterogeneity. Specifically, we calculated the intratumor transcriptional heterogeneity (ITTH) index as Shannon’s diversity index on malignant cell-state proportions (Methods). TCGA samples showed a varied distribution of ITTH score, with the lowest median score in thymoma (THYM), and the highest in adrenocortical carcinoma (ACC) (Figure 5D). The ITTH index was significantly associated with overall survival in five cancer types (Figure 5E, Suppl. Table S13, p-value < 0.05). In clear cell renal carcinoma (KIRC), head and neck squamous cell carcinoma (HNSC), and breast cancer (BRCA), higher heterogeneity correlated with worse patient prognosis, while in liver hepatocellular carcinoma (LIHC) and uterine carcinoma (UCEC), it was linked to a better prognosis. Kaplan-Meier survival analysis stratifying patients by high versus low ITTH confirmed worse survival for patients with high ITTH in BRCA, HNSC, and KIRC, but improved survival in LIHC and UCEC (Suppl. Figure S20).

To assess the clinical relevance of compartment-specific heterogeneity, we further calculated ITTH indexes for global (TME and malignant states) and TME-only states across TCGA cancer types. Malignant ITTH was generally higher than TME ITTH (25/33 cancer types) and correlated more strongly with global ITTH values (Suppl. Figure S21). Global ITTH index was significantly associated with worse patient prognosis in the same three cancer types as the malignant index (KIRC, HNSC, and BRCA), with significantly better prognosis in kidney chromophobe (KICH) and skin melanoma (SKCM), while TME-specific index was significantly linked with worse patient prognosis in four cancer types (KIRC, stomach adenocarcinoma (STAD), SKCM, and lung squamous cell carcinoma (LUSC)) (Suppl. Figure S22).

Further, the malignant ITTH score was significantly associated with tumor molecular subtypes in 14 cancer types, and patient clinical features in 9 cancer types (Figure 5F,G). In particular, HER2-positive BRCA cancers showed the highest malignant ITTH score compared to other subtypes; KIRC tumors showed the highest ITTH in subtype 3 cancers characterized by expression of metabolic and oxidative phosphorylation signatures and previously linked with the worst patient survival [45], and low-grade gliomas (LGG) in m1 and m6 subtypes, IDH wild-type and neural-like, respectively [46]. Higher ITTH was also associated with more advanced disease in KIRC and KIRP cancers (Suppl. Figure S24). We found a higher malignant ITTH score linked with worse patient response to chemotherapy and targeted molecular treatment in four cancer types, including HNSC, bladder urothelial carcinoma (BLCA), LUSC, and THYM, highlighting the potential of ITTH as a predictive biomarker for therapeutic resistance (Figure 5H).

The survival association was evaluated using Cox univariate regression analysis. The dotted lines represent a p-value threshold of 0.05. **D.** TCGA cancer types present a varied degree of intratumor transcriptional heterogeneity (ITTH), calculated as Shannon entropy of deconvolved proportions. **E.** Association between ITTH and patient survival across TCGA datasets. P-value was calculated using Cox univariate regression analysis. **F-G.** Heatmaps show significant associations between ITTH and tumor molecular subtypes (**F**) and patient clinical characteristics (**G**) (Kruskal-Wallis adjusted p-value < 0.05). **H.** Association between ITTH and response to chemotherapy across TCGA cancer types (Kruskal-Wallis p-value < 0.05).

### Candidate genetic drivers of intratumor transcriptional plasticity

To identify potential genetic drivers of ITTH and malignant cell states, we used point-biserial correlation to link heterogeneity scores and state proportions with gene mutations and copy number alterations in transcription factors. Across TCGA, ITTH scores were linked to gene mutations in 13 cancer types and transcription factor copy number alterations in 20 cancer types (Figure 6A, Suppl. Figure S25). Most mutations were associated with a higher ITTH score, but in 4 cancer types, they were linked to lower ITTH values. Specifically, we found significant associations between ITTH and mutations in several known cancer-related genes, including *TP53, KRAS*, *PTEN*, *CTNNB1*, and *PIK3CA* (Figure 6A). In case of copy number changes, gene losses were more often linked to ITTH score than gene amplifications (Suppl. Figure S25). Similarly, we found significant associations between ITTH and gains and losses in several genes previously linked to cancer, including *NFE2L2, SOX17*, and *FOXP1*.

**Figure 6.**
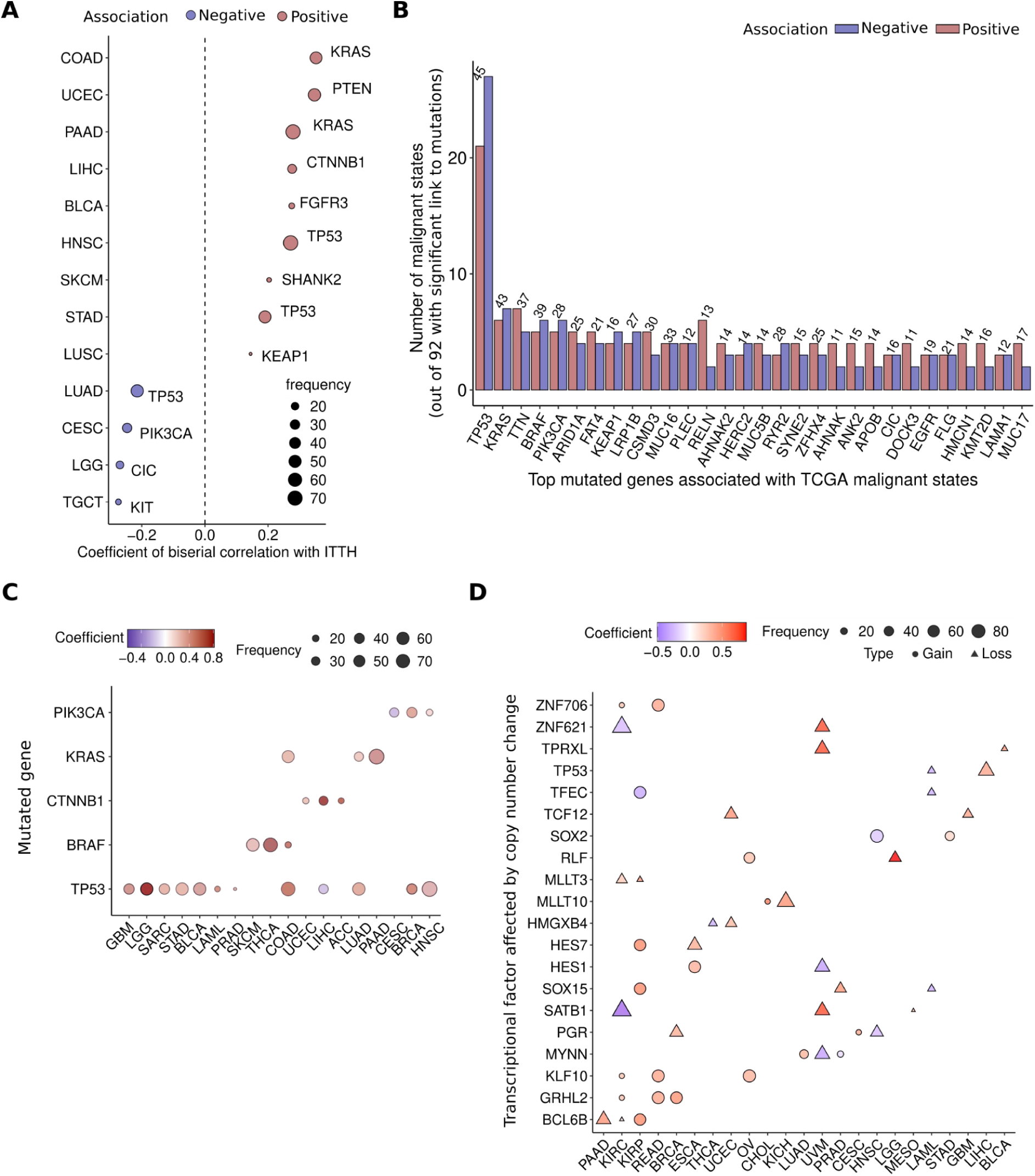
Candidate genetic drivers of TCGA malignant cell heterogeneity. **A.** Gene mutations with the highest absolute biserial correlation (adjusted p-value < 0.05) with cancer-type specific heterogeneity score (ITTH). Cancer-specific frequency of the events is represented by the size of the points. **B.** Top 30 mutated genes most frequently associated with TCGA malignant cell state proportions (adjusted p-value < 0.05). Numbers on top of bar plots represent the mean frequency [%] of mutations across all analyzed TCGA samples. Blue and red bars represent negative and positive associations, respectively. **C.** Gene mutations with the highest significant association (adjusted p-value < 0.05) with malignant state proportions, recurrently linked to states across TCGA cancer types. In case of equal positive and negative associations, only positive ones are reported. **D.** Transcriptional factors affected by copy number alterations with the highest absolute association with malignant state proportions (adjusted p-value < 0.05), recurrently linked to states across TCGA cancer types. Cancer-specific frequency of the events is represented by the size of the points. Copy number losses and gains are represented by point shape. In case of equal positive and negative associations, only positive ones are reported.

We compared the predictive power of identified genetic events and the CDState-estimated ITTH scores using the Cox multivariate survival analysis. In seven out of 22 cancer types with a significant association between ITTH index and genetic events, the ITTH scores had a stronger association with patient survival than the respective mutations or copy number changes (multivariate Cox-regression models, p-value<0.05). Three of the ITTH-associated genetic changes showed stronger association with patient survival than the ITTH score, including the loss of *KIAA1549* in KIRP cancers, mutation of *CIC* in LGG cancers, and mutations of *KRAS* in PAAD (Suppl. Figure S26).

Next, we analyzed genomic events linked to the proportions of specific malignant cell states and assessed the prevalence of these associations across analyzed cancer types. Mutations in *TP53* were most strongly linked to malignant state proportions in 12 cancer types, with *BRAF*, *CTNNB1*, *KRAS*, and *PIK3CA* each linked to a specific cell state in three cancer types. (Figure 6C). Overall, we found mutations in *TP53* as the most frequently linked with malignant state proportions (in 45 out of 193 cell states across cancer types), followed by mutations in *KRAS*, *TTN*, and *BRAF* (linked with 13, 12, and 11 state proportions, respectively). We identified copy number alterations in 20 transcription factors, including *TP53*, *SOX2*, *SATB1*, and *KLF10*, as most strongly associated with malignant state proportions across cancer types (Figure 6D). Copy number alterations in *CASZ1*, *ENO1*, *HES4*, and *JUN* were negatively associated with state proportions, whereas alterations in *TCF20*, *ZFP30*, *ZNF473*, and *FOXM1* were positively associated (Suppl. Figures S27-S28).

## Discussion

Multiple published studies provide evidence on the significance of transcriptional heterogeneity of tumors, particularly in relation to treatment resistance and patient prognosis [47–50]. With the rapid expansion of single-cell technology and its implementation in studying tumor samples, we have gained a broader understanding of malignant cell transcriptional profiles across multiple cancer types. However, current single-cell studies lack sufficiently large cohorts with matched modalities, such as comprehensive clinical annotations and genetic information, which are needed to fully characterize transcriptional heterogeneity, uncover its biological drivers, and assess its impact on clinical outcomes. At the same time, large cancer atlases, such as TCGA, could be utilized for this task. This, however, requires the availability of appropriate methods to separate transcriptional signals coming from different cell states present in bulk data. In this work, we introduce CDState, an unsupervised deconvolution method specifically developed to address this challenge. We demonstrate that CDState outperforms other existing unsupervised methods across bulkified cancer datasets with diverse composition, identifying previously reported malignant cell states and proportions of cells exhibiting these states within tumors. In addition, we show that our method maintains robustness against noisy purity estimates used for optimization and performs comparably well to supervised approaches with reference matching input bulk data. Finally, we demonstrate CDState’s accuracy in real-life scenarios, where the underlying number of states is unknown. Our analysis of TCGA datasets using CDState identified recurrent malignant cell programs, with epithelial-mesenchymal transition as the key driver of malignant cell heterogeneity across cancer types, consistent with recent work [51].

Given the link between intratumor heterogeneity and poor patient survival [52], previous studies focused on quantifying the extent of ITTH using different data types, including copy number aberrations [53], somatic mutations [54–56], or gene expression [55,56]. In our work, we extend this idea by proposing an intratumor transcriptional heterogeneity index (ITTH score) calculated based on proportions of inferred malignant cell states, and show its association with patient survival in several cancer types, consistent with previous findings [55,56]. Although we found the significant association between our ITTH index and patient survival in only five cancer types, we found that proportions of specific states are significantly associated with patient prognosis in 17 out of 33 studied cancer types. These results suggest that other ITTH-linked factors, such as cell-cell interactions or spatial distribution of cell states within tumors, may have stronger links with patient survival than global distribution of state proportions in most cancer types.

Moreover, our ITTH index was significantly linked with patient clinical features, including molecular subtype of disease, tumor stage, and response to therapy. The link with therapy is particularly notable, as growing evidence suggests that a single regimen often targets only a subset of tumor cells, and that ITTH plays a broad role in therapeutic resistance [57,58]. Although scRNA-seq is considered the gold standard for studying the cellular composition of cancer tissues and quantifying ITTH, its high costs and technical limitations make it unlikely to become a routine clinical practice. Alternatively, cost-effective bulk RNA-seq has the potential to be used in clinical settings as part of precision oncology programs, highlighting a use case for CDState-inferred indexes for patient treatment stratification and survival prediction.

Our results on the association between genetic alterations and specific malignant cell states show evidence on the interplay between tumor genotype and transcriptional heterogeneity, highlighting the potential role of specific genes in driving malignant cell heterogeneity. These findings may be particularly important for the development of novel diagnostic and therapeutic approaches, especially for cancer types with low survival rates. For instance, we found *TP53* mutation to be the most frequently associated with either higher or lower malignant state proportions and overall linked with malignant cell heterogeneity score. These results suggest that *TP53* may be involved in mechanisms underlying malignant cell states and intratumor heterogeneity, potentially playing a key role in regulating cellular plasticity and state transitions. For instance, a recent study showed that loss of p53 leads to a shift toward mesenchymal states in progenitor cells within pancreatic ductal adenocarcinoma, highlighting p53 function as a regulator of malignant cell plasticity [59]. Our results showing that EMT is the main contributor to malignant cell heterogeneity, along with previous evidence on the link between EMT and *TP53* [60–62], further support this finding. At the same time, the association between *TP53* mutations and lower ITTH scores suggests that *TP53* mutation uniformly promotes more aggressive states and limits others. While for now there are no approved therapies to restore *TP53* function or target p53 mutants, several clinical trials explore T*P53* mutations as a ground for novel treatments [63,64].

While we showed the utility and high performance of our approach across varied bulk and pseudobulk samples, some limitations should be taken into consideration. Firstly, CDState requires tumor purity estimates, which are used to guide the identification of malignant cell states and optimization of the parameters. Therefore, the deconvolution output heavily depends on the reliability of the provided tumor purity estimates. Recent studies have shown significant discrepancies between purity estimates using DNA and RNA sequencing data, making the decision on the use of an appropriate approach challenging [65,66]. In our analysis of the TCGA data, we used tumor purity estimates obtained using matched DNA-sequencing data and ABSOLUTE [53]. Therefore, given that the purity estimates were obtained from different pieces of tumors than the analyzed bulk RNA-seq data, we acknowledge this could potentially affect our deconvolution results. Nevertheless, our validation using noisy purity estimates confirmed that CDState maintains high performance even when provided with noisy purity estimates (σ = 0.25). Secondly, it is challenging to systematically set the optimal number of components for deconvolution. We showed that increasing the number of deconvolved cell types leads to improved separation of sources. However, the selection of too many states may lead to the identification of sample-specific sources, as observed in our results on bulkified glioblastoma data. Therefore, we recommend users to carefully examine the correlation between inferred and input purity to guide the optimal selection of state numbers for deconvolution, given the number of input samples. Finally, our analysis identified several potential genetic drivers of malignant cell states; however, further extensive experimental validation using CRISPR-Cas9 and similar approaches should be used to validate these findings.

Despite these challenges, CDState is a robust alternative to supervised deconvolution approaches for big cancer atlases with bulk data, when accurate single-cell references are unavailable. Its ability to identify clinically relevant states shows potential for improving patient stratification and diagnostic testing in clinical practice, and can inspire future studies on intratumor heterogeneity.

## Methods

### CDState algorithm

CDState is a computational method designed to identify and enumerate transcriptional cell states within tumor-resident cells. The algorithm operates in two main steps (Figure 1A-B), combining constrained non-negative matrix factorization (NMF) with an iterative optimization strategy inspired by simulated annealing [34], to maximize the dissimilarity between gene expression profiles of inferred transcriptional states.

### Step 1: Identification of initial solutions

1. CDState applies NMF to decompose the bulk RNA-sequencing expression matrix into a predefined number of components (*i.e.*, cell states).
2. The optimization is guided by two parameters, alpha (α) and beta (β) summing to 1, which control the balance between reconstruction error and source transcriptional profile separation. At this stage, the optimization focuses solely on minimizing the reconstruction error, with alpha (α) set to 1 and beta (β) set to 0.
3. The initial solution includes both malignant (MAL) and tumor microenvironment (TME) components, which are refined in Step 2.

### Step 2: Optimization of cell state separation

1. To improve the separation of inferred transcriptional states, β is progressively increased, weighting the optimization towards minimizing the mean cosine similarity between inferred cell state gene expression vectors.
2. As β increases, the method more likely accepts a higher reconstruction error while expanding the optimization space to maximize cell state dissimilarity.
3. This annealing-inspired process continues until the mean squared error between inferred and true tumor purity stops decreasing, at which point β is gradually reduced back to zero.
4. A final round of optimization, focused on minimizing reconstruction error runs until convergence.

### Constrained-Weight Nonnegative Matrix Factorization

The CDState algorithm assumes that the gene expression of a mixed bulk sample is a linear combination of expression vectors of pure source cell types: 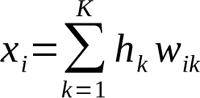 given *K* sources, where *xi* is bulk gene expression of gene *i* in a sample, w*_ik_* is gene expression of gene *i* in pure source *k*, and *h_k_* is a proportion of source *k* in the bulk sample. Therefore, the identification of underlying sources in the *n* input samples with bulk gene expression values for *m* genes can be formulated as a linear problem. CDState is based on the following Nonnegative Matrix Factorization framework:, where *X*, *W* and *H* are *m* x *n*, *m* x *K* and *K* x *n* non-negative matrices representing the input bulk expression, latent cell states and proportions, respectively.

In this formulation, NMF tends to return a sparse matrix of sources, in which each source represents an independent underlying element of the input data [67]. In a biological setting, these sources can be interpreted as individual biological processes, such as proliferation or stress-response. In reality, a living cell which is in a particular state can express more than one of these processes simultaneously. To account for this overlap, we extend the NMF model with the *sum-to-one* constraint on weights, such that: 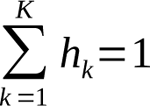 for each sample.

Our CDState algorithm uses the following objective function based on NMF: *L(W, H)=α∥X −W H^T^∥+ βMeanCosine (W)*, where 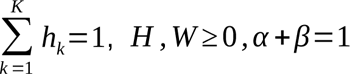 and *MeanCosine* (*W)* is the mean of the pairwise cosine similarities of source gene expression vectors.

### Simulated annealing-based optimization

Simulated annealing is incorporated into CDState to balance the trade-off between minimizing reconstruction error and maximizing the biological difference of inferred transcriptional states. Initially, the optimization focuses purely on reconstructing the bulk RNA-seq data by minimizing the reconstruction error (α = 1, β = 0). However, to avoid inference of highly correlated cell states, in the second step of the algorithm CDState gradually increases the β parameter, which places greater emphasis on minimizing the mean cosine similarity between inferred states. By progressively accepting a higher reconstruction error, CDState encourages the discovery of transcriptional states that are more biologically distinct. However, to ensure that the final decomposition remains relevant, β is only increased as long as the mean squared error (MSE) between the inferred and true tumor purity continues to improve. Once MSE begins to rise, β is reduced back to zero, and a final round of reconstruction optimization is performed until convergence.

### Implementation details

#### Inputs

The CDState method requires as input (a) normalized bulk RNA-seq expression profiles (matrix of shape m genes × n samples), (b) tumor purity estimates, and (c) a predefined range of possible cell state numbers (e.g., “k=2:4” runs CDState for 2, 3, and 4 states).

#### Outputs

The following output information is returned by the method: (a) matrix with gene expression profiles of the identified cell states (m genes × k sources), (b) matrix with proportions of the identified cell states in the input samples (n samples x k sources), and (c) component assignment to malignant or tumor microenvironment states.

#### Optimization Strategy

The weights and source update in CDState step 1 is solved using the SCIPY minimize-maximize function with Sequential Least Squares Programming (SLSQP) optimizer and non negative least squares objective function. In step 2, the source matrix is updated using gradient descent (using the JAX python library).

### Initialization and multiple runs

Due to the stochastic nature of NMF, CDState is run multiple times with different starting sources. The starting sources are selected from bulk samples using K-means clustering, with the number of clusters corresponding to the selected number of states. A random sample from each cluster is then selected as the starting source for each run.

### Recommendation on selection of number of components

We recommend running CDState for a range of components, depending on the number of input bulk samples (*N*), but preferably not higher than *N*/2. For each value of *k* we advise running CDState with multiple runs (*i.e.,* between 10 and 20) for robust results. In order to select the final results, users should consult the mode number of malignant components (*K_mal*) identified across runs for each *k* and select runs that identified number of malignant components equal to that number. Finally, the best *k* can be selected by choosing the number of components with the highest mean correlation between true and CDState-inferred purity for the runs with *K_mal* malignant components per *k*.

### Identifiability of CDState solutions

In its general formulation, NMF is not identifiable, meaning that multiple different pairs of *W* and *H* decompose the original input data *X* with similar accuracy [68][69]. However, for applications in which solution uniqueness is necessary for better interpretability, certain constraints under which NMF becomes identifiable, including sparsity of W and H, source separability condition or volume minimization (VolMin) have been proposed [69–74]. In our formulation of NMF, where we introduce the sum-to-one constraint on columns of *H^T^*, each bulk sample represents a convex combination of sources in *W*. This means that each sample lies within a simplex whose vertices are defined by columns of *W* and represent pure sources. This constraint improves identifiability as it fixes the scale of *H*, therefore removing scaling ambiguity. This is similar to the MinVol criterion, which guarantees identifiability if components are well separated. In addition, if input data points lie near the center of the simplex or if pure samples are missing, it becomes harder to uniquely recover true sources, as (1) there are no clear anchor points for the simplex and many factorizations could equally well explain the data, (2) and the samples do not span the full simplex. We observed this problem when analysing constrained-weight NMF (CW-NMF) results for our synthetic mixes MixA-MixD. The component separability condition essentially ensures that each source appears as pure (with proportion close to 1) in at least one sample. In constrained-weight NMF one of the main limitations is that components may become redundant leading to ambiguity. In CDState, the addition of cosine similarity-based penalty on columns of *W* encourages geometric separation of components, making them more orthogonal. Under appropriate selection of β, sources are forced to point in distinct directions, mimicking the geometric spread of input samples with pure samples present within simplex. Overall, with the convex optimization constraint, the cosine similarity term reduces the size of the solution space by favouring more distinct and identifiable columns of matrix *W*. While our solution may not guarantee full identifiability, it improves robustness and uniqueness of decomposition. In addition, we advise the user to run CDState for a range of *k*, each with several runs, and to select a solution which provides best reproducibility and performance.

#### Gene filtering

Before running CDState, normalized input bulk data is first filtered to minimize the impact of outlier genes and reduce computation time, using the method described in [31]. Specifically, genes with vector norm below 30% or above 99% of the mean vector norm across all genes, are excluded. Within the remaining genes, their expression levels are binned based on 0.1 increase step, and only the top 50% of genes with the highest standard deviation within each bin are retained (Suppl. Figure S29). This filtered gene set is then used for the entire inference process. This filtering strategy ensures that the input data align more closely with the Gaussian noise assumption modelled by the squared-error loss used in CDState. A function to infer gene expression for the complete gene set is also available within the CDState package.

#### Definition of TME and malignant cell states

After the initial step of the algorithm, CDState classifies the inferred sources into tumor microenvironment (TME) and malignant cell states. This is achieved through a greedy search to identify the optimal combination of source proportions that have the strongest Spearman’s correlation with the ground truth purity. Specifically, the proportions of each inferred component are correlated with the provided purity estimates and ranked in descending order of correlation strength. Then, for each consecutive subset of components, the summed proportions are correlated with true purity. The subset of states showing the highest correlation (with a one-sided p-value < 0.05) is selected as representing malignant cell states, while the remaining sources are selected as TME components. If purity estimates are available for fewer than 20 samples, the significance of the correlation is assessed using bootstrap resampling with 1,000 iterations.

#### Selection of the number of cell states for deconvolution

To determine the optimal number of cell states for deconvolution, CDState provides several metrics, including reconstruction error, total error, the assignment of components to malignant and TME states, and the final Pearson’s correlation with true purity based on inferred malignant cell state proportions. Users may select the most suitable number of states for deconvolution using these metrics as described above (“Recommendation on selection of number of components”).

#### Generation of bulkified datasets

To generate bulkified datasets, we followed two distinct approaches: one for cell type deconvolution and another specifically for malignant cell state inference. For the general cell type mixtures, we selected three distinct cell types from the *Kim et al.* lung adenocarcinoma dataset [24]: malignant cells, B cells, and fibroblasts. Then, we used the Dirichlet distribution to simulate varying cellular compositions, with the concentration parameter (alpha) controlling the level of variation. Specifically, we generated four mixtures in which proportions of each bulk sample were drawn from the following distributions:

- MixA *θ_i_* ∼ *Dir* (1,1,1)
- MixB *θ_i_* ∼ *Dir* (10,10,10)
- MixC *θ_i_* ∼ *Dir* (0.1,0.1,0.1)
- MixD *θ_i_* ∼ *Dir* (1,20,1), where *θ_i_* - cell type proportions for sample *i*.

Then, corresponding numbers of cells were randomly sampled from these three cell types, and the count gene expression of these selected cells was then aggregated to create the final bulk sample. To ensure comparability across samples, we normalized the resulting expression values by the total read count per sample.

For the bulkification of cancer datasets with malignant cell states, we used five datasets [5,24–27] where state annotations were either directly available or inferred. In cases where annotations were missing, we employed a marker gene scoring approach using Scanpy [75] as described earlier [5], to assign each cell to the state for which it had the highest score. To construct a bulk sample, we aggregated gene expression of malignant and nonmalignant cells coming from a particular sample and normalized the expression values by the total read count. For glioblastoma data where only TPM values were available, we aggregated these values directly without additional normalization. Ground truth gene expression profiles were calculated by averaging the expression of cells within each identified cell type or state.

#### Evaluation of the annealing procedure used in CDState

To verify the impact of our annealing-based optimization, we compared performance of CDState using solutions from four different scenarios: the first phase of CDState - described in the manuscript also as CW-NMF (Initial), solution from running CDState with fixed beta (Fixed beta), solution from running CDState without final decreasing of beta (No cooldown), and final CDState solution (Final). For the “Fixed beta” solution, we used the value of beta that was the most frequently used by the algorithm in the “Final” scenario. Then, CDState was run from the beginning with the selected value of beta until final convergence. For the “No cooldown” scenario, we run the initial and final steps of CDState, but the algorithm was stopped immediately once the MSE with true purity increased. Each scenario was run 20 times for each of the four mixtures.

#### Evaluation of CDState performance under noisy purity estimates

To verify robustness of CDState against unreliable true purity estimates of input samples, we generated noisy purity estimates for the synthetic mixtures by adding noise drawn from Gaussian distributions: *N(0,0.05^2^)*, *N(0,0.1^2^)*, and *N(0,0.25^2^)*. Noisy purity values were then cut to the range [0,1]. For benchmarking using bulkified cancer datasets, noisy purity estimates were generated using the highest level of noise, drawn from *N(0,0.25^2^)*.

#### Comparison with existing methods

We compared CDState performance with four existing methods: CellDistinguisher [28], Linseed [29], Archetype Analysis (AA) [32,33], and debCAM [30,31]. The methods were run with default parameters. As these methods do not provide a dedicated methodology to assign malignant and tumor microenvironment states, we applied the CDState approach to assign these identities for all methods (section “Definition of TME and malignant cell states” and “Bulkified tumor datasets”). Additionally, debCAM, CellDistinguisher, and Linseed perform their own gene filtering, which limits our ability to compare results across all genes. Consequently, the performance comparison was made only using the subset of common genes retained by these methods. Furthermore, AA and Linseed do not output proportions that sum to one; therefore, we rescaled the proportions from these methods to fulfill this condition.

To evaluate the performance of CDState in proportions inference against supervised approaches, we used InstaPrim, a fast implementation of the BayesPrism method [22,37]. InstaPrism was run on the synthetic mixtures using single-cell data of the same cell types that were used to generate the four mixtures. To evaluate the stability of InstaPrism results, we run the method 20 times for each mixture, sampling 25% of all cells used to generate bulkified samples for the reference dataset each time.

#### Bulkified mixtures

For the bulkified mixtures, the methods were run for three components, and each method (except deterministic CellDistinguisher) was run 20 times. Methods were compared in terms of root mean squared error (RMSE) and Pearson’s correlation coefficient with the ground truth proportions and source expression profiles.

#### Bulkified tumor datasets

To ensure a fair comparison, we ran CDState and the four selected deconvolution methods across a range of component numbers (*k*) ranging from 2 to 15, except debCAM, which, due to the runtime, could not be run for k>10 (Suppl. Figures S30-S33). Since the existing methods do not inherently distinguish malignant from TME components, we applied the same assignment approach as the one used in CDState. Specifically, for each method and each run for a value of *k*, we selected the components that maximized correlation with the true tumor purity and assigned them as malignant states (Suppl. Figures S30-S33, Suppl. Table S1). This allowed us to determine the number of inferred malignant states per *k*. Knowing the true number of malignant states in each dataset, we selected the *k* that best matched the expected number of malignant states (Suppl. Table S1). If the exact number was not identified by a method, we chose the solution with the closest higher value.

The performance of each method was evaluated based on the inferred proportions and source gene expression profiles. To quantify the performance, we used three different metrics:

- Pearson’s correlation coefficient, defined as 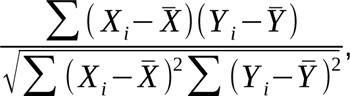,
- Bray-Curtis dissimilarity index, defined as 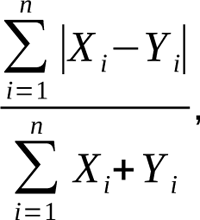,
- and the relative root mean squared error (rRMSE), defined as 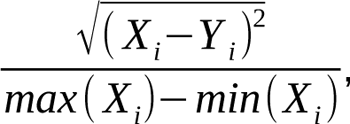,

where *X_i_* is corresponds to expression level of gene *i* in ground-truth source *X*, *X*^-^ is mean expression of genes in ground-truth source *X*, *Y _i_* corresponds to expression level of gene *i* in estimated source *Y*, and *Y*^-^ is mean expression of genes in estimated source *Y*. Similarly, when calculating these metrics for proportions, *X_i_* is corresponds to ground-truth proportion of source *X* in bulk sample *i*, *X*^-^ is mean of true sample proportions of source *X*, *Y _i_* corresponds to estimated proportion of source *Y* in bulk sample *i*, and *Y*^-^ is the mean of the estimated proportion of source *Y*.

For Pearson’s correlation coefficient, we considered only non-negative coefficients to assign inferred states; therefore, all the calculated metrics ranged between 0 and 1, allowing direct comparison across datasets and states. Finally, to summarize the overall performance of each method, we transformed rRMSE values to *1 - rRMSE* and BC values to *1 - BC*, ensuring that values closer to 1 represent better performance, and then mean values of these scores across datasets were summed, providing an overall performance score for each method.

#### CDState deconvolution of bulkified datasets when the true number of malignant states is unknown

The bulkified datasets of glioblastoma [5], ovarian cancer [26], breast cancer [25], lung adenocarcinoma [24] and squamous cell carcinoma [27] were deconvolved with CDState for 2 to 15 states. For each dataset, we selected the best solution based on the highest correlation between estimated and true tumor purity. The best run for each *k* was selected as the one with the highest correlation with true purity, among the runs that identified the mode value of malignant states for the particular *k*. For each dataset, we calculated Pearson’s correlation between the ground truth proportions obtained during bulkification and CDState-estimated proportions.

To identify marker genes for each inferred cell state in bulkified glioblastoma data, we selected the top 100 genes with the highest difference in expression between the particular source and the mean of its expression in the remaining components. Identified marker genes were then used to score the single-cell populations used for bulkification, applying the ANS method [43].

Finally, we followed the approach described in *Neftel et al.* [5] to visualize glioblastoma malignant cells within a two-dimensional space defined by OPC-, NPC-, AC-, and MES-like cell states. The x- and y-axis position of cells was calculated as described in the original study by Neftel *et al.* [5].

#### Pancancer deconvolution of TCGA data

RSEM-normalized bulk RNA-seq data for all TCGA tumor types were downloaded from Xena Browser (https://xenabrowser.net/datapages/), along with ABSOLUTE tumor purity estimates from the TCGA PanCanAtlas supplemental data (https://gdc.cancer.gov/about-data/publications/pancanatlas). CDState was applied separately to each cancer type, performing deconvolution with *k* ranging from 2 to 15, with 10 independent runs for each value of *k* (Suppl. Figures S34-S37). The selected number of *k* for each cancer type can be found in Suppl. Table S9.

To determine the optimal number of transcriptional cell states for each cancer type, we used the following selection criteria:

- We used *k* which has the highest correlation between estimated and true tumor purity;
- At least two malignant cell states should be identified;
- A plateau in the number of malignant states is observed across increasing values of *k*.

#### Calculation of ITTH score

The ITTH score is calculated as follows: 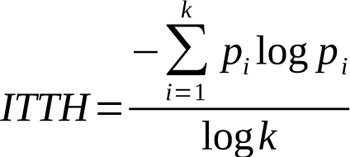, where *pi* represents the proportion of malignant state *i*, and *k* is the total number of malignant cell states. Malignant state proportions were rescaled to sum to one. A higher ITTH score indicates increased transcriptional heterogeneity within the tumor. For global and TME indexes, we applied the same approach, using all identified and TME-specific state proportions, respectively.

#### Gene set enrichment analysis and state annotation

Marker genes of inferred malignant cell states were identified using the procedure described in the section “CDState deconvolution of bulkified datasets when true number of malignant states is unknown”. For each state, a hypergeometric test implemented in the fgsea R package [76] was performed to assess the association between the marker genes and hallmark of cancer gene sets, downloaded from MsigDB (https://www.gsea-msigdb.org/gsea/msigdb). Hallmarks with a p-value < 0.05 were considered significantly enriched. To simplify biological interpretation, the 50 hallmark gene sets were manually grouped into 11 functional categories (Supplementary Table S10). Each malignant cell state was assigned to a specific functional group based on the classification of the hallmark set with the lowest p-value association with the given state (Supplementary Table S11).

The same procedure was used to calculate the enrichment of marker genes of the known malignant cell states in CDState-identified sources of true bulk IDH wild-type glioblastoma [37], ovarian carcinoma [38], breast cancer [39], and lung adenocarcinoma [40] datasets. For these datasets, we used the best solution for *k* matching the expected number of malignant cell states.

#### Clinical significance of malignant cell states

The ITTH index and the proportions of identified malignant cell states were analyzed for associations with patient clinical features. The Kruskal-Wallis and Mann-Whitney U tests were used to assess the relationship between the ITTH score and tumor molecular subtypes, as well as other clinical characteristics and treatment response information of TCGA patients. To evaluate the prognostic relevance of ITTH indexes and malignant cell state proportions, Cox univariate survival analysis was used (using *survival* and *survminer* R packages). For survival analysis with ITTH and associated genetic mutations and copy number aberrations, multivariate Cox survival analysis was used, and we reported feature-specific p-values. Kaplan-Meier analysis was performed using *survminer* R package, with *surv_cutpoint* function used to identify the optimal split of patients based on the ITTH score.

#### Identification of potential genetic drivers of malignant cell states

Gistic2 copy number data and MC3 gene-level non-silent mutation data for TCGA samples was downloaded from Xena Browser (https://xenabrowser.net/datapages/). For better interpretability, copy-number changes were encoded as binary events for gains (Gistic2 values 1 and 2) or losses (Gistic2 values −1 and −2). For each cancer type, we first filtered out genes with genetic change in less than 10% of samples. Then, we used point biserial correlation for each CDState-identified malignant state proportions/ITTH index and gene alteration profiles, to assess the association. P-values obtained within each cancer type were adjusted for the number of tests. For analysis of copy number changes, we used only changes in transcriptional factor genes defined in the FANTOM5 database (https://fantom.gsc.riken.jp/5/).

## Supporting information

CDState - Suppl. Tables

## Data and code availability

### Code availability

CDState is implemented in Python and is freely available at https://github.com/BoevaLab/CDState. Tutorials on how to use the method are also available on github.

### Data availability

The single-cell studies used in this study can be downloaded from:

– Broad Institute Single-Cell Portal: Single cell RNA-seq of adult and pediatric glioblastoma, Neftel et al., https://singlecell.broadinstitute.org/single_cell/study/SCP393/single-cell-rna-seq-of-adult-and-pediatric-glioblastoma; A single-cell and spatially resolved atlas of human breast cancers, Wu et al., https://singlecell.broadinstitute.org/single_cell/study/SCP1039/a-single-cell-and-spatially-resolved-atlas-of-human-breast-cancers;
– The Gene Expression Omnibus (GEO): Single cell RNA sequencing of lung adenocarcinoma, Kim et al., GSE131907;
– Cellxgene portal: MSK Spectrum - Ovarian cancer mutational processes drive site-specific immune evasion, Vazquez-Garcia et al., https://cellxgene.cziscience.com/collections/4796c91c-9d8f-4692-be43-347b1727f9d8;
– Curated Cancer Cell Atlas: Skin squamous cell carcinoma, Ji et al., https://www.weizmann.ac.il/sites/3CA/skin;

The bulk data used in this study can be downloaded from:

– Xena Browser (https://xenabrowser.net/datapages/): TCGA IlluminaHiSeq (RSEM-normalized) data; Gistic2 thresholded copy number data; MC3 gene-level non-silent mutation data;
– PanCanAtlas supplementary files
(https://gdc.cancer.gov/about-data/publications/pancanatlas): TCGA Absolute purity/ploidy file; TCGA-clinical Data Resource Outcome;
– TCGAbiolinks R package (https://rdrr.io/bioc/TCGAbiolinks): full clinical characterization of TCGA samples; subtype annotation of TCGA samples;

## Declarations

### Ethics approval

Not applicable.

### Consent for publication

Not applicable.

## Author information

### Authors’ contributions

AK developed CDState, performed computational analyses, and prepared the manuscript with input from JY, FB, and VB. AK and VB conceived the study. VB supervised the study. All authors read and approved the final manuscript.

## Acknowledgments

We sincerely thank Prof. Daniela Witten, University of Washington, for inspiring the idea to use purity information within the CDState algorithm. We also thank Prof. Judith Zaugg, University of Basel and EMBL, and Prof. Ewa Szczurek, University of Warsaw and Helmholtz Munich, for their invaluable feedback on the manuscript.

## Supplementary Material

### Supplementary Tables

**Supplementary Table S1.** Number of malignant cell states across values of k and methods used for benchmarking.

**Supplementary Table S2.** Number of components selected for the benchmark on bulkified tumor datasets.

**Supplementary Table S3.** Summary statistics of proportion inference across bulkified cancer datasets and methods.

**Supplementary Table S4.** Summary statistics of source gene expression inference across bulkified cancer datasets and methods.

**Supplementary Table S5.** Overall performance of CDState and exisitng methods across bulkified cancer datasets.

**Supplementary Table S6.** The number of malignant cell states detected by available unsupervised methods across bulkified cancer datasets. Purity with added noise (N∼(0, 0.25)) was used to select malignant components.

**Supplementary Table S7.** The number of selected “k” in CDState analysis on bulkified datasets with noisy purity.

**Supplementary Table S8.** Overall performance of CDState and exisitng methods across bulkified cancer datasets with noisy purity estimates.

**Supplementary Table S9**. Identified optimal number of CDState components across TCGA cancer types.

**Supplementary Table S10.** Functional grouping of MsigDB hallmarks of cancer sets.

**Supplementary Table S11.** Results of gene set enrichment analysis using MsigDB hallmarks of cancer, on the CDState-inferred malignant cell states across TCGA cancers.

**Supplementary Table S12**. Recurrent hallmark groups expressed in malignant cell states across TCGA cancer types.

**Supplementary Table S13**. Results of Cox univariate survival analysis across malignant cell states and ITTH index identified for TCGA cancers.

## Supplementary Figures

**Suppl. Figure S1A.**
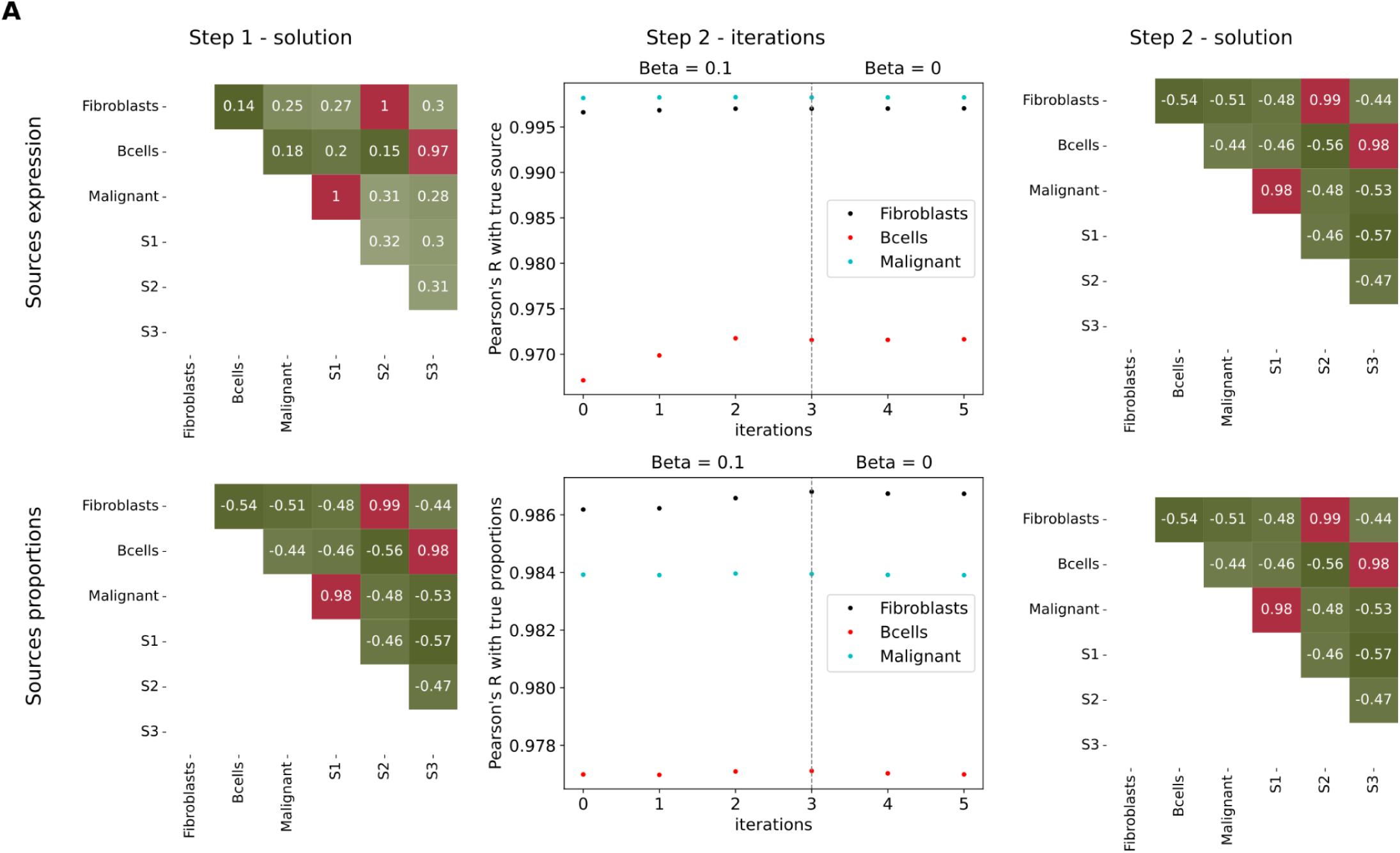
Pearson’s correlation between ground truth sources and proportions of the three cell types used to create Mix A samples. Correlations between output from the first and second steps CDState are compared. Values of Beta used in the second step are reported on the middle plot. **Suppl. Figure S1B.** Pearson’s correlation between ground truth sources and proportions of the three cell types used to create Mix B samples. Correlations between output from the first and second steps CDState are compared. Values of Beta used in the second step are reported on the middle plot. **Suppl. Figure S1C.** Pearson’s correlation between ground truth sources and proportions of the three cell types used to create Mix C samples. Correlations between output from the first and second steps CDState are compared. Values of Beta used in the second step are reported on the middle plot. **Suppl. Figure S1D.** Pearson’s correlation between ground truth sources and proportions of the three cell types used to create Mix D samples. Correlations between output from the first and second steps CDState are compared. Values of Beta used in the second step are reported on the middle plot.

**Suppl. Figure S1B.**
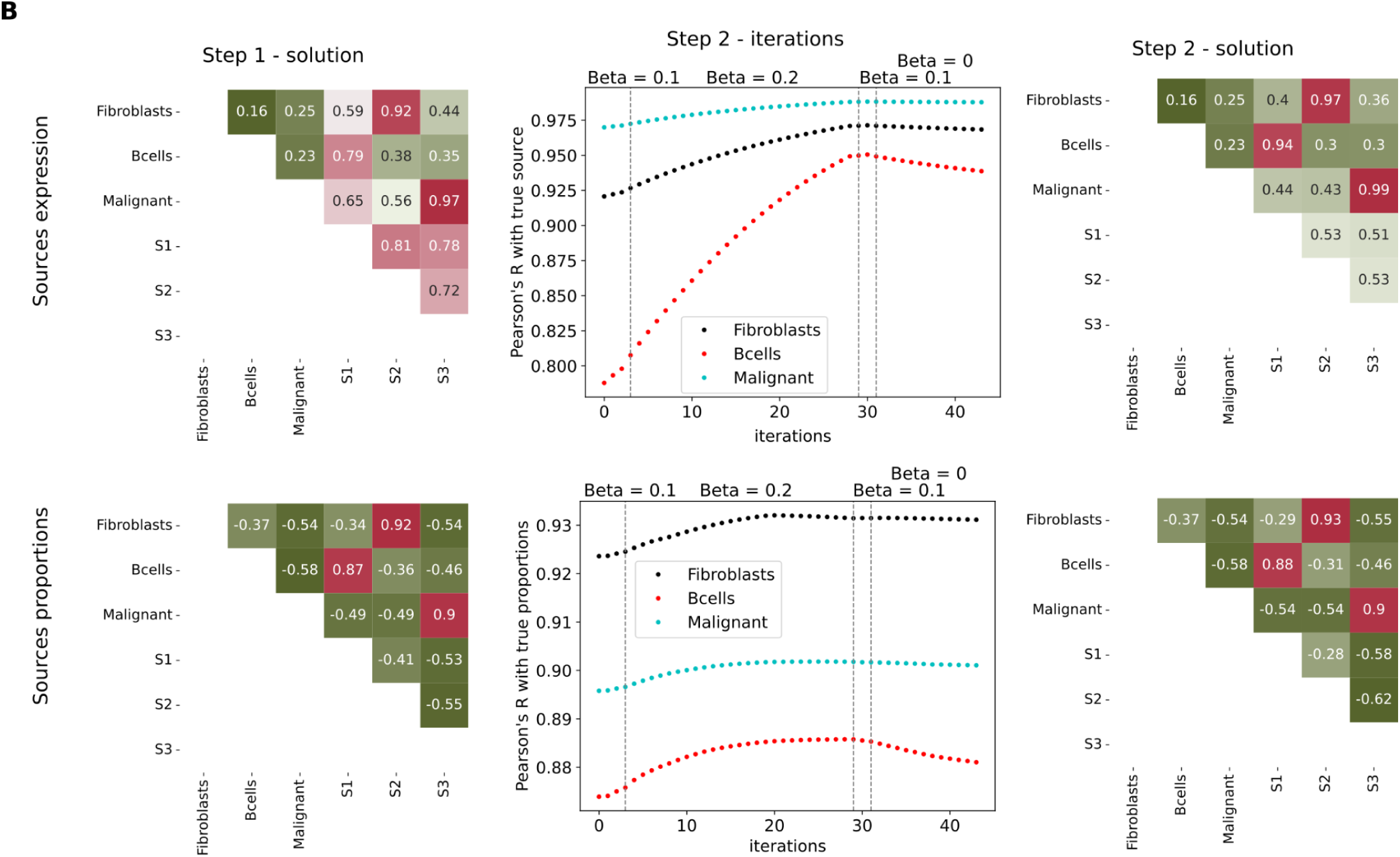
Pearson’s correlation between ground truth sources and proportions of the three cell types used to create Mix B samples. Correlations between output from the first and second steps CDState are compared. Values of Beta used in the second step are reported on the middle plot. **Suppl. Figure S1C.** Pearson’s correlation between ground truth sources and proportions of the three cell types used to create Mix C samples. Correlations between output from the first and second steps CDState are compared. Values of Beta used in the second step are reported on the middle plot. **Suppl. Figure S1D.** Pearson’s correlation between ground truth sources and proportions of the three cell types used to create Mix D samples. Correlations between output from the first and second steps CDState are compared. Values of Beta used in the second step are reported on the middle plot.

**Suppl. Figure S1C.**
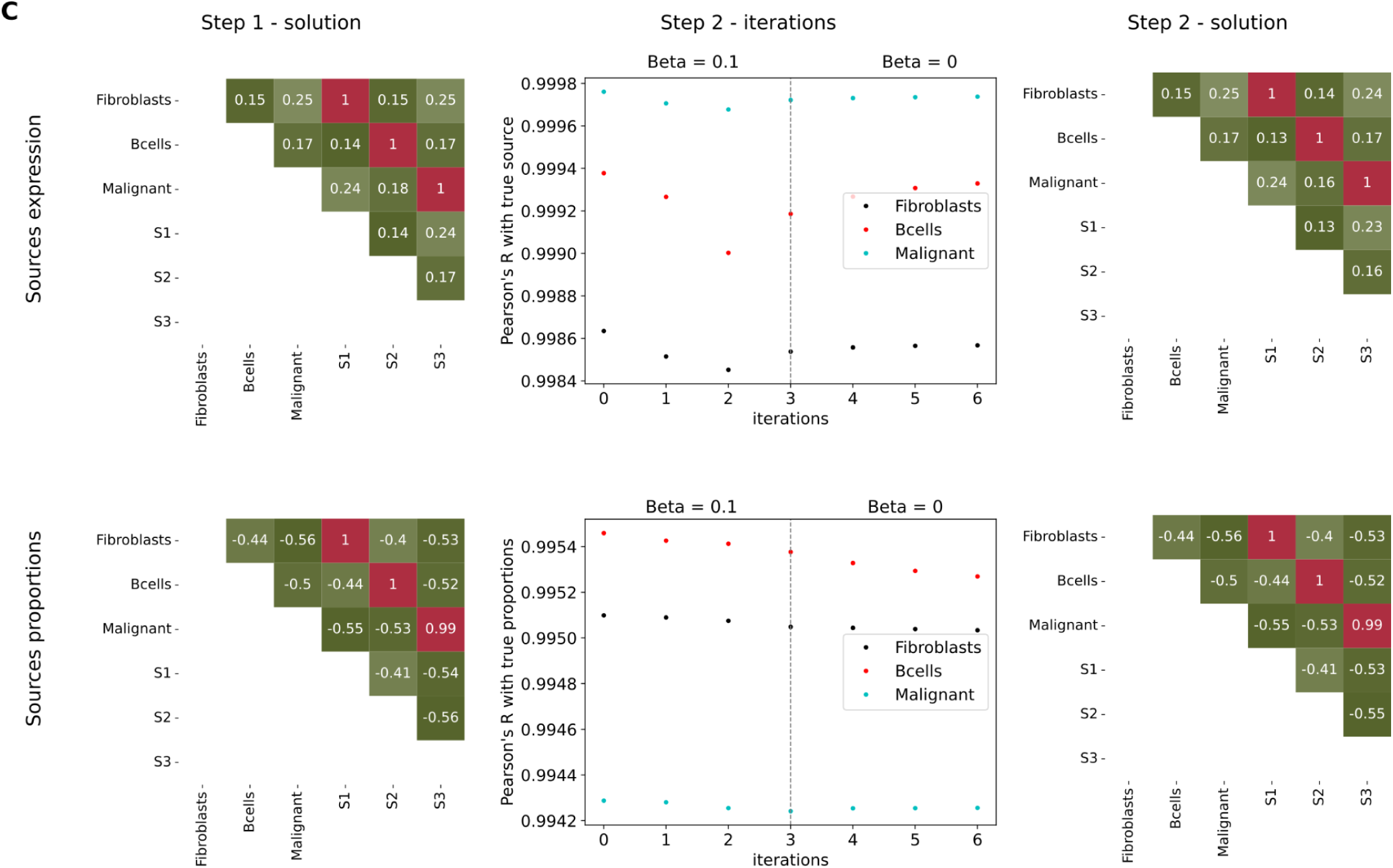
Pearson’s correlation between ground truth sources and proportions of the three cell types used to create Mix C samples. Correlations between output from the first and second steps CDState are compared. Values of Beta used in the second step are reported on the middle plot. **Suppl. Figure S1D.** Pearson’s correlation between ground truth sources and proportions of the three cell types used to create Mix D samples. Correlations between output from the first and second steps CDState are compared. Values of Beta used in the second step are reported on the middle plot.

**Suppl. Figure S1D.**
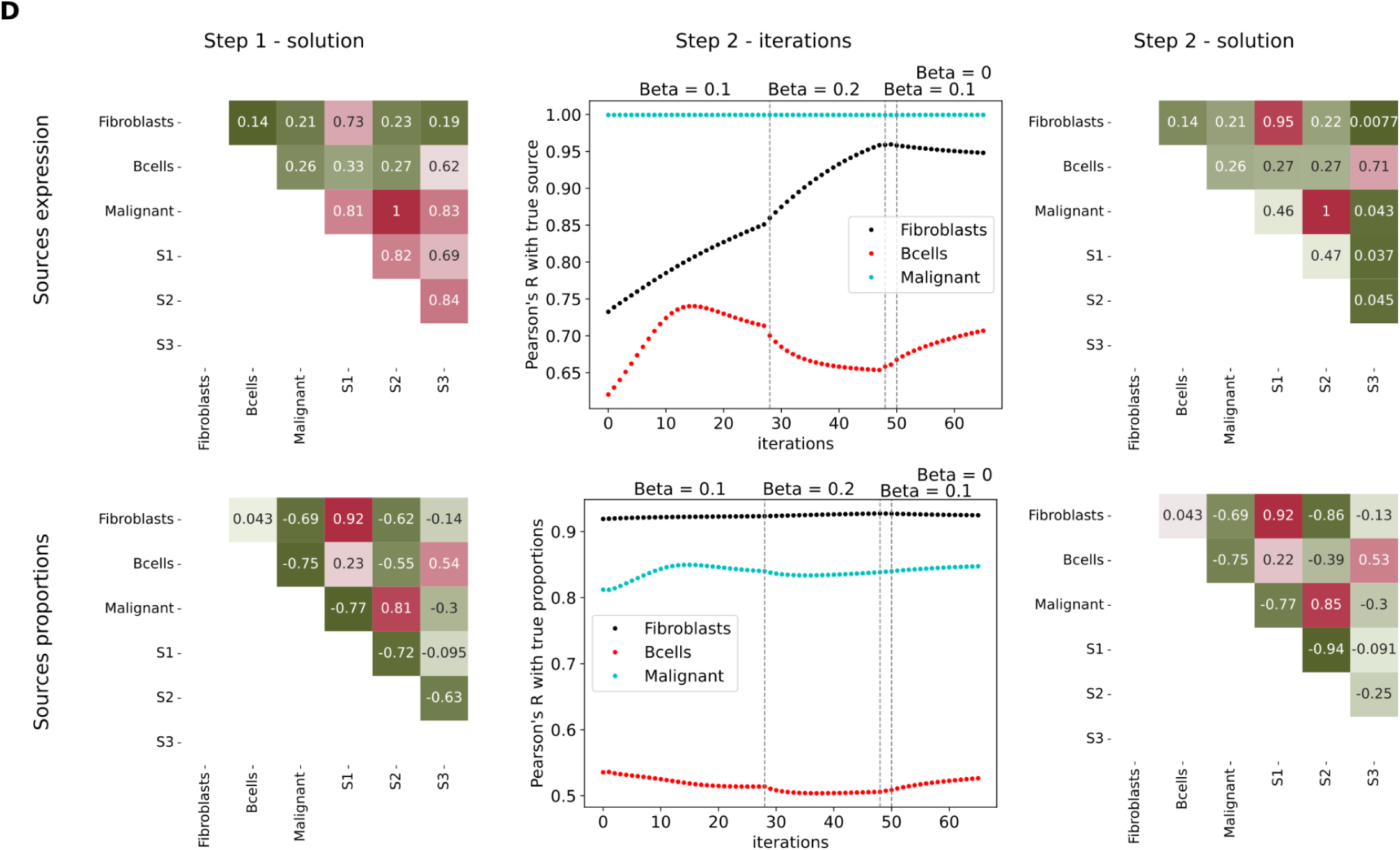
Pearson’s correlation between ground truth sources and proportions of the three cell types used to create Mix D samples. Correlations between output from the first and second steps CDState are compared. Values of Beta used in the second step are reported on the middle plot.

**Suppl. Figure S2.**
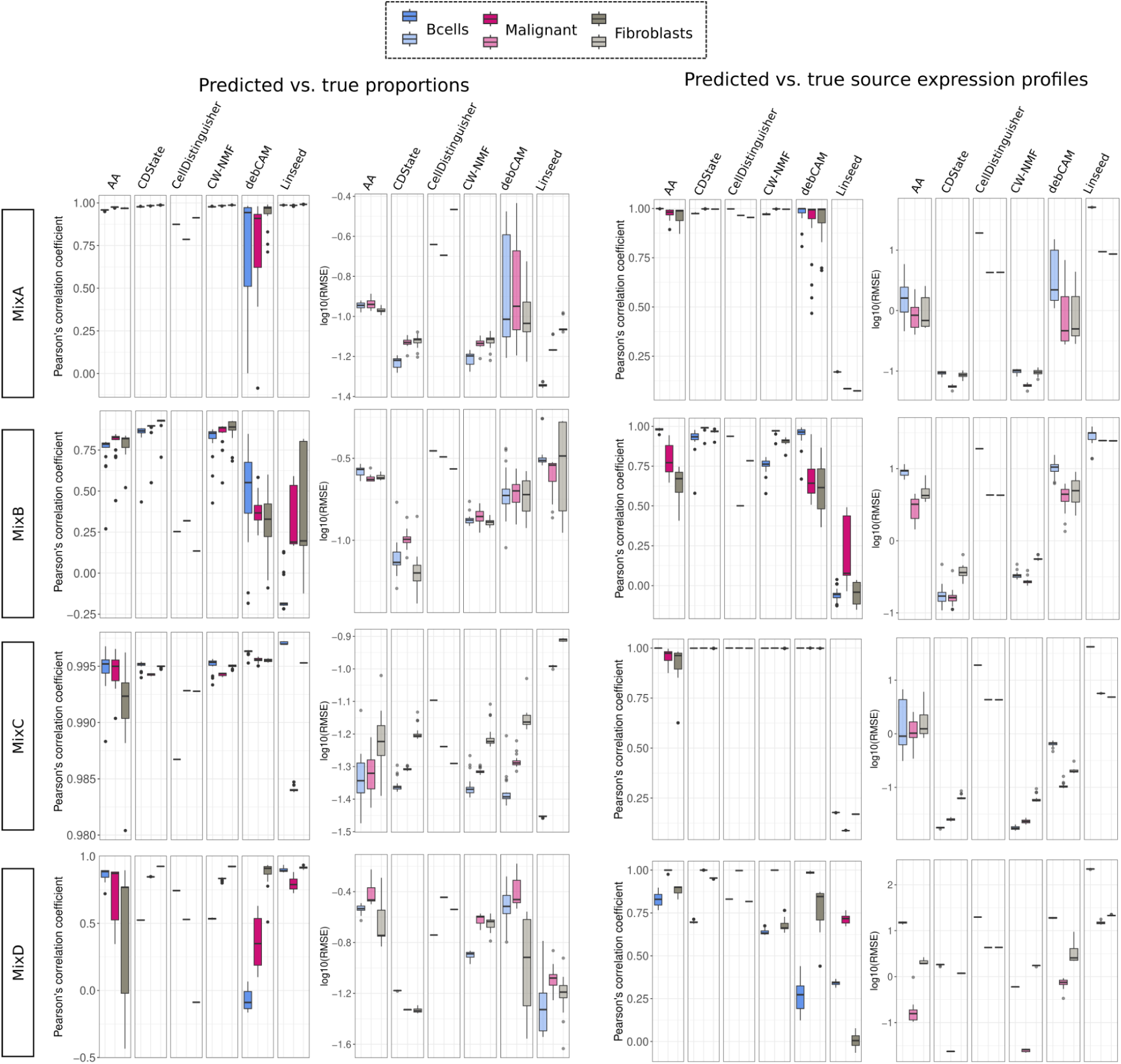
Performance comparison between CDState and existing unsupervised deconvolution methods: Archetype Analysis (AA), Linseed, CellDistinguisher, debCAM, and output from the first step of CDState (CW-NMF). Barplots represent Pearson’s correlation coefficient and root mean squared error (RMSE) values from comparing inferred and true source gene expression and proportions of B-cells, Malignant cells and Fibroblasts from *Kim. et al.* single cell dataset.

**Suppl. Figure S3.**
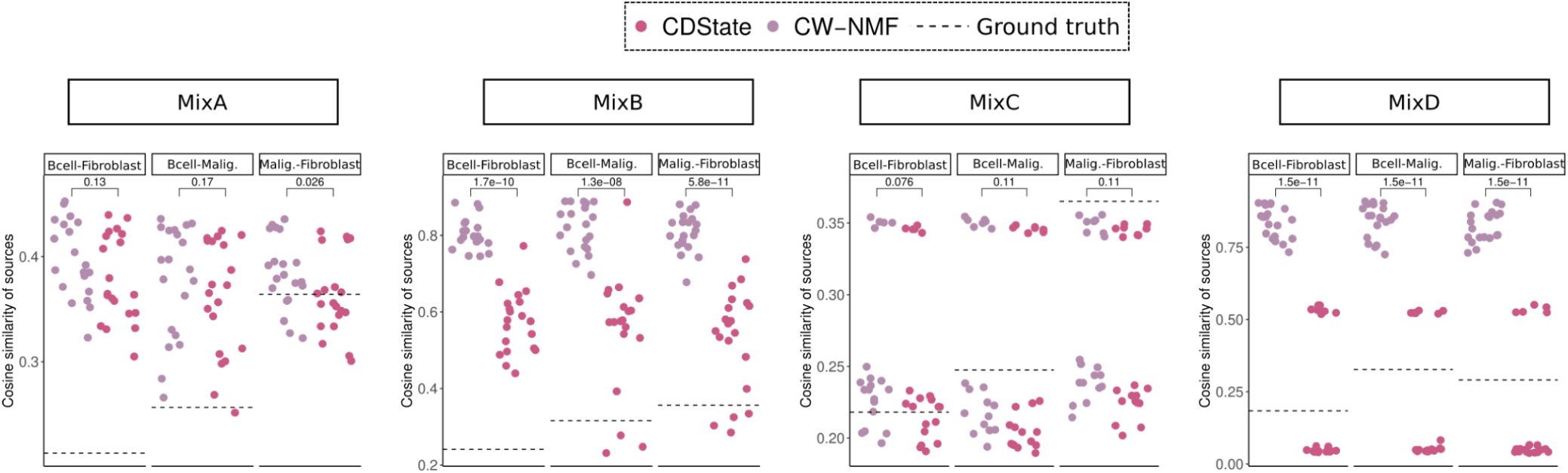
Comparison of pairwise cosine similarity between gene expression vectors of B-cells, Fibroblasts, and Malignant cells between inferred source gene expression after the first step of CDState (CW-NMF) and the second step (CDState). Dotted lines represent cosine similarity between ground truth gene expression vectors.

**Suppl. Figure S4.**
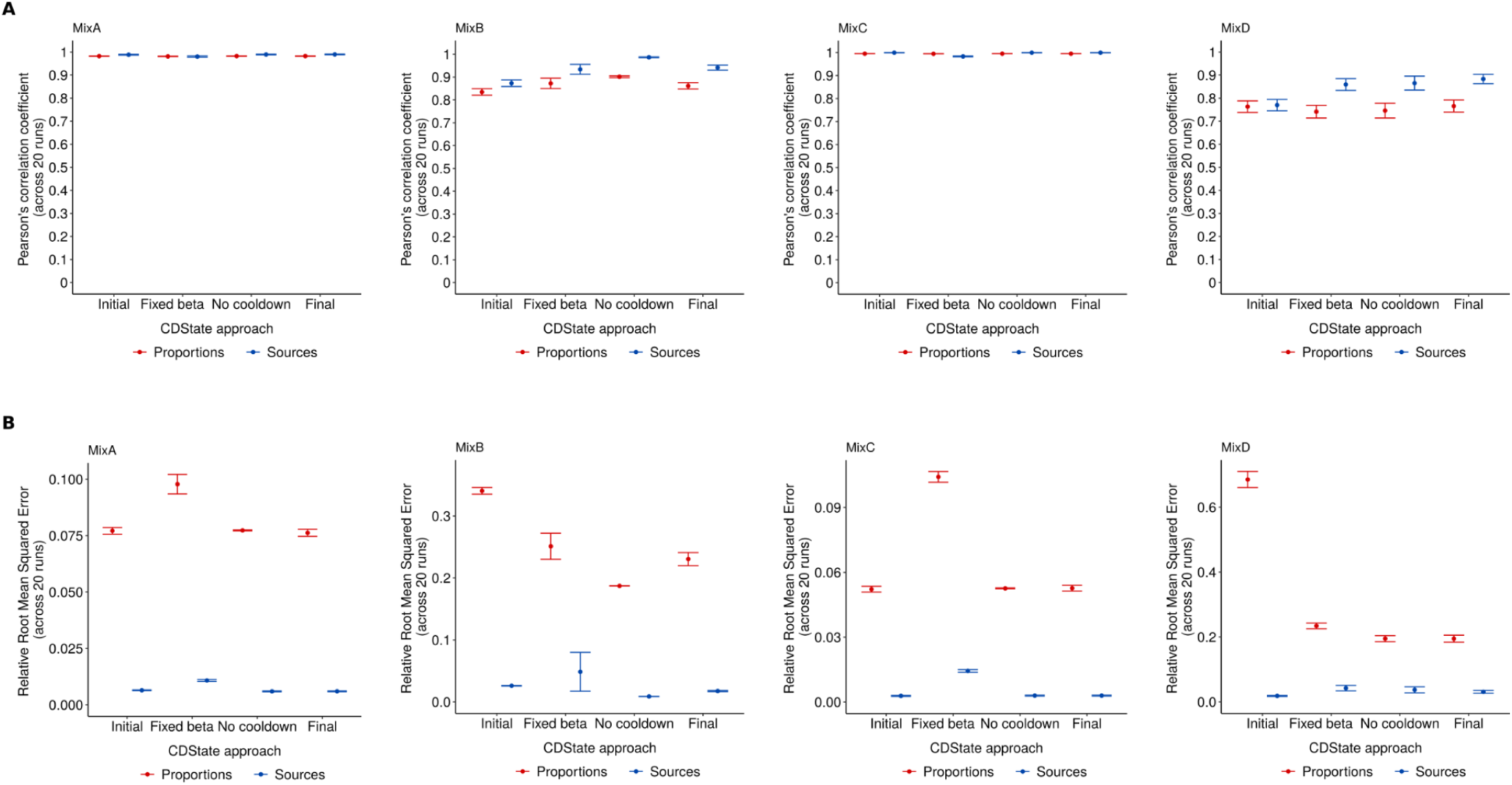
Comparison of CDState performance using solution from the initial CDState phase (Initial), solution from running CDState with fixed beta (Fixed beta), solution from running CDState without final decreasing of beta (No cooldown), and final CDState solution (Final). Mean Pearson’s correlation coefficient **(A)** and mean relative root mean squared error **(B)** calculated for estimated and true proportions and sources across 20 runs. Error bars represent standard error.

**Suppl. Figure S5.**
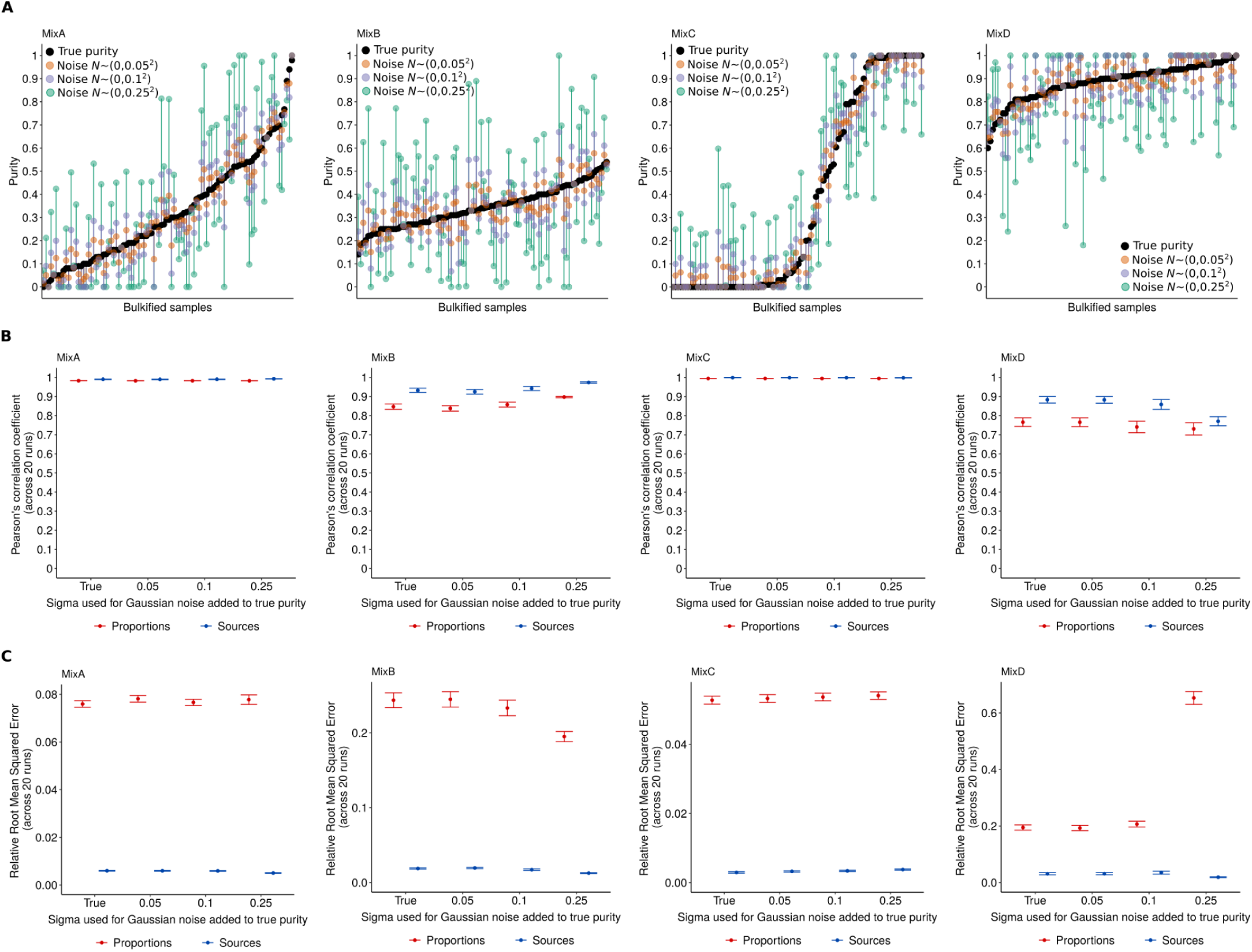
CDState robustness against noisy purity estimated. **A.** True and noisy purity estimates used for validating CDState performance. For each of the four mixtures, 3 noisy purity sets were generated by adding Gaussian noise with varied sigma and clipping the resulting values to range [0,1]. Mean Pearson’s correlation coefficient **(A)** and mean relative root mean squared error **(B)** calculated for estimated and true proportions and sources across 20 runs and the four types of purity estimates. Error bars represent standard error.

**Suppl. Figure S6.**
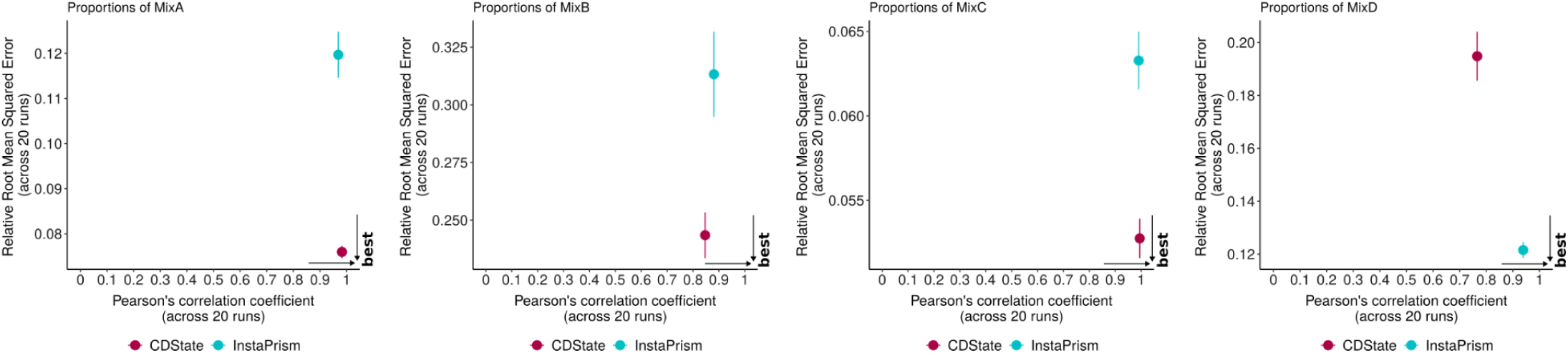
Comparison of CDState and InstaPrism performance in proportion estimation across the four mixtures. Error bars represent standard error.

**Suppl. Figure S7.**
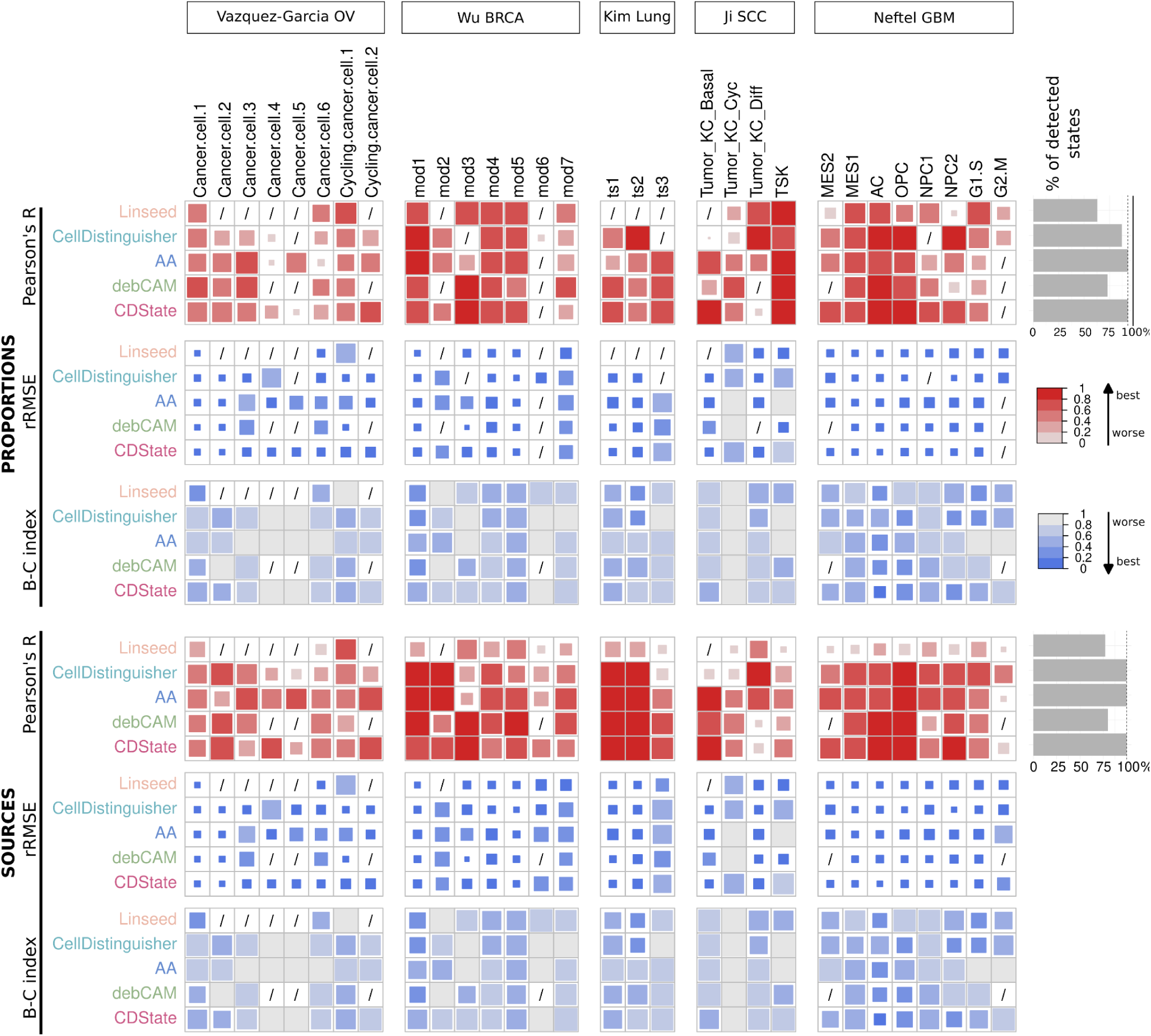
Performance comparison in detection of malignant cell states reported in the original studies. Heatmaps represent methods’ performance when comparing estimated and true proportions and source gene expression, by evaluating Pearson’s correlation coefficient, relative Root Mean Squared Error (rRMSE) and Bray-Curtis dissimilarity index (B-C index). Histograms report the percentage of detected states across the five datasets based on positive correlation with true proportions and source gene expression. / - state was not identified.

**Suppl. Figure S8.**
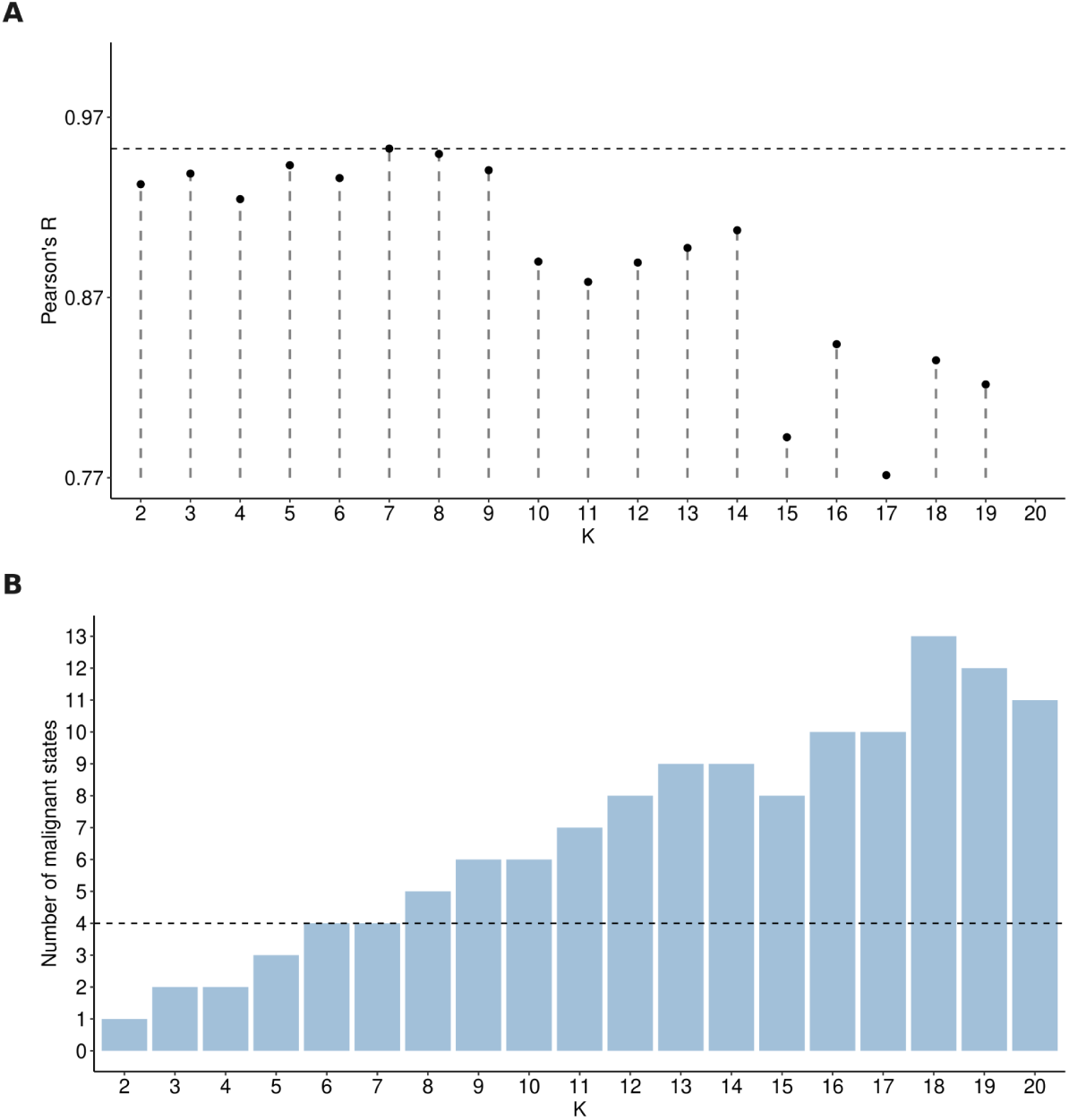
Selection of optimal number of cell states from CDState deconvolution of bulkified glioblastoma dataset. **A.** Correlation between CDState-inferred and true tumor purity across best CDState runs for a range of *k*. **B.** Number of malignant components identified in the run with the highest correlation between true and inferred tumor purity across a range of *k*.

**Suppl. Figure S9.**
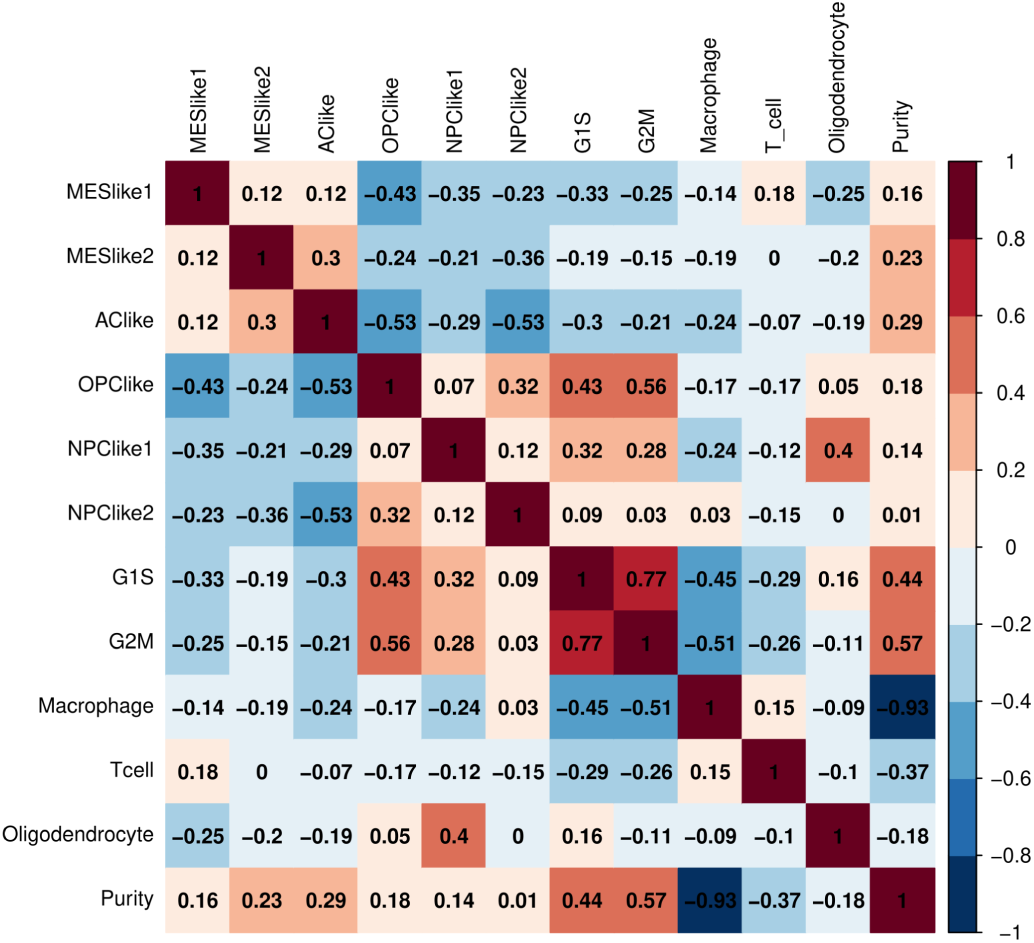
Correlation between true proportions of cells in the bulkified GBM *Neftel et al.* dataset. Numbers correspond to Pearson’s correlation coefficient.

**Suppl. Figure S10.**
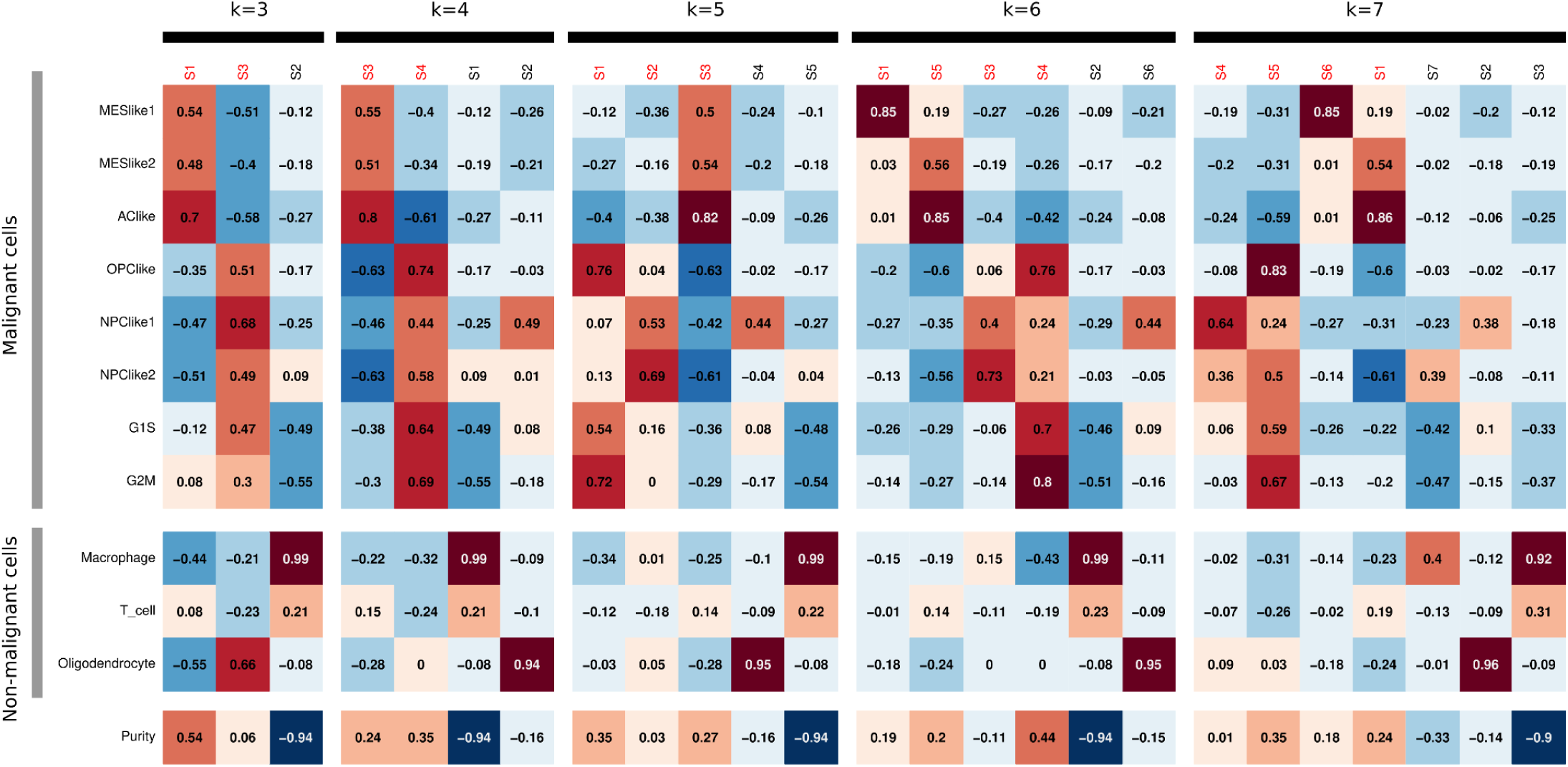
Correlation between true and CDState-identified cell state proportions of the glioblastoma dataset, across a range of *k* (3 to 7). Red font above heatmaps corresponds to malignant cell states.

**Suppl. Figure S11.**
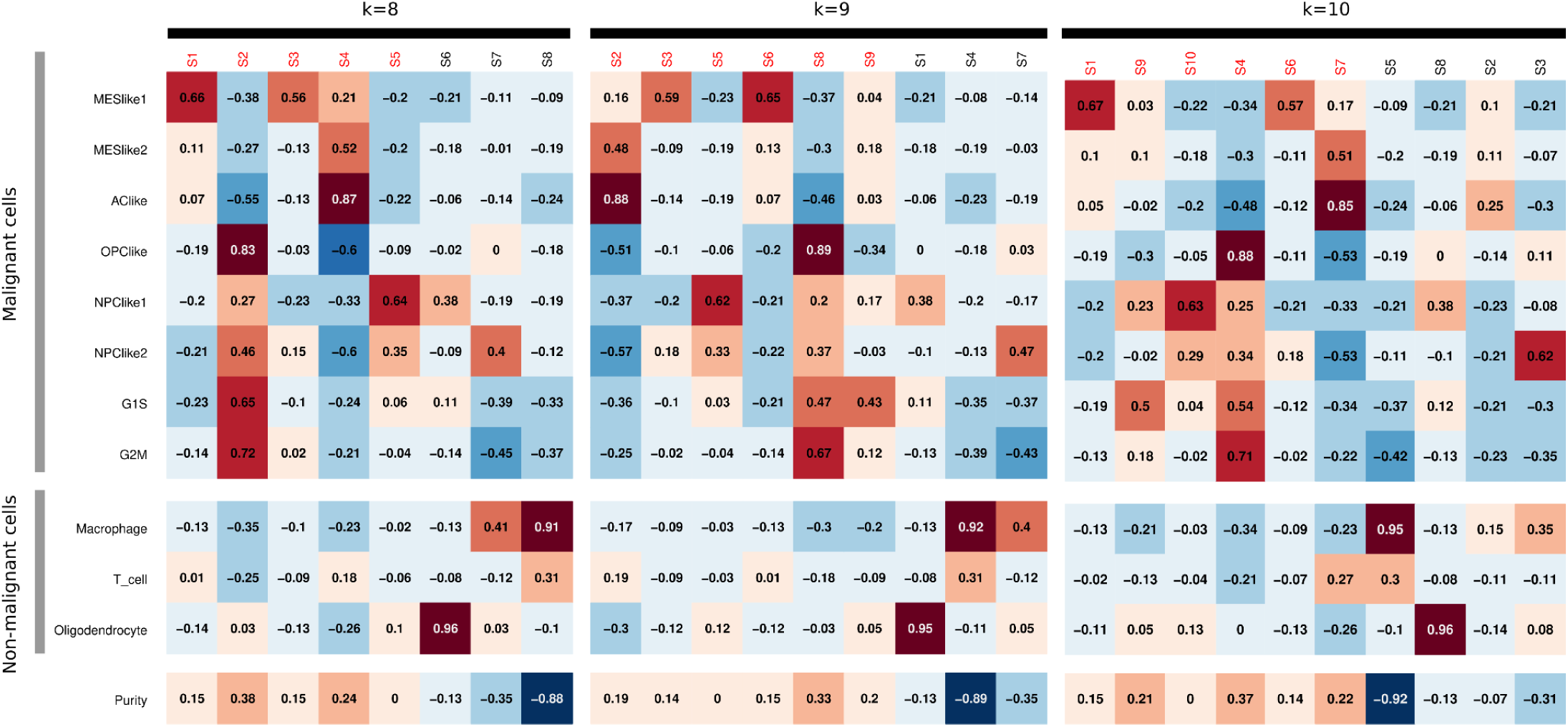
Correlation between true and CDState-identified cell state proportions of the glioblastoma dataset, across a range of *k* (8 to 10). Red font above heatmaps corresponds to malignant cell states.

**Suppl. Figure S12.**
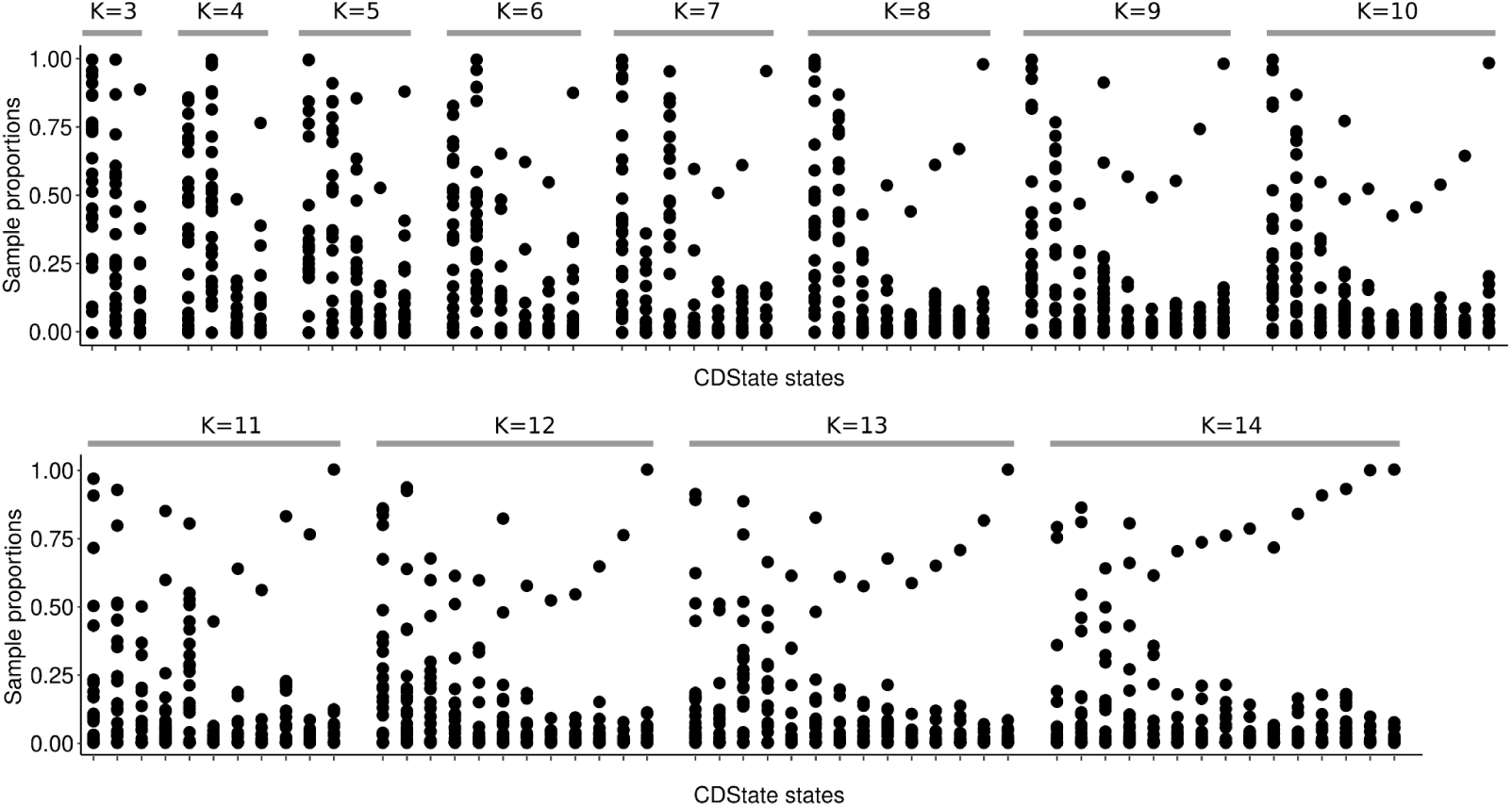
Distribution of CDState-identified malignant cell state proportions in bulkified glioblastoma data.

**Suppl. Figure S13.**
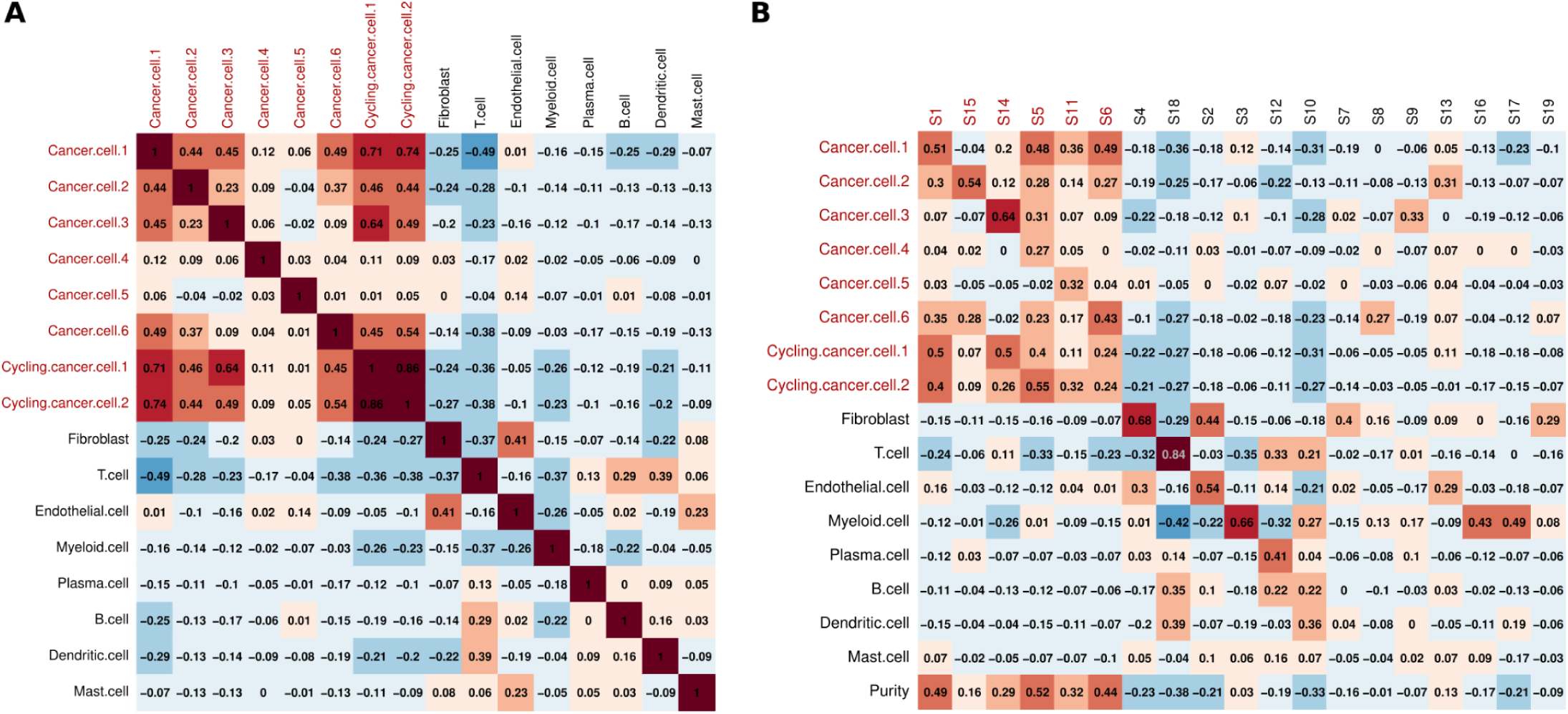
Correlation between true proportions of cells **(A)** and true and CDState-identified state proportions **(B)** in the bulkified OV *Vazquez-Garcia et al.* dataset. Numbers correspond to Pearson’s correlation coefficient.

**Suppl. Figure S14.**
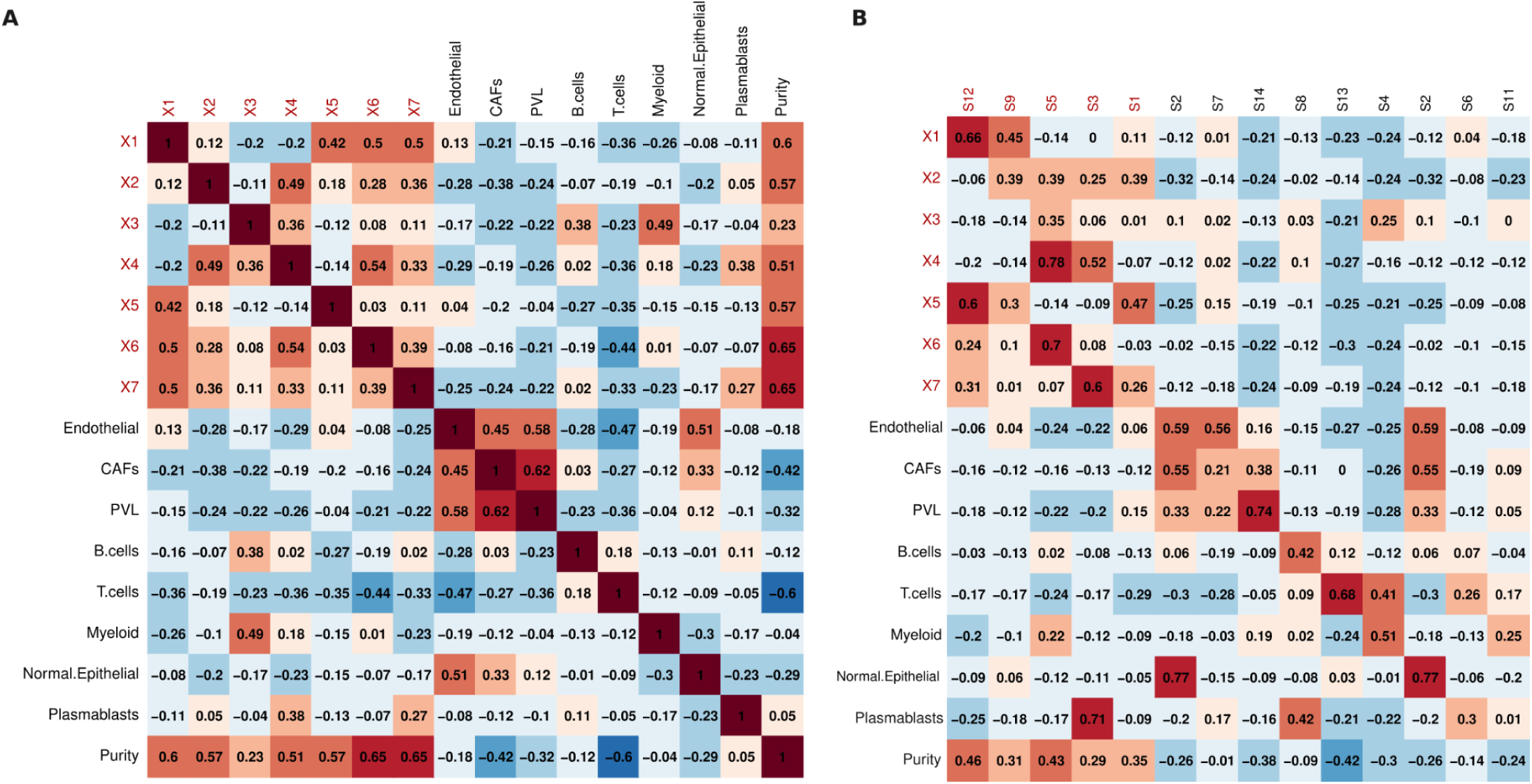
Correlation between true proportions of cells **(A)** and true and CDState-identified state proportions **(B)** in the bulkified BRCA *Wu et al.* dataset. Numbers correspond to Pearson’s correlation coefficient.

**Suppl. Figure S15.**
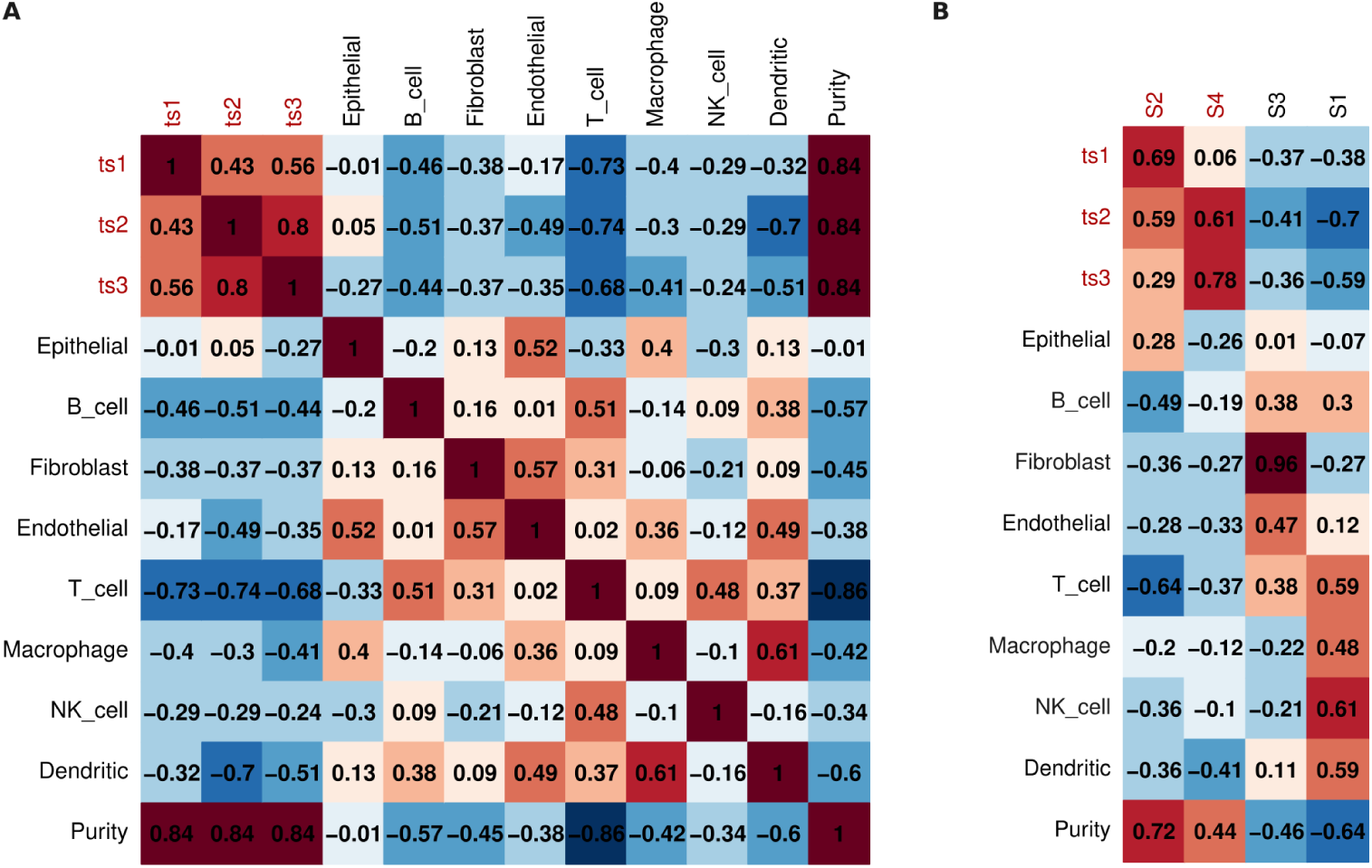
Correlation between true proportions of cells **(A)** and true and CDState-identified state proportions **(B)** in the bulkified LUAD *Kim et al.* dataset. Numbers correspond to Pearson’s correlation coefficient.

**Suppl. Figure S16.**
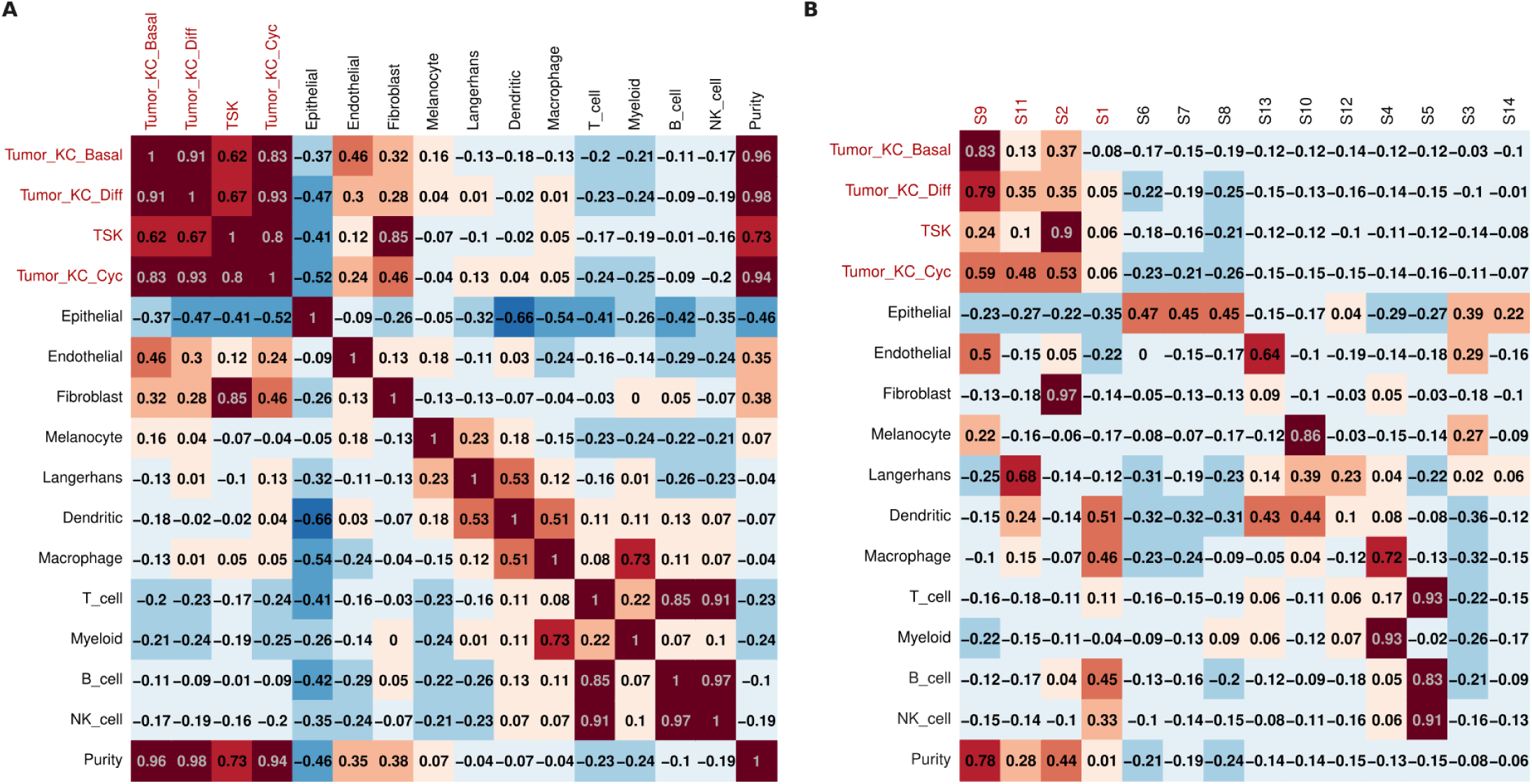
Correlation between true proportions of cells **(A)** and true and CDState-identified state proportions **(B)** in the bulkified SCC *Ji et al.* dataset. Numbers correspond to Pearson’s correlation coefficient.

**Suppl. Figure S17.**
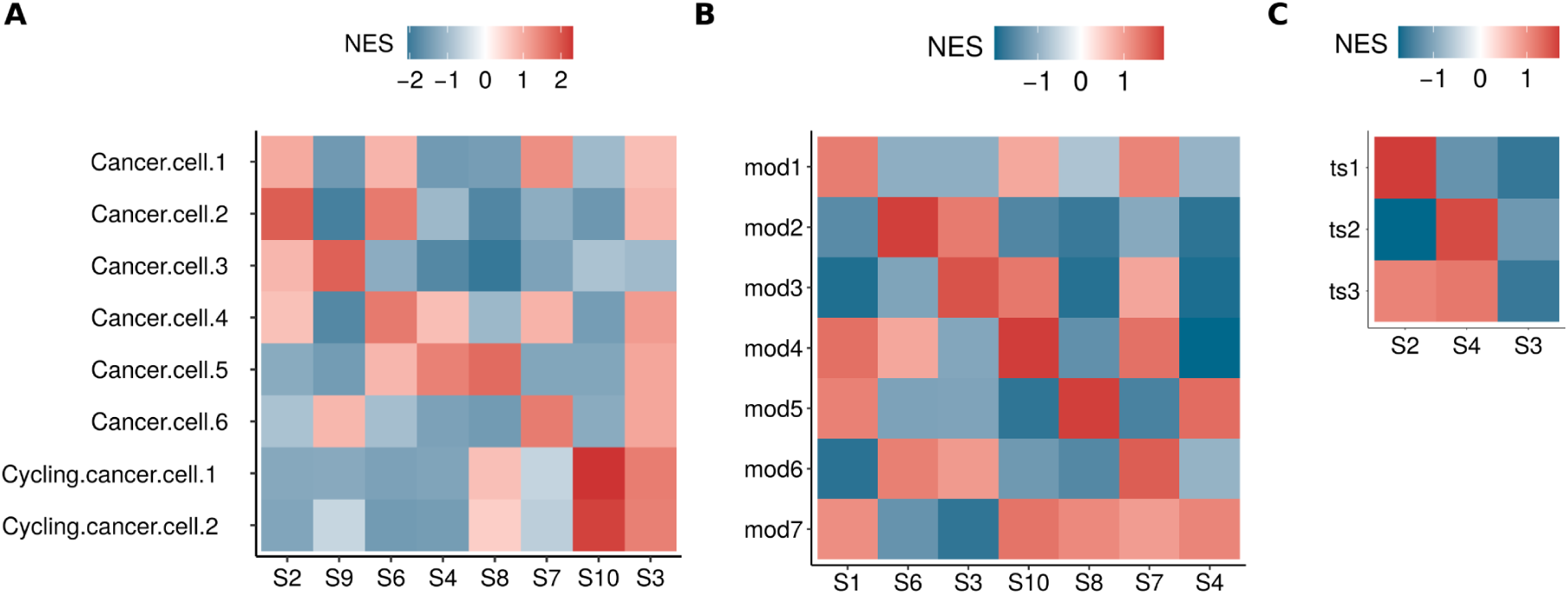
Normalized enrichment score of known malignant cell state signatures in true bulk datasets from ovarian carcinoma [40], breast cancer [41], and lung adenocarcinoma [42].

**Suppl. Figure S18.**
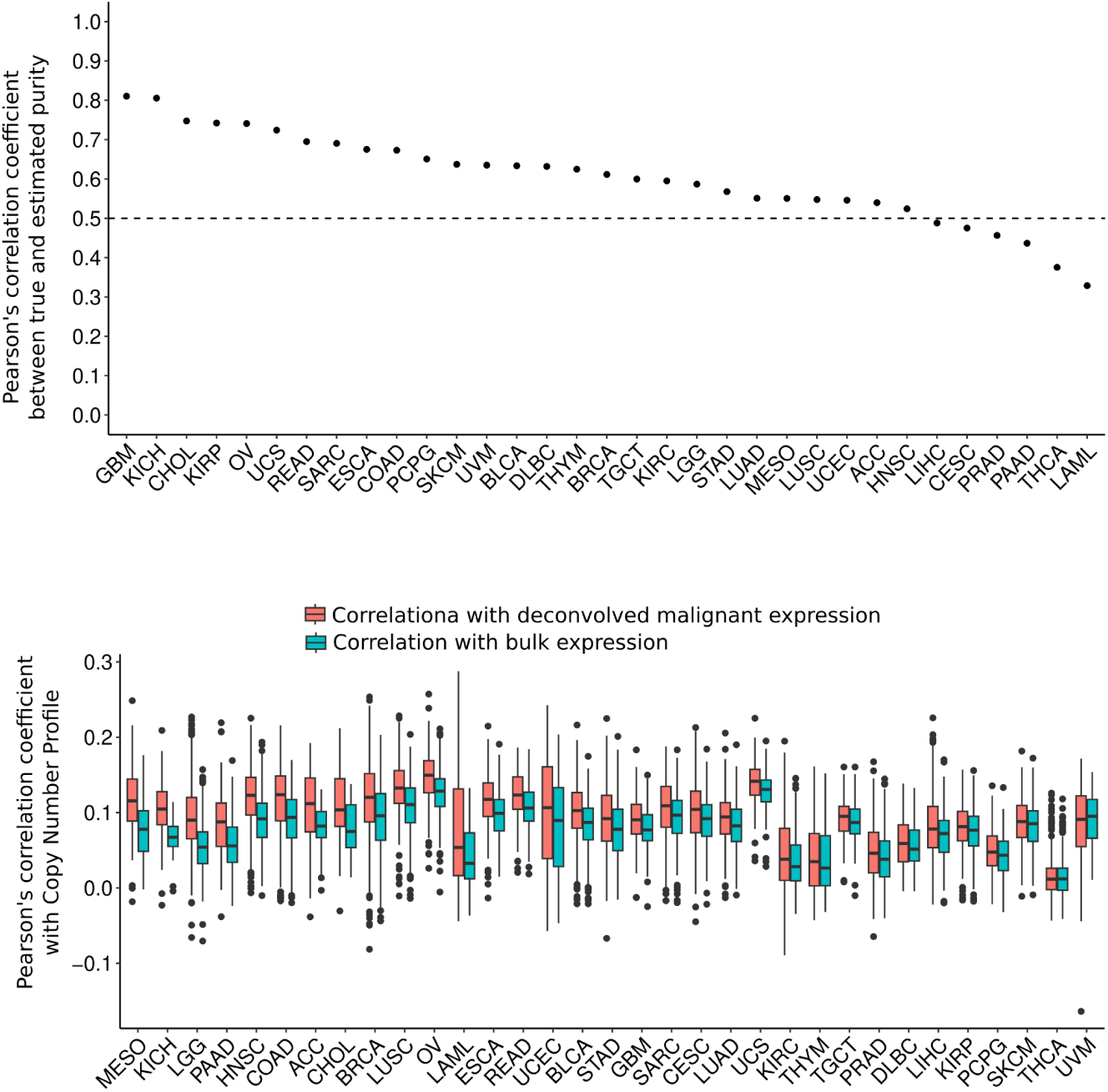
Deconvolution of TCGA datasets with CDState. Pearson’s correlation between CDState-inferred and true tumor purity of TCGA datasets (top). Comparison of correlation between gene copy number profile and deconvolved malignant (blue) or bulk (pink) gene expression across TCGA datasets (bottom). The deconvolved malignant expression was calculated per sample, by subtracting CDState-inferred tumor-microenvironment source expression multiplied by corresponding inferred proportions from the bulk RNA-seq data.

**Suppl. Figure S19.**
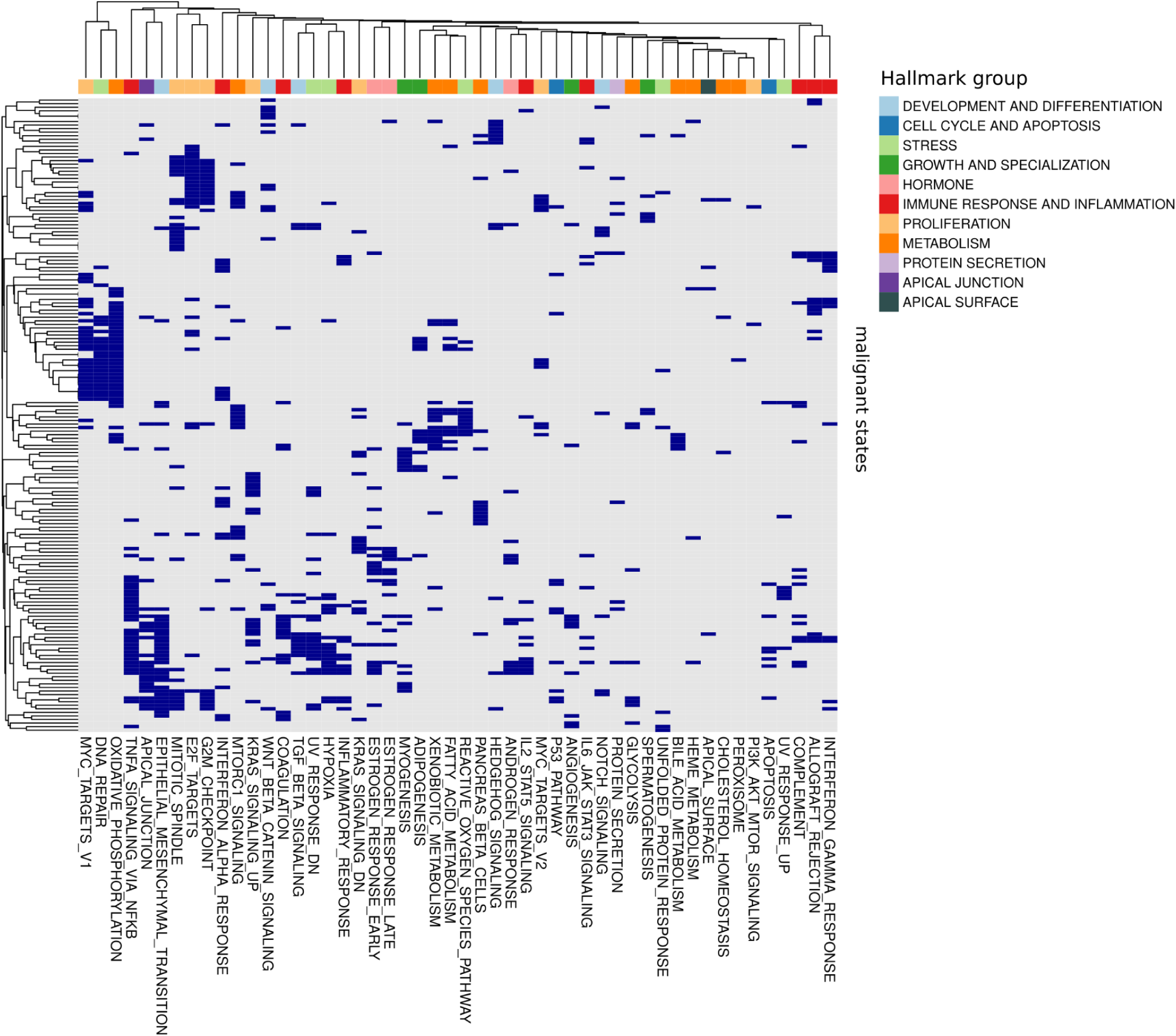
Clustering of hallmark gene programs expressed across TCGA malignant cell states. States are annotated with the group of the most enriched hallmark set.

**Suppl. Figure S20.**
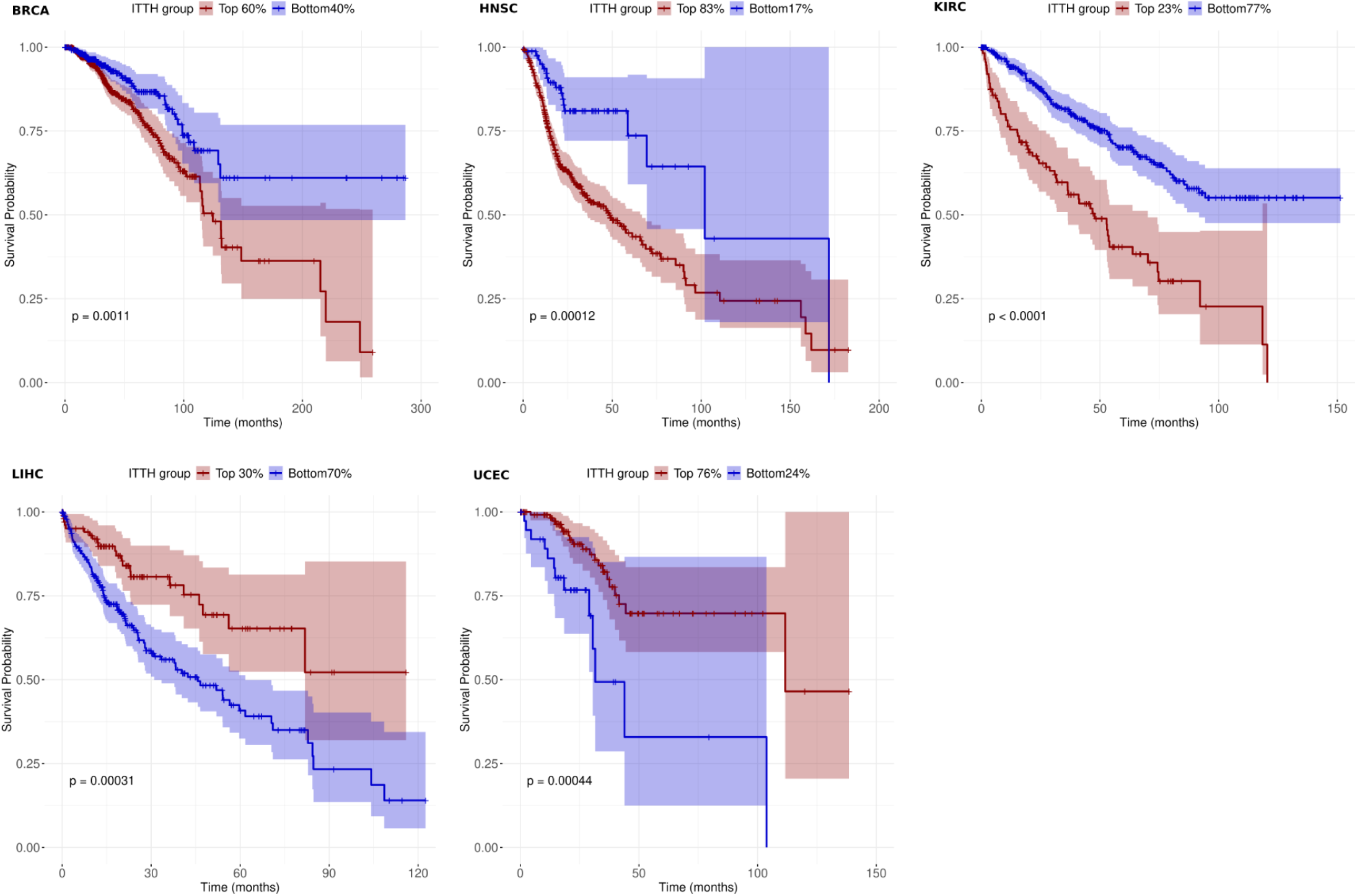
Kaplan-Meier plots comparing patient survival probability between groups with high and low ITTH index in five TCGA cancer types. Log-rank test p-value is reported for each comparison.

**Suppl. Figure S21.**
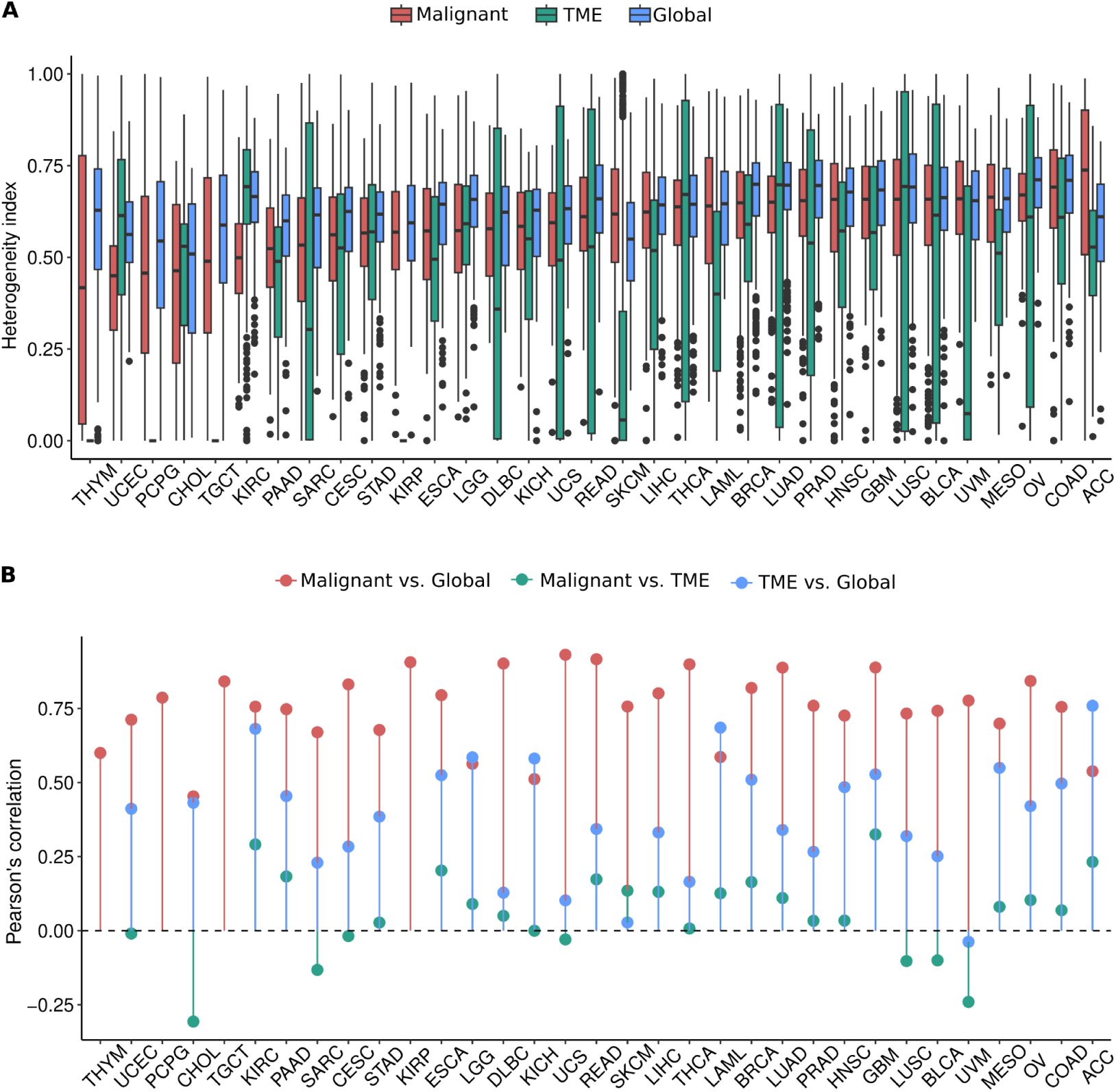
Comparison of ITTH indexes across TCGA cancer types. **A.** Distribution of malignant, non-malignant (TME) and global ITTH indexes, calculated using source proportions of malignant, non-malignant and all CDState-deconvolved states, respectively. **B.** Correlation between malignant, non-malignant and global ITTH indexes.

**Suppl. Figure S22.**
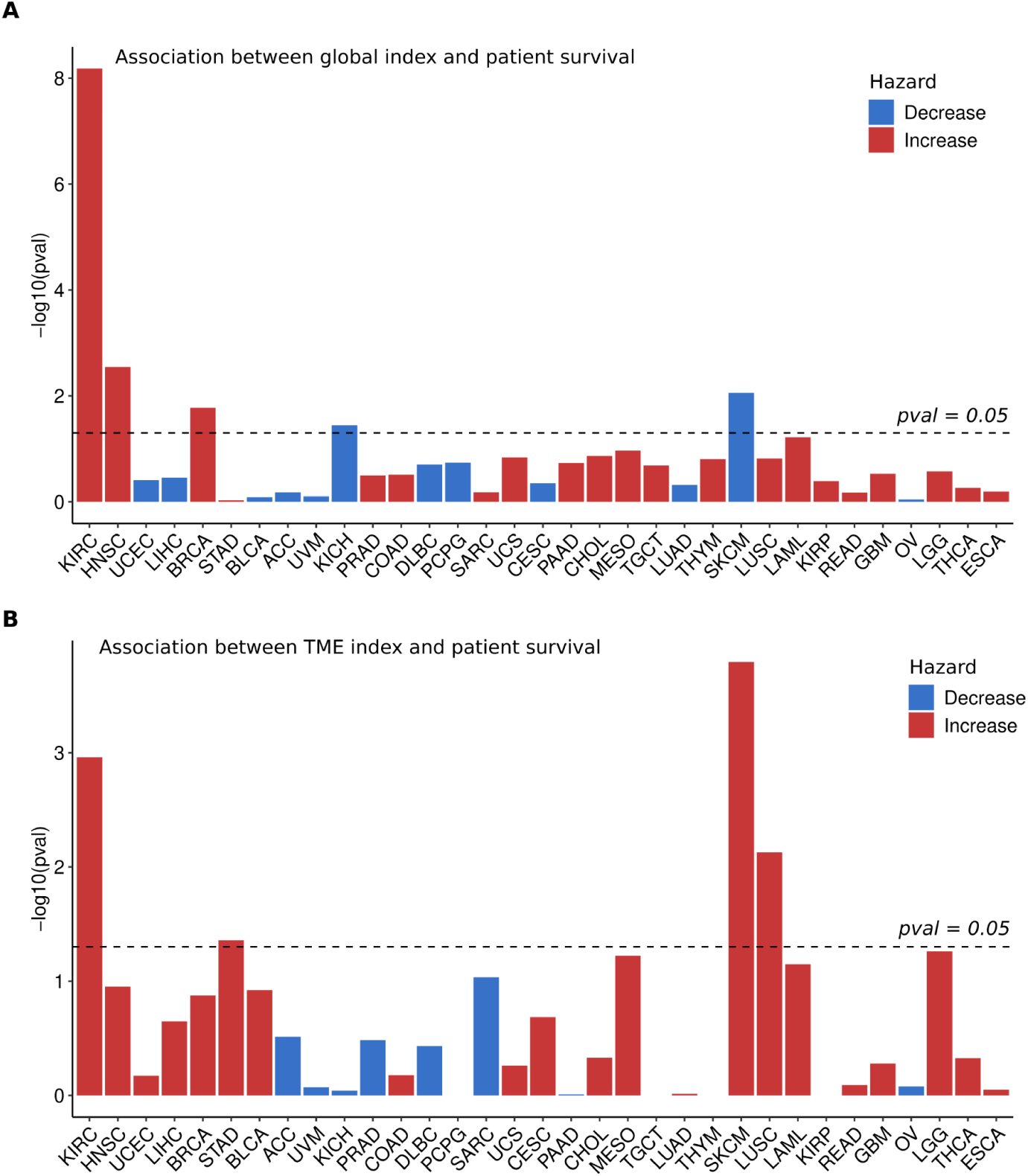
Association between global **(A)** and non-malignant **(B)** ITTH indexes and patient prognosis. Cox-regression p-value is reported on y-axis.

**Suppl. Figure S23.**
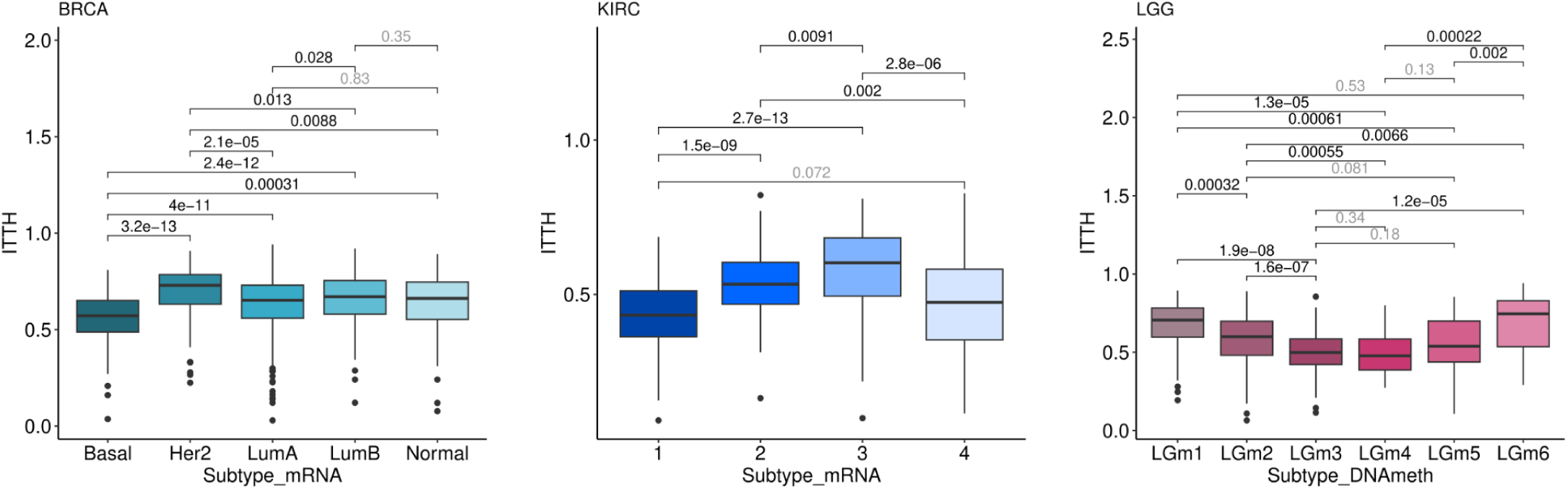
Distribution of ITTH index across mRNA subtype of TCGA BRCA and KIRC samples, as well as across DNA methylation subtype of TCGA LGG samples. Significance computed using Mann-Whitney U test.

**Suppl. Figure S24.**
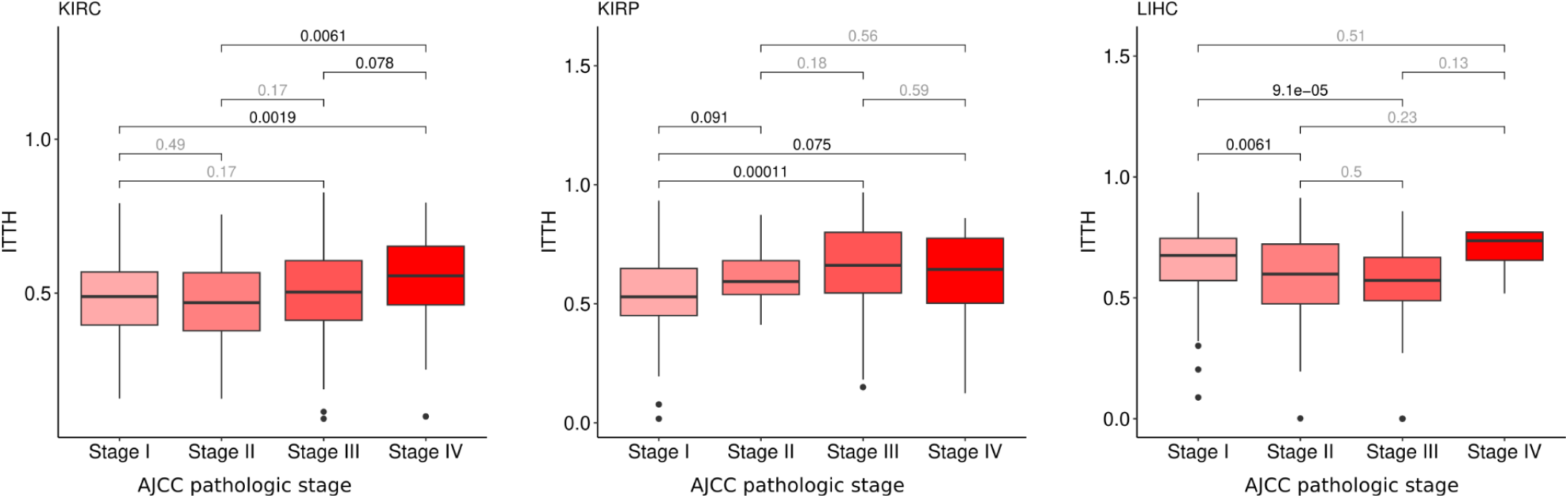
Distribution of ITTH index across AJCC tumor stage in TCGA KIRC, KIRP and LIHC samples. Significance computed using Mann-Whitney U test.

**Suppl. Figure S25.**
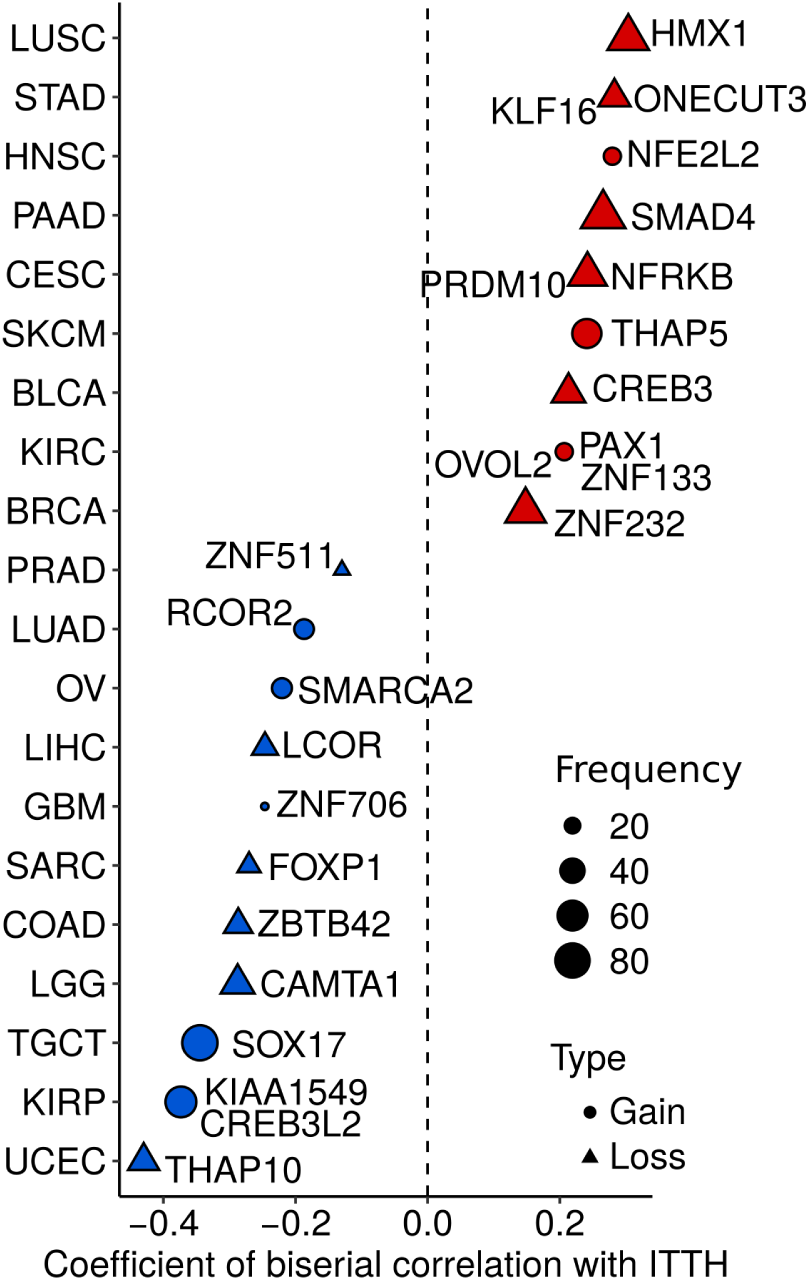
Transcriptional factors whose copy number alterations have the highest absolute biserial correlation (adjusted p-value < 0.05) with cancer-type specific ITTH score. Cancer-specific frequency of event is represented by size of points, while shape represents type of event.

**Suppl. Figure S26.**
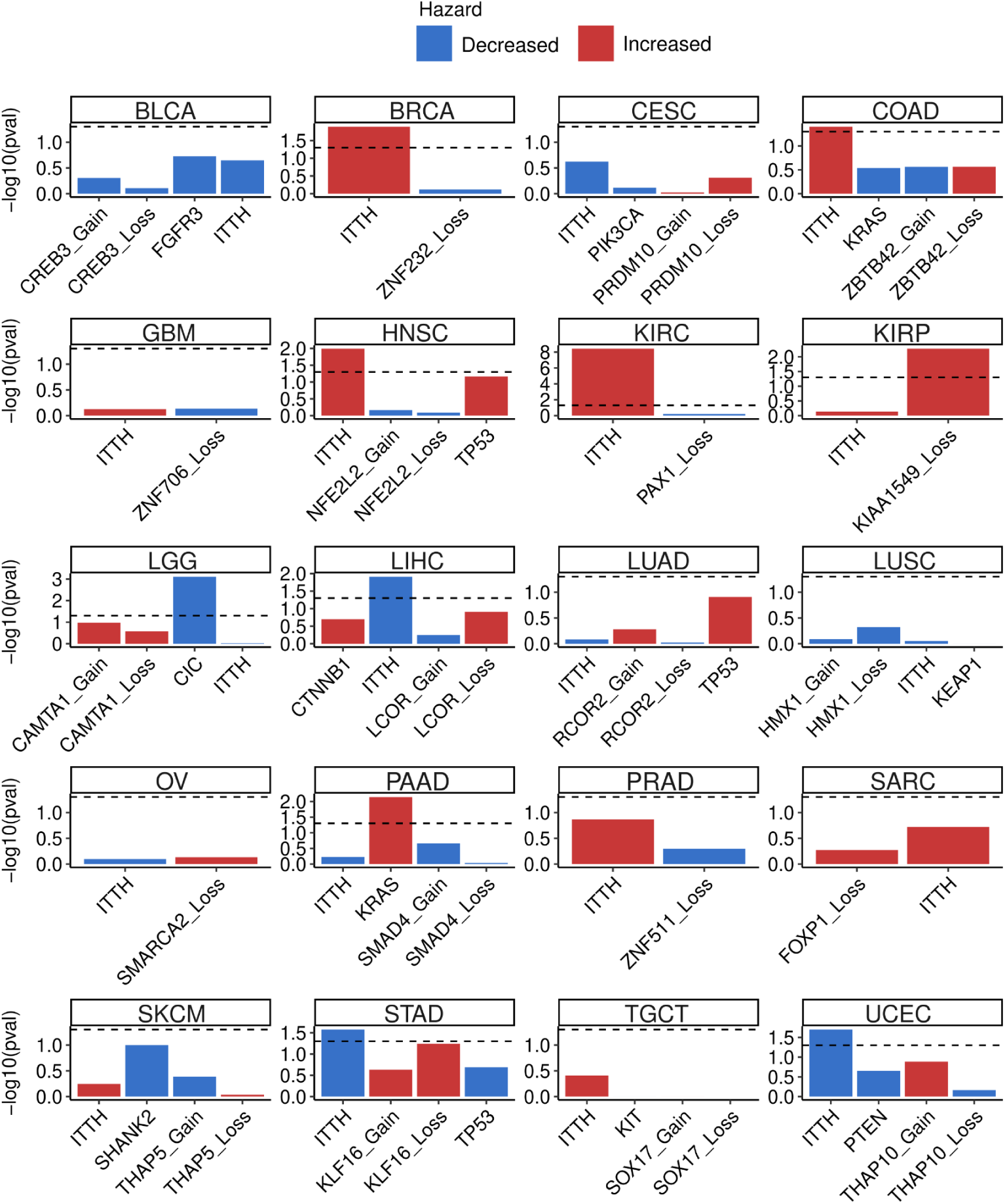
Association between malignant ITTH scores and their associated genetic events with patient survival. Feature-specific p-value obtained from multivariate Cox-regression models is reported on y-axis. Dotted horizontal line corresponds to p-value = 0.05.

**Suppl. Figure S27.**
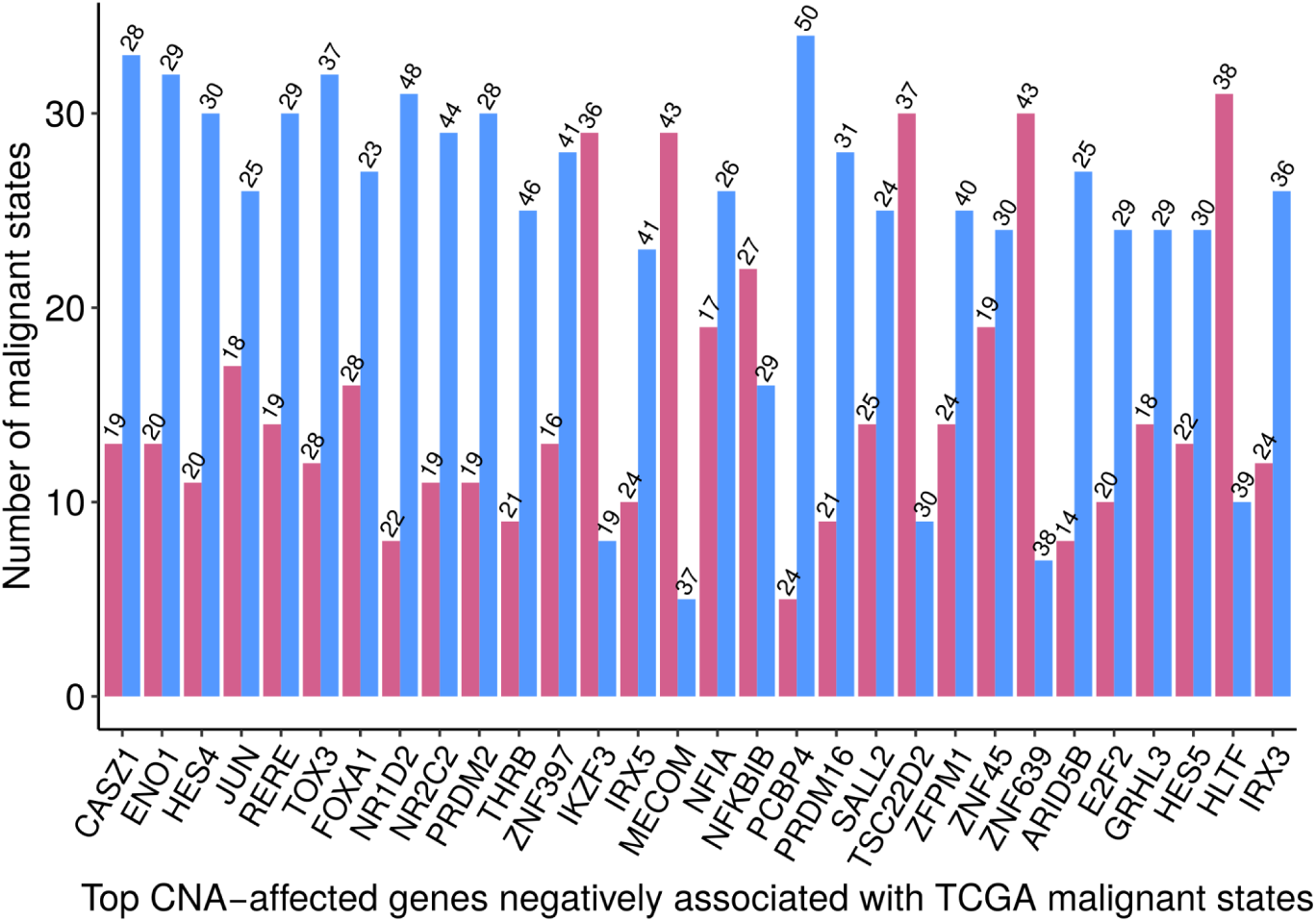
Top 30 most frequent transcriptional factors whose copy number alterations are negatively associated with TCGA malignant cell state proportions (adjusted p-value < 0.05). Numbers on top of bar plots represent the mean frequency [%] of events across all analyzed TCGA samples. Blue and red bars represent negative and positive associations, respectively.

**Suppl. Figure S28.**
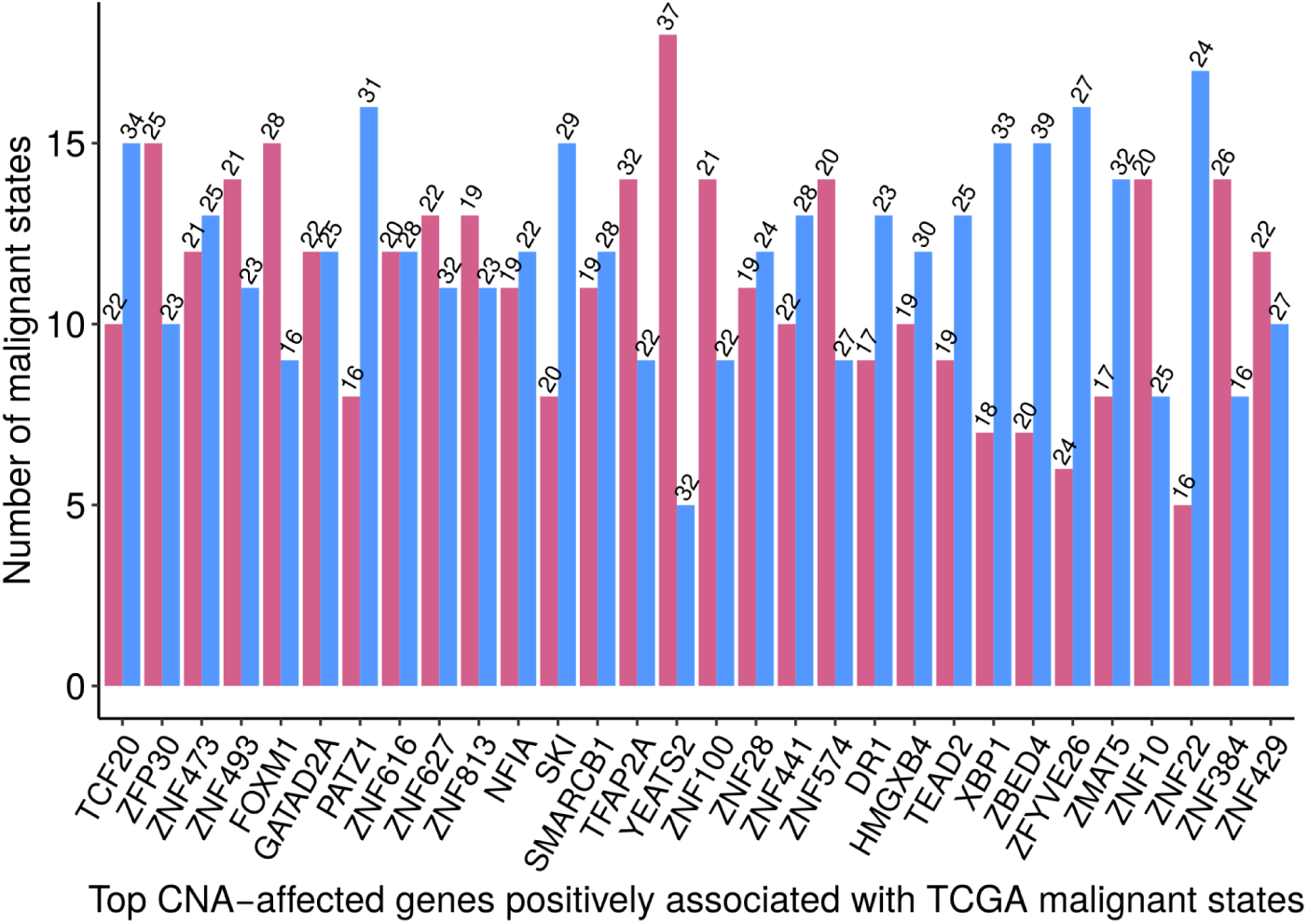
Top 30 most frequent transcriptional factors whose copy number alterations are positively associated with TCGA malignant cell state proportions (adjusted p-value < 0.05). Numbers on top of bar plots represent the mean frequency [%] of events across all analyzed TCGA samples. Blue and red bars represent negative and positive associations, respectively.

**Suppl. Figure S29.**
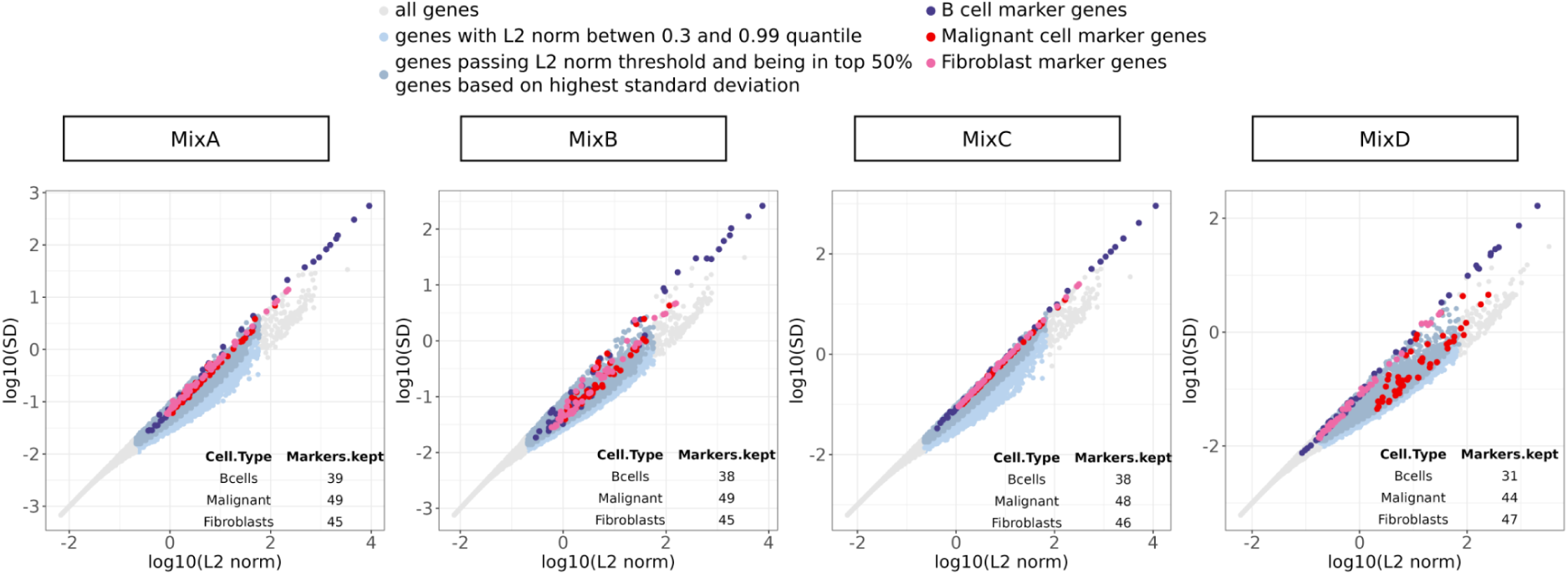
Overall magnitude of expression and standard deviation of all and filtered genes for deconvolution with CDState. Marker genes of B-cells, Fibroblasts and Malignant cells from *Kim et al.* lung adenocarcinoma single-cell data are colored with blue, pink and red, respectively. For the three cell types, 50 top marker genes were selected (Methods) using true gene expression of these cell types. The numbers in tables report the number of marker genes kept after applying CDState gene filtering procedure before deconvolution.

**Suppl. Figure S30.**
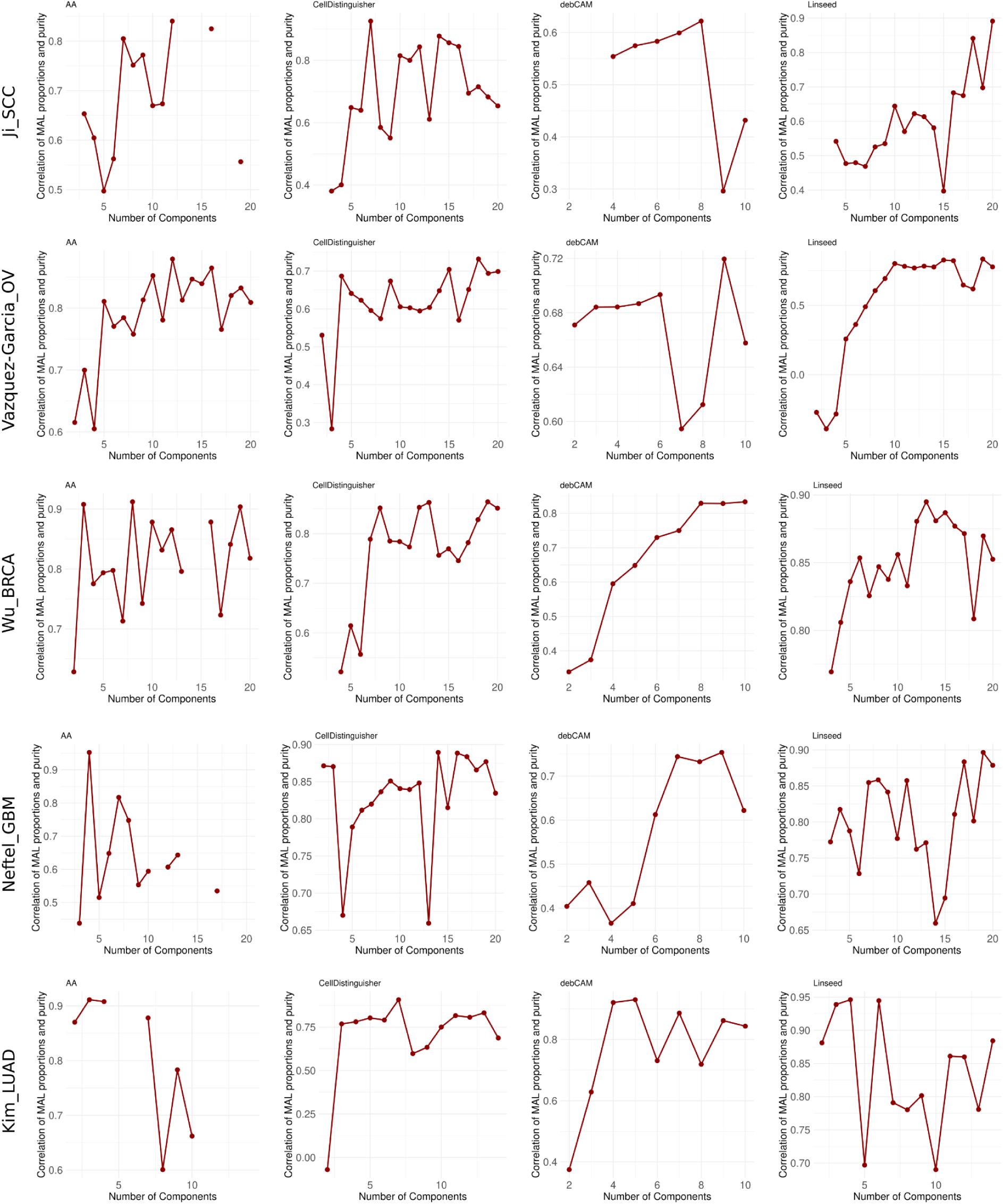
Correlation between estimated and true purity across a range of *k* and bulkified single-cell cancer datasets, inferred by existing deconvolution methods.

**Suppl. Figure S31.**
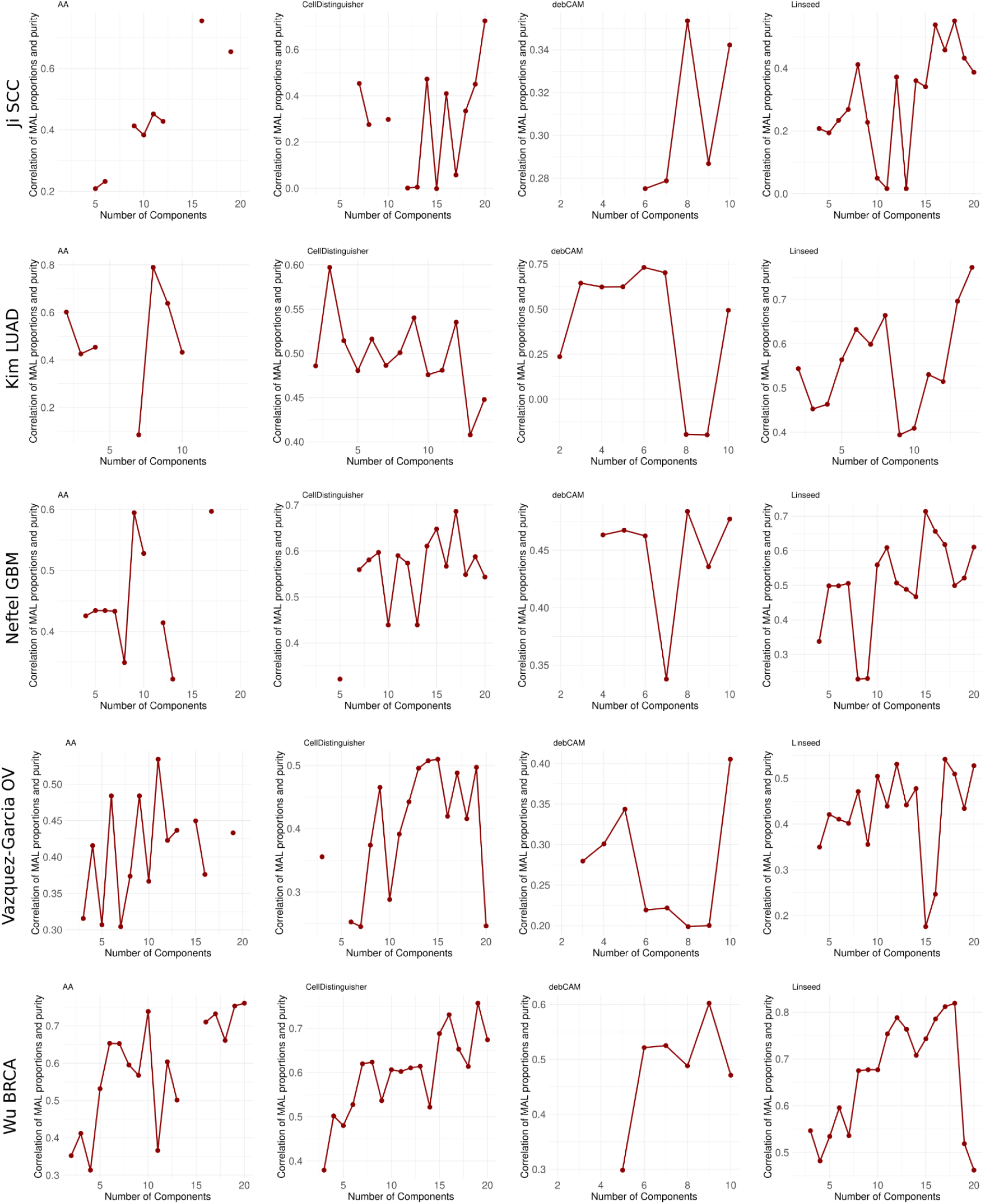
Correlation between estimated and purity with added noise across range of *k* and bulkified single-cell cancer datasets, inferred by existing deconvolution methods.

**Suppl. Figure S32.**
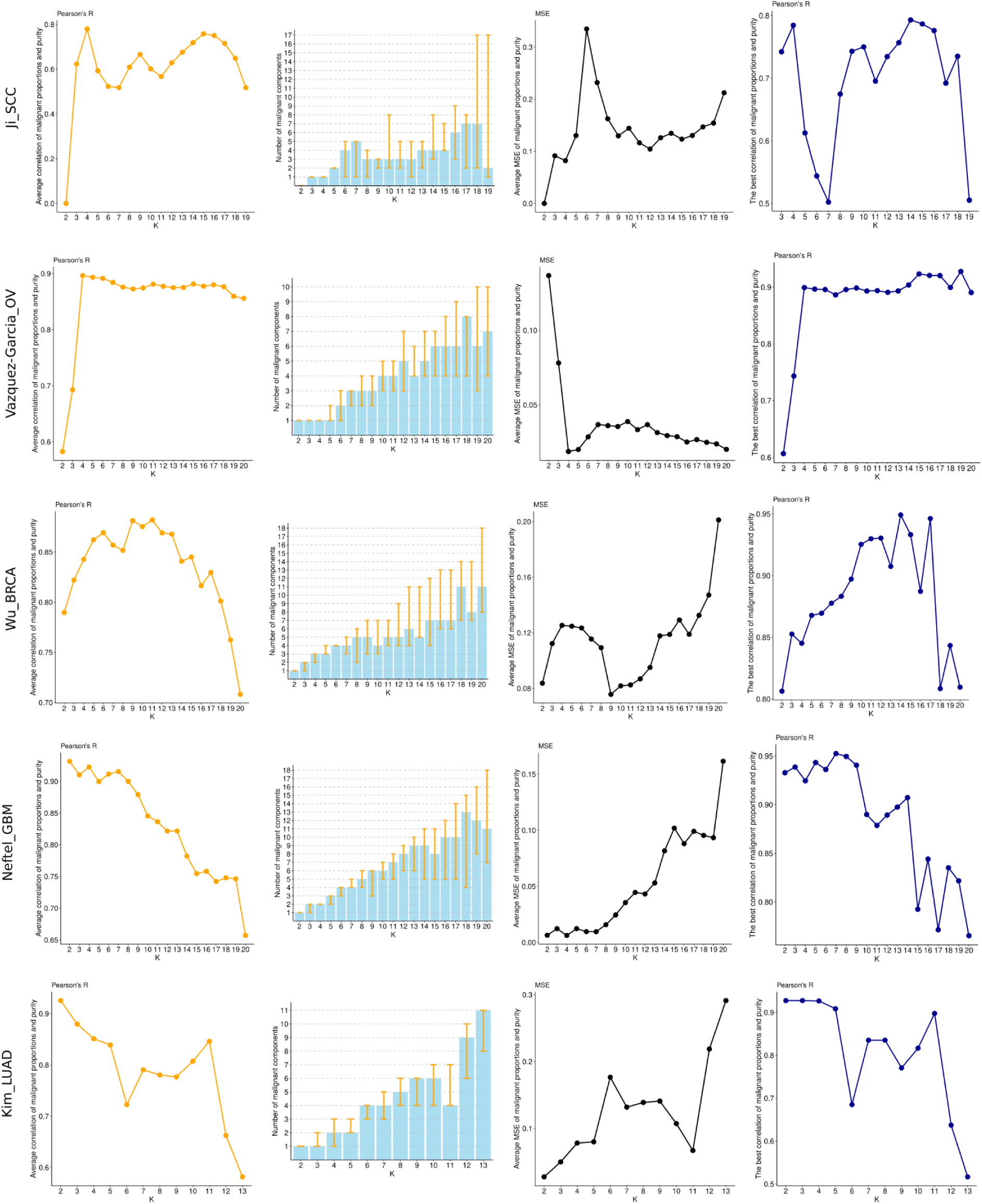
Average correlation and average mean squared error (MSE) between estimated and true purity along with mode number of detected malignant cell states for the best run out of 20 runs for a range of *k* and bulkified single-cell cancer datasets, inferred by CDState. Error bars represent the smallest and the highest number of detected malignant cell states for a given *k*. Top right column presents correlation with purity from the best selected run for each *k*.

**Suppl. Figure S33.**
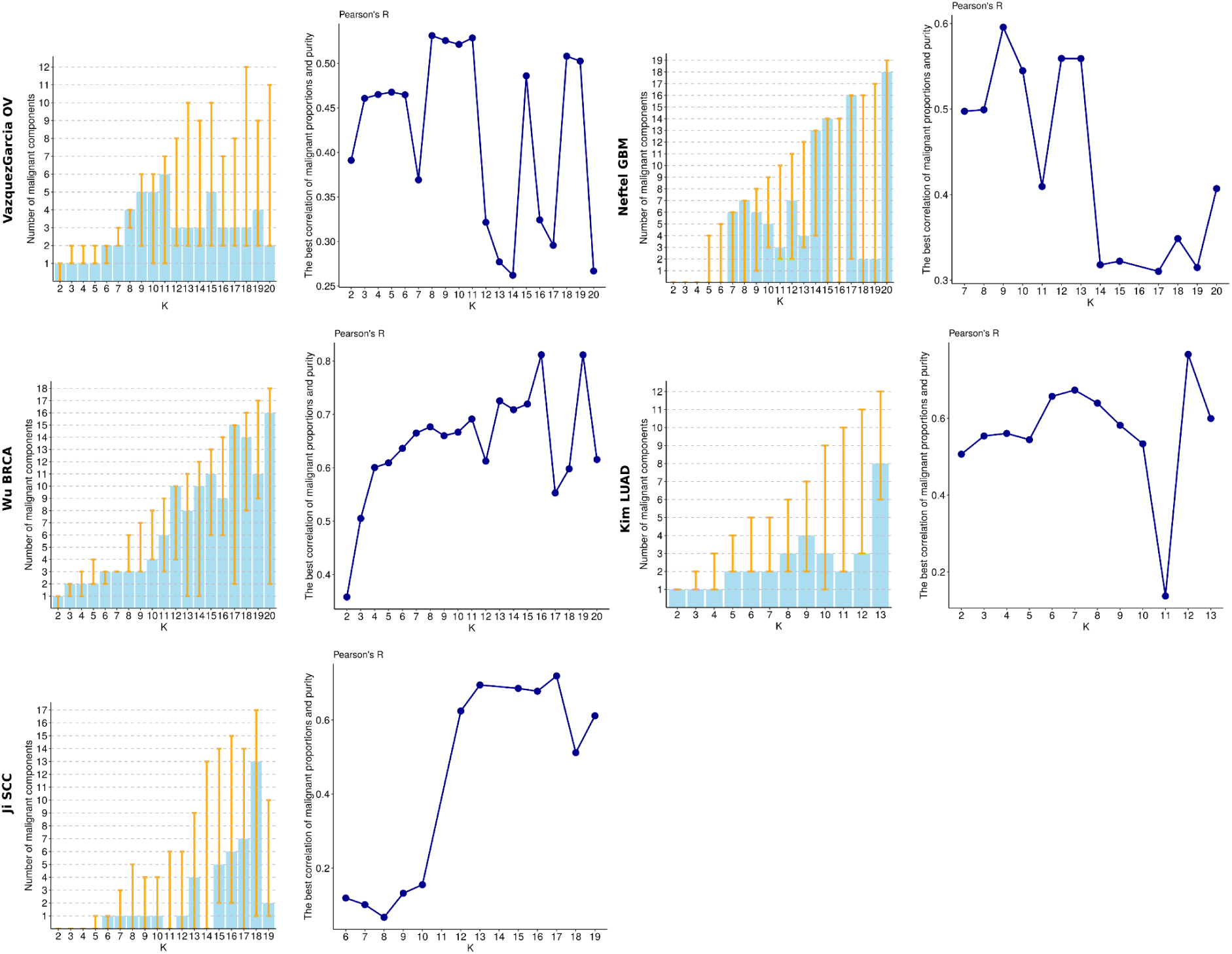
Mode number of malignant cell states, and the correlation between purity with added noise and CDState estimated purity from the best run across those in which the mode malignant count for each *k* was identified.

**Suppl. Figure S34.**
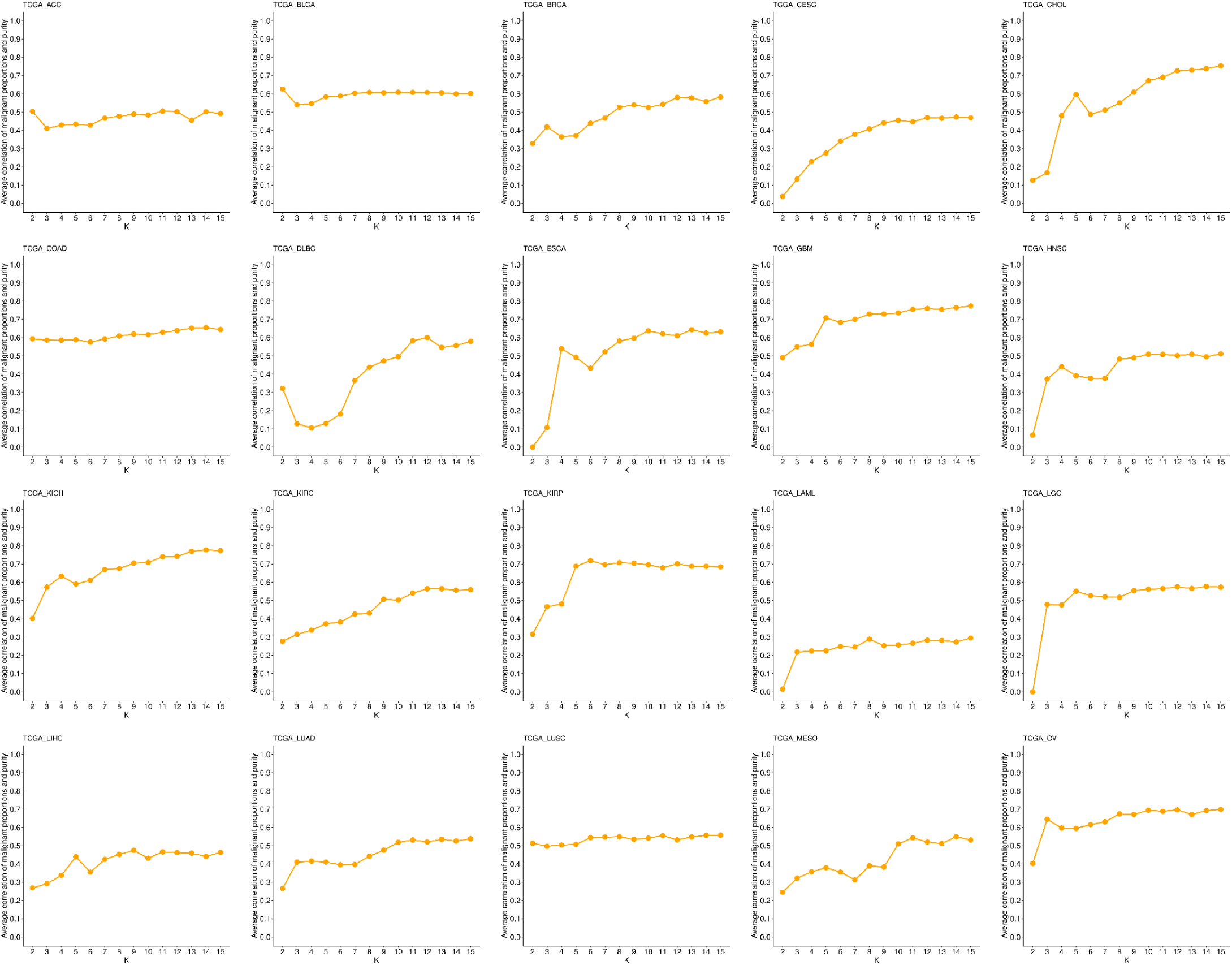
Average Pearson’s correlation across 10 runs and a range of *k* between CDState-inferred and true tumor purity of 20 cancer types from TCGA.

**Suppl. Figure S35.**
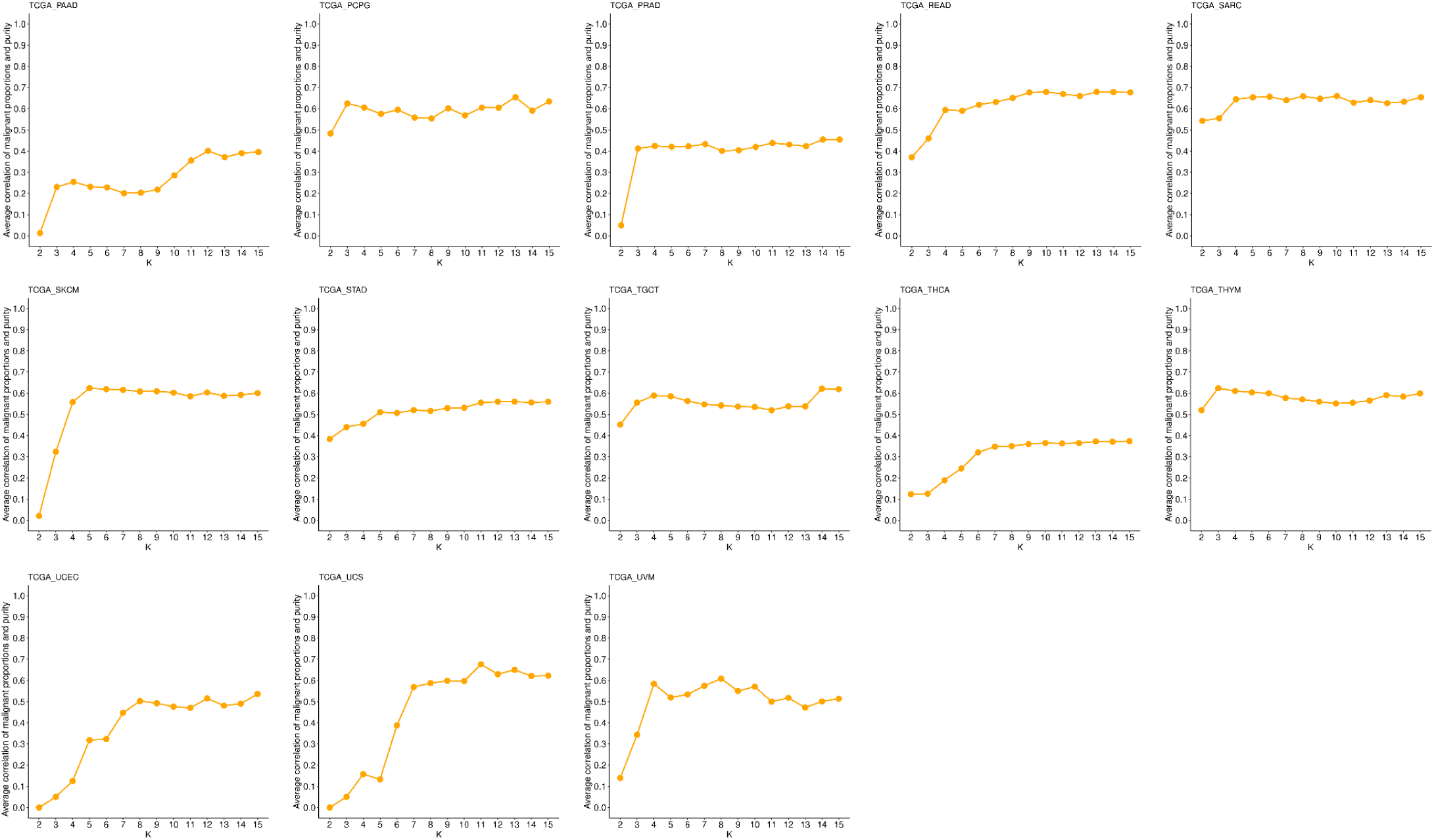
Average Pearson’s correlation across 10 runs and a range of *k* between CDState-inferred and true tumor purity of the remaining 13 cancer types from TCGA.

**Suppl. Figure S36.**
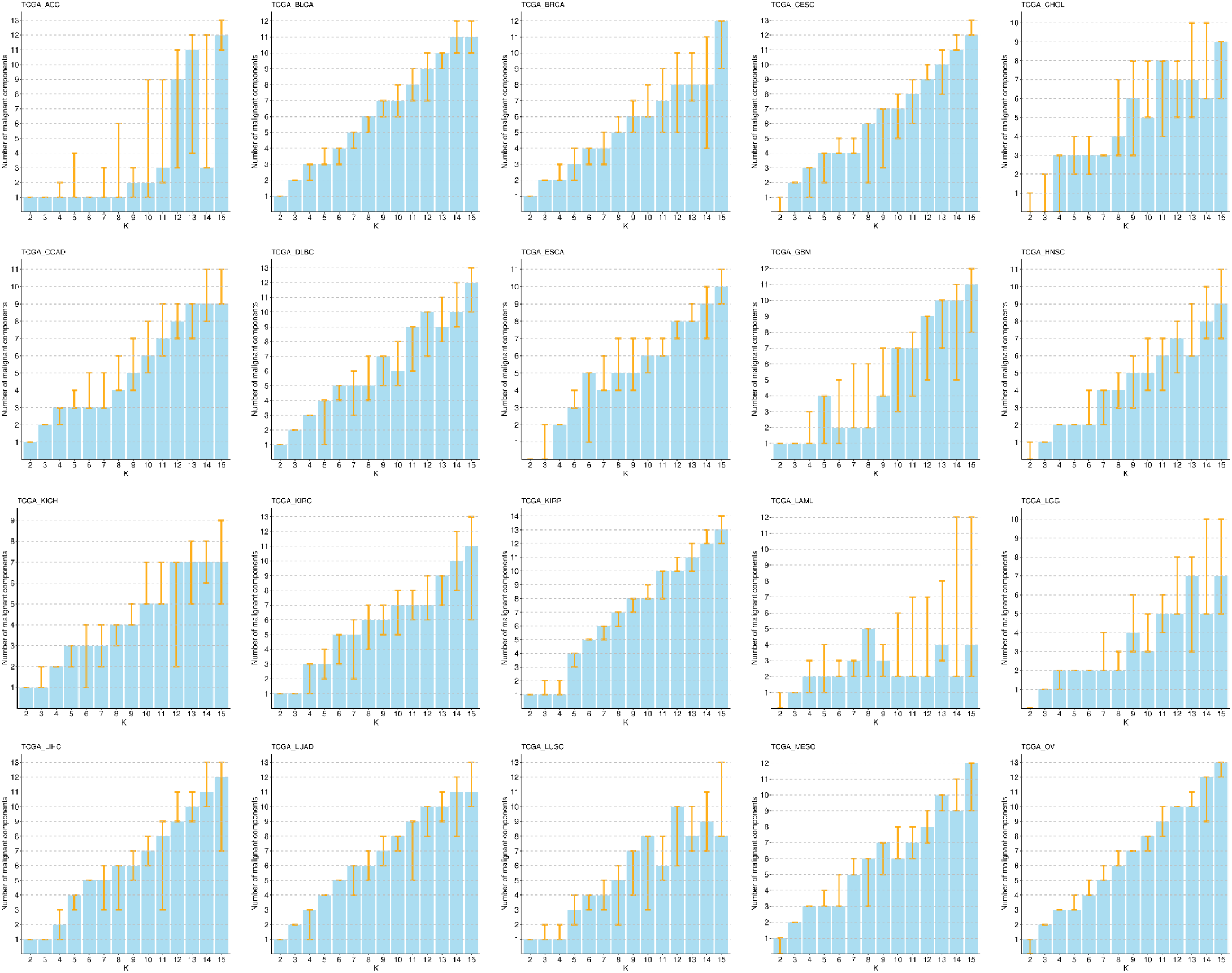
Mode number of malignant cell states identified across 10 runs and a range of *k* of CDState deconvolution of 20 TCGA datasets. Error bars represent the smallest and the highest number of identified malignant cell states for given *k*.

**Suppl. Figure S37.**
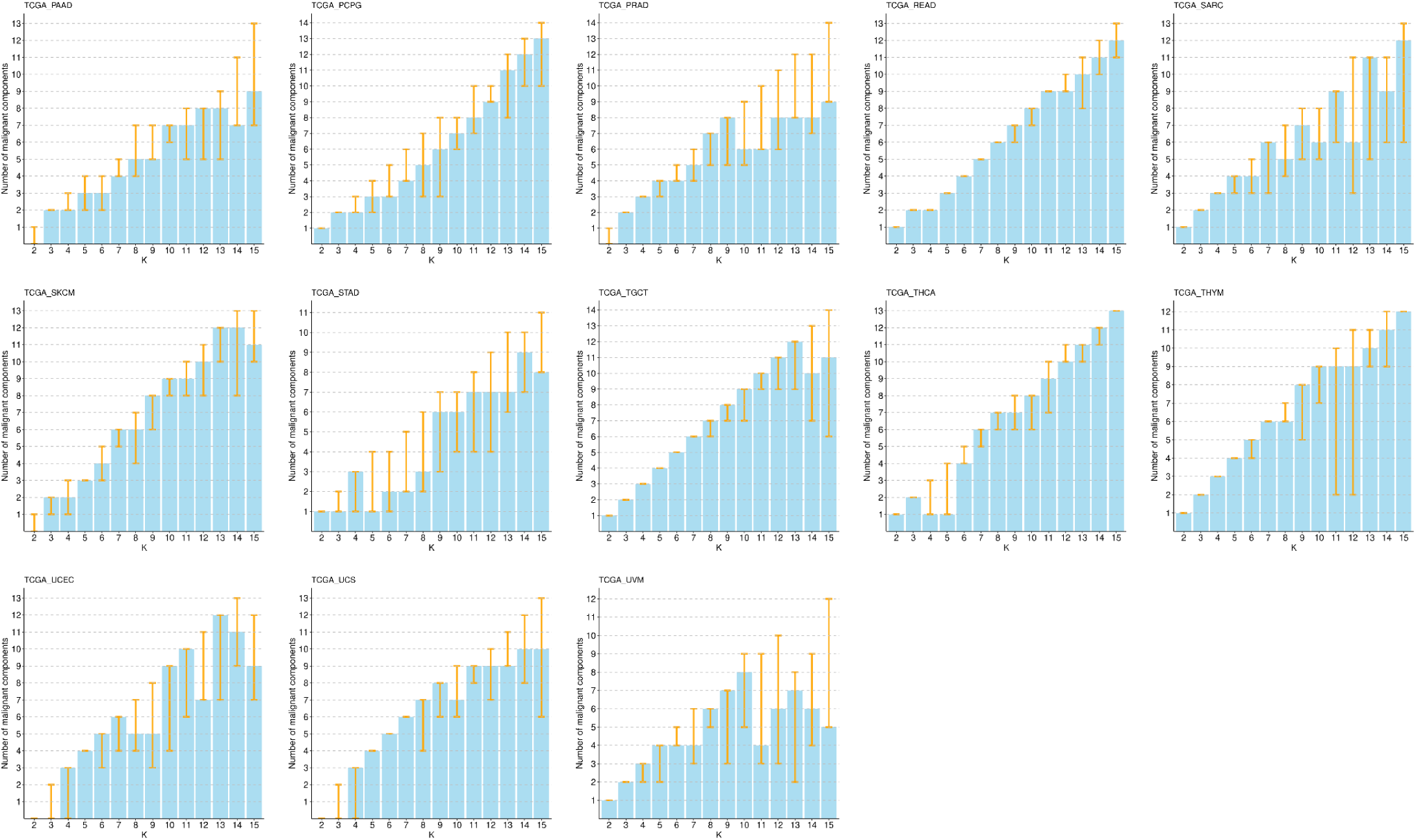
Mode number of malignant cell states identified across 10 runs and a range of *k* of CDState deconvolution of the remaining 13 TCGA datasets. Error bars represent the smallest and the highest number of identified malignant cell states for given *k*.

